# Proximity-induced rewiring of oncogenic kinase triggers apoptosis

**DOI:** 10.1101/2025.06.13.659082

**Authors:** Manuel L. Merz, Veronika M. Shoba, Rajaiah Pergu, Zachary C. Severance, Dhanushka N. P. Munkanatta Godage, Arghya Deb, Hui Si Kwok, Prashant Singh, Sameek Singh, Jonathan B. Allen, Wenzhi Tian, Pallavi M. Gosavi, Santosh K. Chaudhary, Viktoriya Anokhina, Ellen L. Weisberg, N. Connor Payne, Yao He, Rohil Dhaliwal, Reilly Osadchey, Mrinal Shekhar, Ralph Mazitschek, Matthew G. Rees, Jennifer A. Roth, Qiang Cui, James D. Griffin, Brian B. Liau, Amit Choudhary

## Abstract

The active site’s electric field is integral to enzymatic catalysis (e.g., substrate recognition) and nature employs charge-altering post-translational modifications (e.g., phosphorylation) to perturb this electric field and regulate enzymes. A chromosomal translocation converts Abelson kinase (ABL) to BCR-ABL, whose hyperactivity drives several cancers. Here, we developed a small molecule, BRD8833, that induces BCR-ABL phosphorylation, which perturbs its active site’s electric field with loss of hyperactivity. Unlike “occupancy-driven” inhibitors that require stoichiometric concentrations, BRD8833 operates through an event-driven, substoichiometric mechanism by inducing proximity between two BCR-ABL molecules to trigger the inhibitory phosphorylation and selective apoptosis of BCR-ABL-dependent cancer cells. Furthermore, BRD8833 is effective against other oncogenic ABL fusions or clinically observed resistance mutations, including those to occupancy-driven drugs with the same binding site as BRD8833, suggesting differences in their resistance mechanisms. These studies lay the foundation for electric-field and “event-driven” modalities to control hyperactive enzymes with orthogonal resistance mechanisms to occupancy-driven drugs.

**GRAPHICAL ABSTRACT (TOC):** 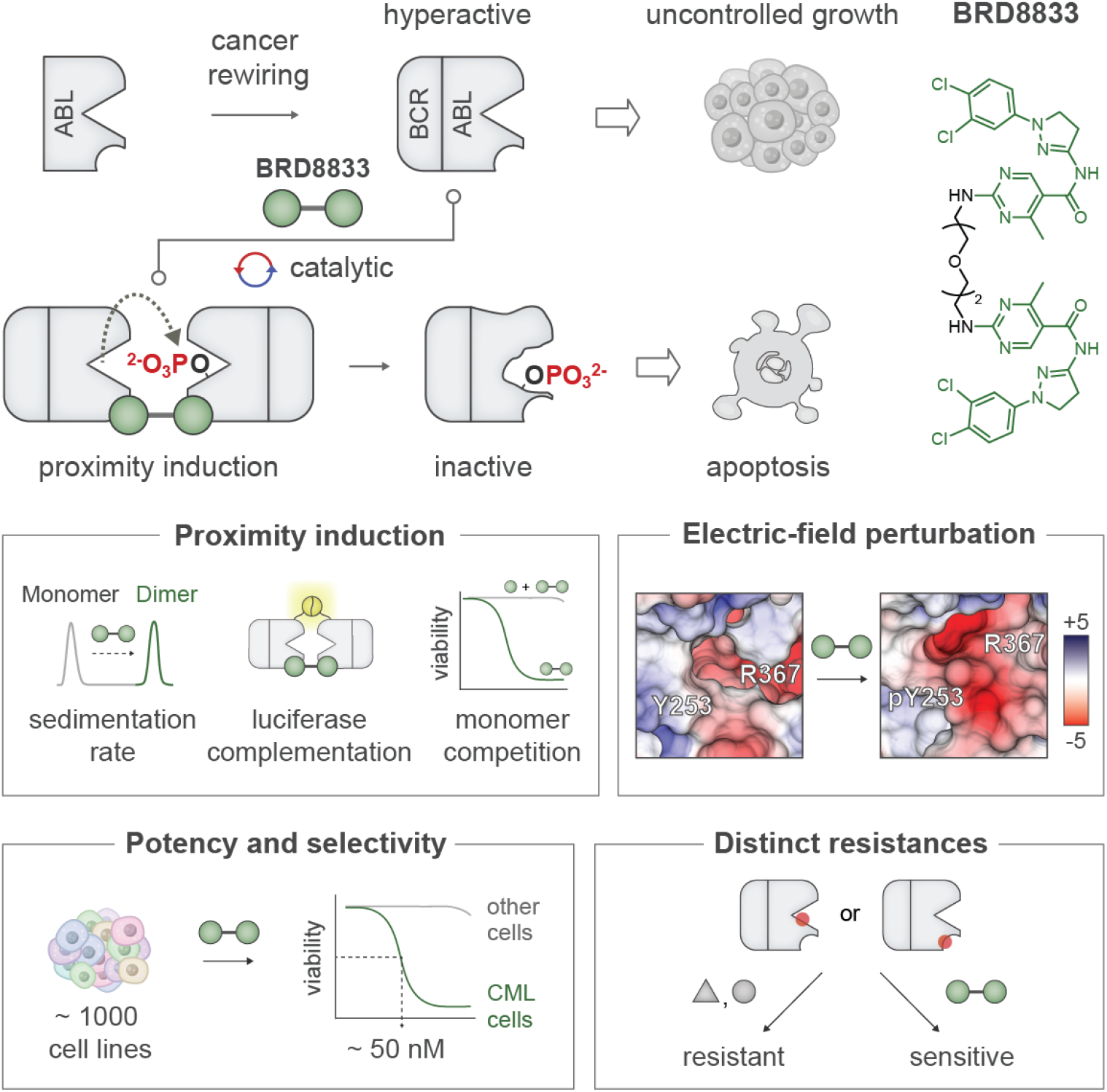

## INTRODUCTION

Enzymes’ active sites have finely-tuned electric fields to enhance catalysis, including for substrate recognition and transition-state stabilization.^1,2^ Perturbing the active site’s electric field by appending or removing charged post-translational modifications (e.g., phosphorylation) can impair enzymatic activity, a regulatory mechanism exploited by nature.^3, 4^ For example, phosphorylation of an active site tyrosine of cyclin-dependent kinase 2 (Cdk2) keeps it in an inactive state, preventing a cell’s progression from G1 to S phase.^5^ Since aberrant enzymatic activity underlies many disorders, including cancer, and resistance has rapidly emerged to classical “occupancy-driven” inhibitors, there is an unmet need to develop fundamentally new mechanisms to inhibit enzymes. Inspired by nature’s use of phosphorylation to regulate enzymes by perturbing their active-site electric field, we ventured to develop small molecules that induce such phosphorylation on a dysregulated enzyme.

We chose BCR-ABL, whose hyperactivity arising from fusion of *BCR* to *ABL* after a chromosomal translocation results in uncontrolled cell growth, triggering chronic myelogenous leukemia (CML) and other myeloproliferative disorders.^6-9^ Like Cdk2, phosphorylation of the active site tyrosine of BCR-ABL is inhibitory and reduces the oncogenicity of CML. We envisioned that a chemical inducer of proximity (CIP) that dimerizes BCR-ABL and utilizes its hyperactivity to induce inhibitory phosphorylation onto itself would have several attractive attributes. First, these CIPs will operate by an “event-driven mechanism”^10-13^ vs. the “occupancy-driven mechanism” of active site or allosteric inhibitors that require sustained BCR-ABL occupancy for the pharmacological effect. CIP-induced inhibitory phosphorylation will persist even after dissociation from BCR-ABL, enabling turnover and sub-stoichiometric potency.^14^ Second, CIP will have different resistance mechanisms from occupancy-driven inhibitors, for which resistance has rapidly emerged.^15-18^ Third, the requirement of both binding and phosphorylation induction events provides an additional layer of selectivity that may reduce off-targets.^19^ Finally, since active sites are rich in phosphorylable amino acids and CIP can be developed for nearly all kinases utilizing their abundantly available kinase inhibitors,^10^ this approach is potentially generalizable to other oncogenic kinases and enzymes.

Here, we report the development of BRD8833 (generated by dimerizing ABL binders) that forms a ternary complex with BCR-ABL and induces inhibitory phosphorylation at Y253 in the active site’s P-loop, equivalent to the regulatory site on Cdk2.^20^ Molecular dynamics simulation confirms that this phosphorylation reconfigures the electrostatic arrangement of residues in the active site. BRD8833 potently induces apoptosis of BCR-ABL-dependent cancer cells at a low nanomolar concentration, which is ∼20-fold lower than the concentration required for complete BCR-ABL occupancy by BRD8833, suggesting a sub-stoichiometric mechanism. This potency was specific, as BRD8833 showed selectivity for BCR-ABL-dependent cancer cells in a pooled screening of ∼1000 cancer cells from diverse lineages and genotypes. BRD8833 was also active in cells harboring a gatekeeper mutation (T315I), which renders active-site drugs (e.g., imatinib) ineffective, and in cells with other oncogenic ABL fusions (e.g., TEL-ABL) that are insensitive to approved drugs (e.g., asciminib).^8, 21, 22^ CRISPR-based mutagenesis studies showed that BRD8833, which binds to the same site as the allosteric inhibitor asciminib, was effective in asciminib-resistant cells and exhibited a resistance profile distinct from that of asciminib. Since > 40% of the human kinases and most GTPases contain P-loops with S/T/Y residues that can be phosphorylated, we reason that this inhibitory mechanism may be applied beyond BCR-ABL to other enzymes. Overall, BRD8833 introduces a fundamentally new modality that operates by electric-field effects and event-driven mechanism to target an oncogenic enzyme with a different resistance mode than occupancy-based drugs.

## RESULTS

### BRD8833 potently inhibits the growth of BCR-ABL-dependent K562 cells

We hypothesized that a CIP consisting of a homodimer of ABL binders might inhibit BCR-ABL by inducing inhibitory phosphorylation (**Figure 1a**). Towards that goal, we synthesized homodimer **2** by connecting ABL binder **1** from a solvent-exposed site^23^ (**Figure S1a-c**) and used mass spectrometry to assess alteration in phosphorylation levels of ABL in the presence of **2**. We were gratified to see an increase in levels of Y253 phosphorylation of ABL in the presence of **2**, while such phosphorylation was undetectable in DMSO or monomer **1** treated ABL (**Figure S1d**). Previous studies^24^ have shown that phosphorylation at Y253, which is part of the ATP-binding loop (P-loop) and proximal to the DFG motif, reduces BCR-ABL oncogenicity (**Figure 1b**). In agreement with these studies, **2** reduced the viability of BCR-ABL dependent K562 cells with an EC_50_ of 740 nM (**Figure S1e**).

**Figure 1.**
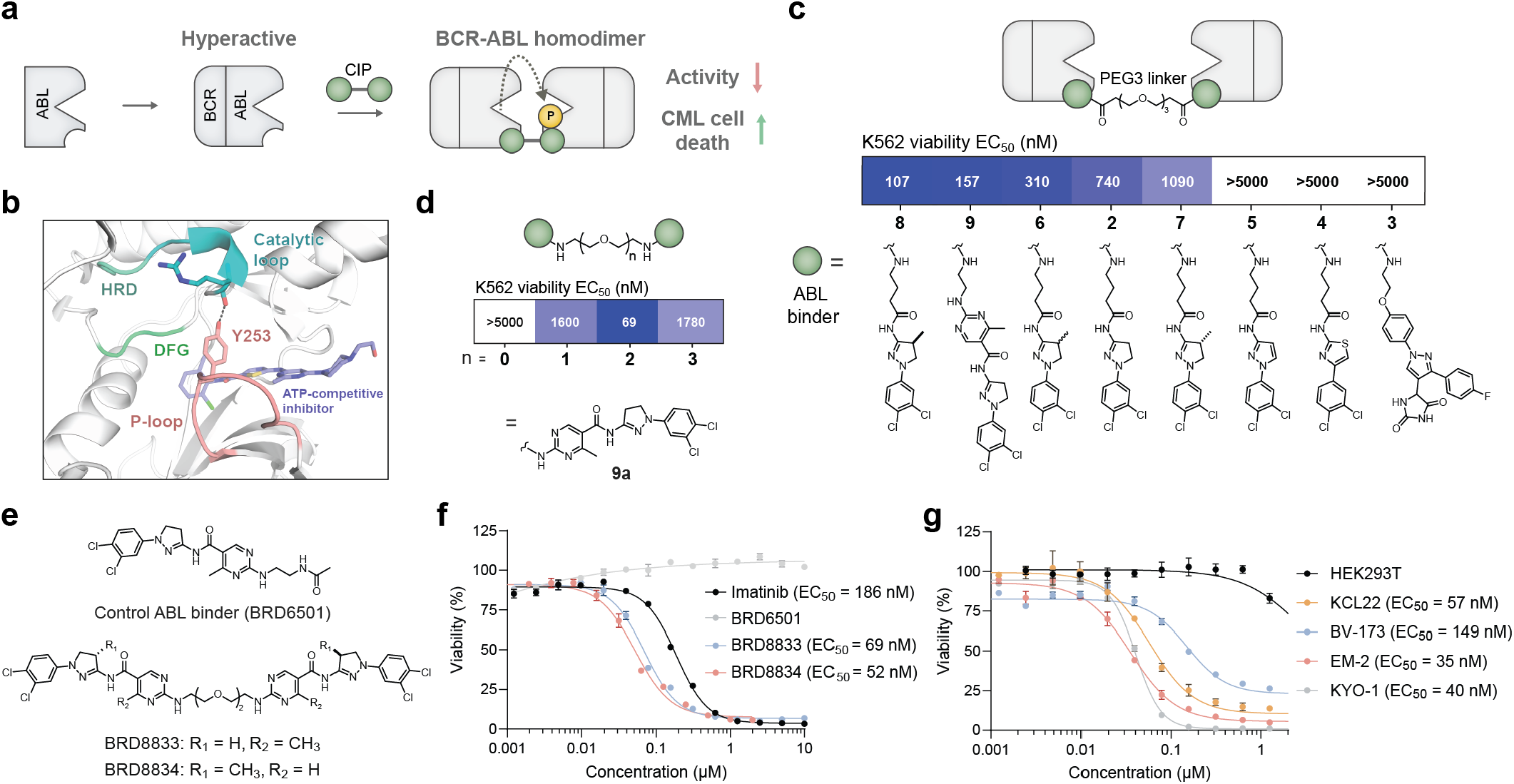
Development of a potent inhibitor of BCR-ABL oncogenesis. **a** Dimerization of BCR-ABL with CIPs catalyzes inhibitory autophosphorylation. **b** Co-crystal structure of ABL and ATP competitive inhibitor dasatinib (PDB 2GQG). DFG motif, catalytic loop with HRD motif, and hydrogen bonding of Y253 (black dotted line) are shown. P-loop residues 248–255 are highlighted. **c** Structure-activity relationship of BCR-ABL CIPs. PEG3 linker was used to join different ABL activators, and cytotoxicity in K562 cells expressing BCR-ABL was measured. **d** Different PEG linkers were used to join two ABL ligands **9a**, and cytotoxicity in K562 cells was measured. **e** Structures of optimized CIPs (BRD8833 and BRD8834) and respective ABL binder control (BRD6501). **f** Dose response for cytotoxicity in K562 for different BCR-ABL inhibitors (BRD8833, BRD8834, imatinib) and the control ABL binder BRD6501. **g** Cytotoxicity of BRD8833 in HEK293T and different cell lines expressing the oncogenic BCR-ABL fusion. EC_50_ values in **c** and **d** were calculated from three independent replicates. **f** and **g** show the mean and SD of three independent replicates.

Using the viability of K562 cells as a readout, we performed structure-activity relationship studies by systematically varying the linker between the previously reported ABL binders (**Figure 1c and Figure S2a**).

Compounds **3** (hydantoin scaffold),^25^ **4** (thiazole scaffold) and **5** (pyrazole scaffold) exhibited loss of activity (EC_50_ > 5000 nM). Compound **6**, which is assembled from a methylated analog derived from the dihydropyrazole scaffold of **2**, exhibited increased K562 cytotoxicity (EC_50_ = 310 nM) in agreement with prior studies^23^ that methylation of the dihydropyrazole scaffold increases ABL activation by 1.3 fold. Furthermore, the *S*-enantiomer (**8**) of compound **6** is approximately 10-fold more potent than the (*R*) enantiomer (**7**) (EC_50_ = 107 nM and 1090 nM, respectively), in agreement with their reported degree of ABL activation. Finally, linking a methyl pyrimidine on the amino dihydropyrazole through an amide bond afforded compound **9**, which displayed comparable activity to **8** (EC_50_ = 157 nM). Using a methyl pyrimidine scaffold **9a** for synthetic simplicity and avoiding the chiral centers in **8**, we varied the length of PEG linkers (n = 0–3) between the two ABL binders and identified PEG2 (BRD8833) as optimal, achieving an EC_50_ of 69 nM (**Figure 1d,e and Figure S2b**). Substituting the methyl group from pyrimidine to (*S*)-4-methyl dihydropyrazole further improved potency slightly (BRD8834, EC_50_ = 52 nM). However, due to minimal gain and challenges with chiral purification, we proceeded with BRD8833 as the optimized molecule. Notably, BRD8833 exhibited higher potency than the first-in-class BCR-ABL drug imatinib in reducing the viability of K562 cells (**Figure 1f**). Next, we confirmed the activity of BRD8833 in other BCR-ABL positive CML cell lines (KCL22, BV-173, EM-2 and KYO-1). We observed dose-dependent cytotoxicity (EC_50_: 35–149 nM), but not on HEK293T cells that lack BCR-ABL (EC_50_: > 3000 nM, **Figure 1g**). The observed dependence of compound activity on linker length, the potency of ABL activation by the binder scaffold, and BCR-ABL dependence points to an induced-proximity mechanism, which we investigated next.

### BRD8833 promotes ternary-complex formation and induces Y253 phosphorylation on BCR-ABL

We first assessed the binding to ABL using differential scanning fluorimetry (DSF). BRD8833 and the monomer ABL binder control BRD6501 increased the protein melting temperature (T_m_), indicative of ABL-binding (**Figure S3a– c**). We established a TR-FRET assay to monitor binding to the myristoyl allosteric pocket (**Figure S3d**,**e**). Here, the TR-FRET signal is observed when the tracer (derived from an allosteric myristoyl pocket binder) is in proximity to a CoraFluor-labeled anti-GST nanobody donor—thus, signal decrease upon competitive displacement of the tracer by BRD8833 measures target engagement.^26^ As expected, BRD8833, ABL binder BRD6501 (**Figure 2a**), and asciminib (**Figure S3f**) reduced the TR-FRET signal dose-dependently, consistent with their overlapping binding pocket. Next, we used analytical ultracentrifugation to confirm ternary complex formation by BRD8833. BRD8833, but not vehicle or the control ABL binder BRD6501, caused a shift in the sedimentation coefficient, consistent with ABL dimerization (**Figure 2b**).

**Figure 2.**
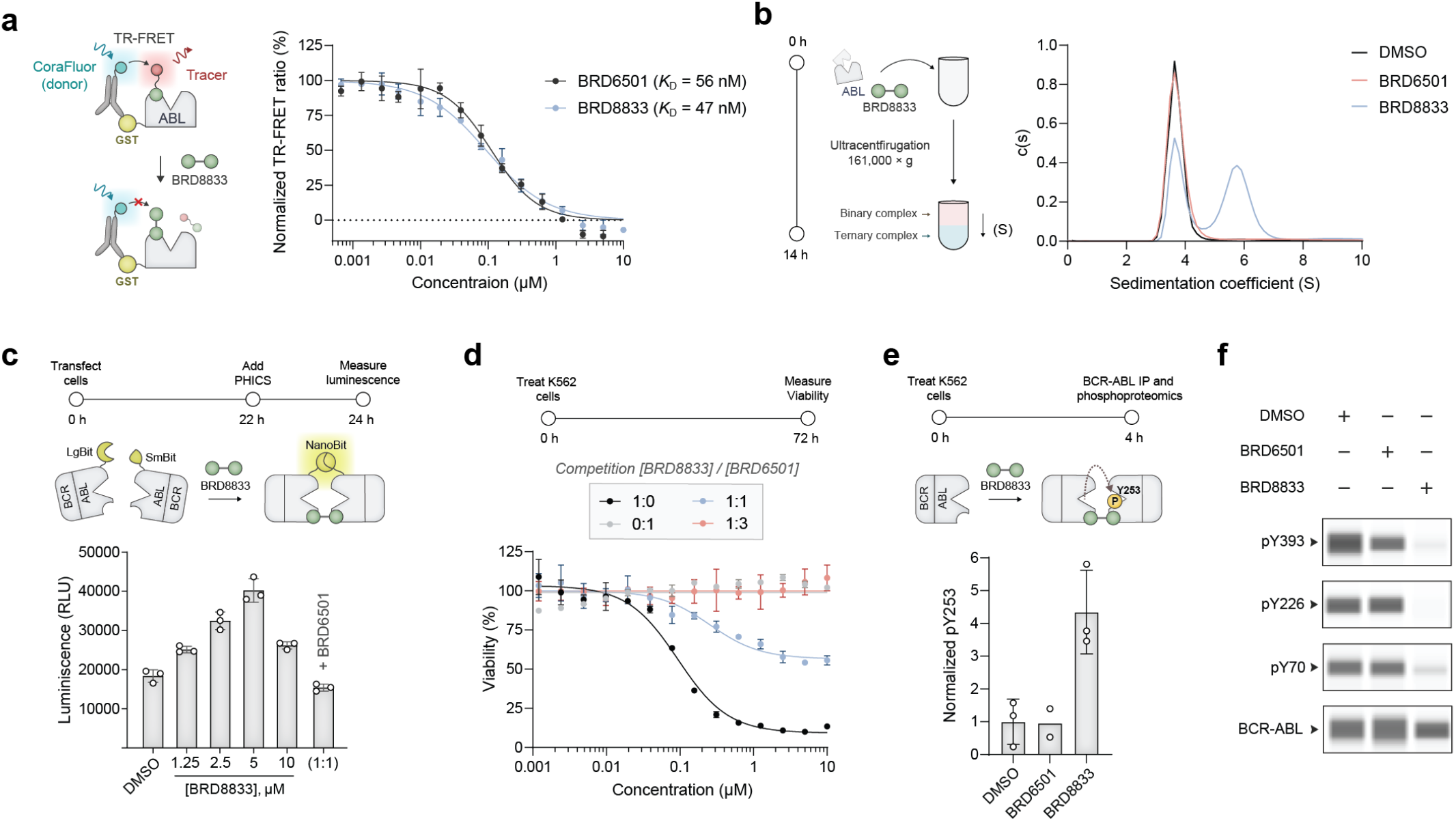
Assessment of ternary complex formation and phosphorylation induction by BRD8833. **a** TR-FRET assay to confirm binding to the allosteric myristoyl pocket of the ABL kinase. Dose-dependent displacement of the tracer by BRD8833 and control BRD6501 was measured. Mean and SD of three independent replicates are shown. **b** Analytical ultracentrifugation to measure the sedimentation coefficient and biochemical ternary complex formation of the ABL kinase in the presence of either BRD8833 or its control BRD6501 (1.5 µM). **c** Dose-dependent cellular ternary complex formation by BRD8833 in HEK293T cells transfected with LgBiT and SmBiT BCR-ABL. For competition, a 1:1 ratio of BRD8833 to BRD6501 was added (5 µM each). Mean and SD for three independent replicates are shown. **d** Rescue of dose-dependent cytotoxicity of BRD8833 in K562 cells. Mean and SD of three independent replicates are shown. **e** Phosphorylation analysis by LC-MS/MS. Relative phosphorylation levels to unmodified peptides were measured after treatment with either BRD8833 or its control BRD6501. Vehicle (DMSO) and BRD8833-treated samples were measured in three independent replicates, while BRD8833 was measured in duplicates. Mean and SD (for BRD8833 and DMSO) are shown. **f** Immunoblotting to probe autophosphorylation on BCR-ABL after treatment of K562 cells with of either BRD8833 (1 µM) or BRD6501 (2 µM). A representative example for three independent measurements is shown.

To further confirm ternary complex formation in cells, we used a split luciferase NanoBiT assay, where BCR-ABL constructs were fused to luciferase components LgBit and SmBit (**Figure 2c**). Treatment with BRD8833 increased luminescence, indicating ternary complex formation, while competition with BRD6501 reduced the signal. Indicative of an induced-proximity mechanism, we observed a characteristic “hook effect”^27^ by increasing the concentration of BRD8833. In agreement with these findings, we found that ABL binder BRD6501, which is not cytotoxic to K562 cells, rescued the killing activity of BRD8833 when K562 cells were co-treated with BRD6501 and BRD8833 (**Figure 2d**). To determine alteration in BCR-ABL phosphorylation, we performed mass spectrometry on immunoprecipitated BCR-ABL from K562 cells treated with DMSO, BRD6501, or BRD8833 (**Figure 2e**). While treatment with BRD6501 had similar phosphorylation levels at Y253 to the vehicle (DMSO), BRD8833 selectively increased phosphorylation at Y253. Phosphorylation at Y253 correlates with reduced BCR-ABL activity^24^ and in agreement with these studies, we observed reduced autophosphorylation at multiple activation sites on BCR-ABL (pY70, pY226, and pY393) in BRD8833-treated K562 cells using immunoblotting (**Figure 2f**).^6, 8, 28^ We further performed a global phosphoproteomic analysis and confirmed increased Y253 phosphorylation and decreased phosphorylation at sites correlating with autoactivation (pY226 and pY393). In contrast, the global proteome was comparably unaltered relative to DMSO treatment (**Figure S3g–i**). These findings suggest that BRD8833 forms a ternary complex with a BCR-ABL homodimer to selectively promote Y253 phosphorylation.

### Molecular dynamics (MD) simulation suggests Y253 phosphorylation perturbs active-site’s electric field

In the apo state, phosphorylation of Y253 (Y253TP2) is observed to induce a significant conformational change in the P-loop, leading to the formation of a salt bridge between the phosphate group with R367 that obstructs ATP binding (**Figure 3a and Figure S4a**,**b**). Electrostatic potential calculations further reveal that Y253 phosphorylation enhances the negative potential of the active site, disfavoring the binding of ATP’s triphosphate moiety (**Figure 3b**,**c**). Simulations of activated BCR-ABL in complex with ATP·Mg^2+^ also suggest that phosphorylation of Y253 impedes the catalytic competency. Intuitively, one might expect that the negatively charged Y253TP2 and ATP would strongly repel each other. Instead, the MD simulations suggest that the high negative charge density draws sodium ions into the binding pocket, forming a stable ionic cluster (compare **Figure 3d,e**) between Y253TP2’s phosphate group, ATP’s triphosphate moiety, R367, and three to four sodium cations (**Figure 3e,f and Figure S4c**,**d**). The formation of this tightly packed cluster causes inversion of the P-loop and prevents access to ATP’s γ-phosphate, thereby precluding substrate phosphorylation.

**Figure 3.**
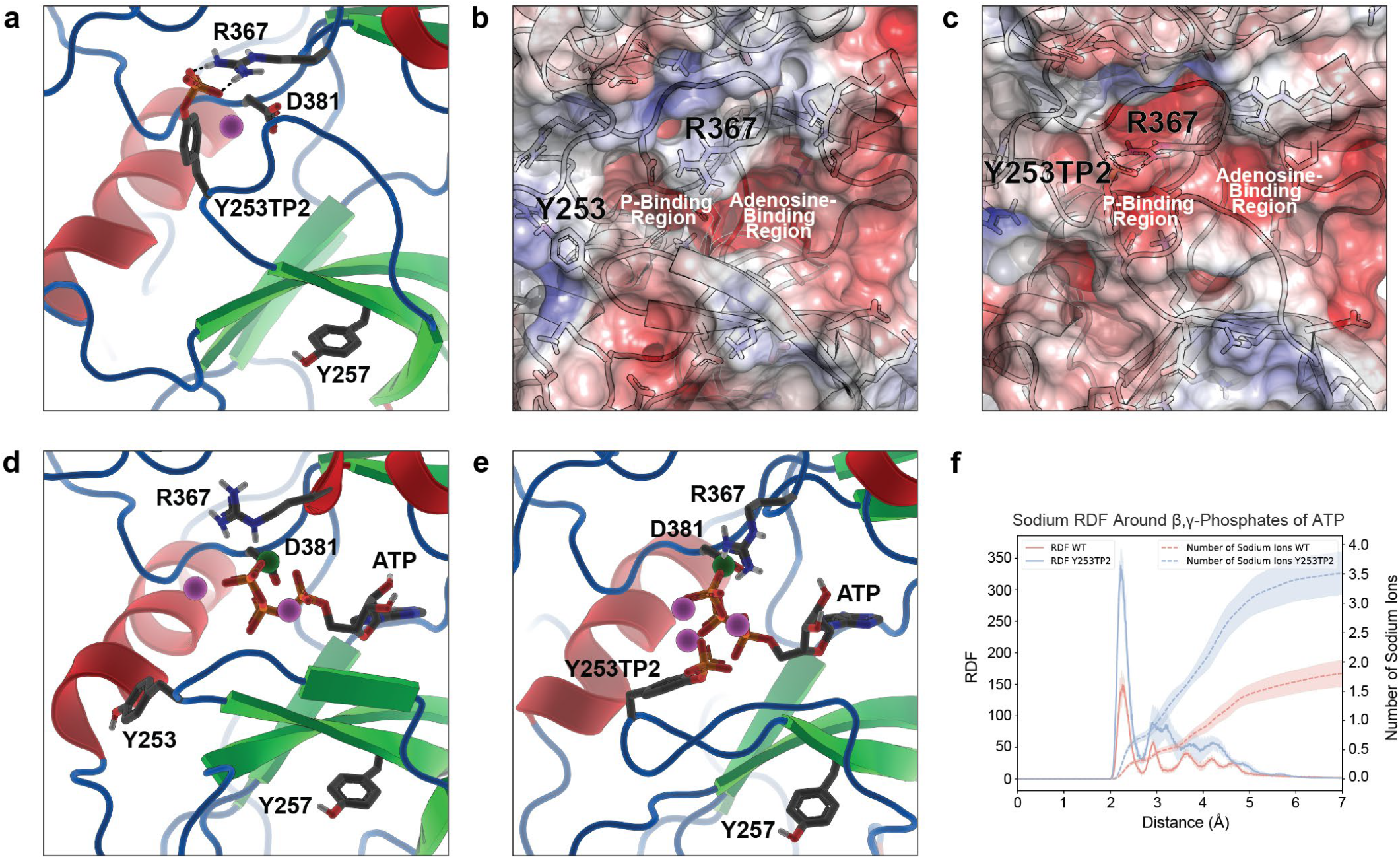
MD simulations suggest an inhibitory mechanism of Y253 phosphorylation. **a** Phosphorylation of Y253 (Y253TP2) in the apo state results in salt bridge formation with R367, blocking ATP binding. A nearby Na^+^ ion (violet sphere) neutralizes the ion pair’s net charge. **b–c** Electrostatic potential of BCR-ABL binding pocket in the (**b**) wild-type and (**c**) Y253TP2 apo states. Scale ranges from -5 (red) to +5 (blue) *k*_B_*T*/*e*, where *k*_B_ is Boltzmann’s constant, *e* is the elementary charge, and *T* = 303.15 K. **d–e** Snapshots of BCR-ABL complexed to ATP · Mg^2+^ in (**d**) wild-type and (**e**) Y253TP2 states illustrate the formation of an ionic cluster formed by the phosphorylated Y253, the *β,γ*-phosphates of ATP, R367 and several Na^+^ ions that impedes the kinase function. Na^+^ and Mg^2+^ ions are shown as violet and dark green spheres, respectively. **f** Radial distribution functions of Na^+^ ions around the *β,γ*-phosphates of ATP in the wild-type (red) vs. Y253TP2 (blue) states demonstrate increased concentration of Na^+^ ions within the active site following Y253 phosphorylation. The dashed lines indicate the integrated number of Na^+^ ions up to a distance to the center of *β,γ*-phosphates of ATP. The shaded area indicates statistical uncertainty.

To investigate whether this mechanism could be applied more broadly to other kinases, we searched the human kinome for residues that can be phosphorylated on their P loop. Nearly 56.1% of human kinases have a canonical “GXGXXG” P-loop motif, and an additional 12.3% have a related near P-loop motif “GXGXX[A/S]” (**Figure S5a**,**b**). 63.3% of those motifs harbor at least one S/T/Y that can get phosphorylated, and 22.7% even contain the conserved Y that corresponds to Y253 in the BCR-ABL P-loop. Similarly, GTPases also contain a P-loop motif that features a conserved S/T residue that coordinates to the nucleotide phosphates.^29^ Many of the identified kinases and GTPases (e.g., KRAS) are therapeutically relevant (**Figure S5c**), pointing to phosphorylation-induced perturbation of the active-site electric field as a potential regulatory mechanism for disease-associated enzymes.

### BRD8833 inhibits BCR-ABL signaling

We next sought to investigate the effects of BRD8833 on BCR-ABL signaling. Treatment of K562 cells with BRD8833 (1 µM) inhibited substrate phosphorylation and downstream signaling pathways of BCR-ABL, including pSTAT5, pERK, and pCRKL,^6, 8, 30^ while the control ABL binder BRD6501 (2µM) had no effect (**Figure 4a**). We observed cleavage of both caspase 8 and PARP, consistent with induction of apoptosis by BRD8833 upon inhibition of BCR-ABL signaling (**Figure 4b**). Collectively, these findings demonstrate that BRD8833 reduces the viability of BCR-ABL-dependent cell lines by selectively inducing apoptosis and suppressing the oncogenic BCR-ABL signaling. Finally, we compared the transcriptomic effects of BRD8833, the allosteric inhibitor asciminib, and the ABL binder BRD6501 using RNA sequencing (RNA-seq). K562 cells were treated with DMSO (vehicle), asciminib (100 nM), BRD8833 (1 µM), or BRD6501 (2 µM) for 6, 12, or 24 hours in biological quadruplicates. Gene expression levels were analyzed using replicate PCA plots, gene set enrichment analysis (GSEA), and moderated t-tests to compare treatment groups (**Figure 4c,d and Figure S6**). We observed nearly identical gene expression profiles for cells treated with BRD8833 and asciminib (Group A in **Figure 4d**). Similarly, the gene expression profiles for the vehicle control and BRD6501 were almost indistinguishable, confirming that the ABL binder BRD6501 is not cytotoxic and behaves similarly to the vehicle. Remarkably, these findings suggest that converting the allosteric binder for ABL into a CIP transforms a ligand with no detectable effects, comparable to the vehicle control, into a potent and selectively cytotoxic compound that induces gene expression changes closely matching those of a clinically approved drug.

**Figure 4.**
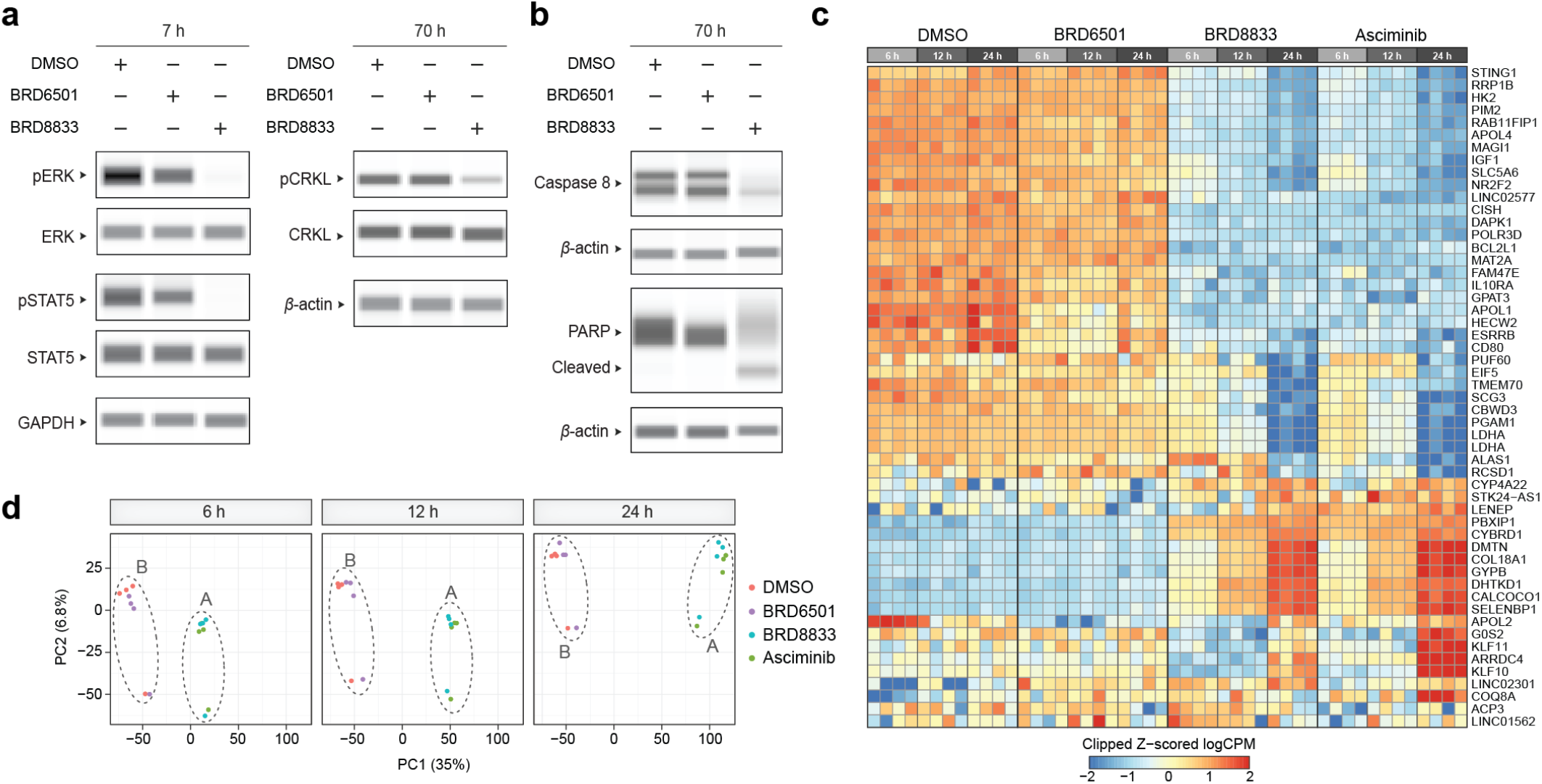
BRD8833 disrupts BCR-ABL signaling pathway. **a** Inhibition of BCR-ABL downstream signaling. K562 cells were treated with either BRD8833 (1 µM) or its control BRD6501 (2 µM), and pSTAT5, pERK, and pCKRL were measured relative to the respective total protein level. **b** Activation of pro-apoptotic pathways. Immunoblotting after 70-hour incubation with either BRD8833 (1 µM) or its control BRD6501 (2 µM) is shown. Immunoblotting experiments in **a** and **b** were measured in at least three independent replicates, and a single representative example is shown. **c** Global transcriptomic analysis in K562 cells treated with DMSO, BRD6501, BRD8833, or asciminib. Equal number of genes scoring the highest p-values for each comparison are shown. **d** PCA plot for gene expression analysis. Cluster A is formed by BRD8833 and asciminib, while cluster B is formed by DMSO and BRD6501. Gene expression was analyzed across four independent biological replicates.

### BRD8833 selectively inhibits BCR-ABL and other oncogenic ABL fusions

Beyond BCR-ABL, several other oncogenic ABL fusions are known.^8^ These different fusion partners can change the localization, catalytic efficiency, sensitivity to inhibitors, and substrate preferences of the kinase.^31, 32^ For example, the TEL-ABL fusion has much higher *in vitro* and *in vivo* activity than BCR-ABL. To thus test BRD8833 against these alternative fusion proteins, we treated Ba/F3 cells expressing TEL-ABL with BRD8833, which reduced the viability with an EC_50_ value of 280 nM (**Figure 5a**). In contrast, the clinically approved allosteric ABL-inhibitor, asciminib,^33, 34^ which binds to the same myristoyl pocket as BRD8833, did not affect the cell viability. However, the low efficacy of asciminib is not a result of reduced binding affinity, as the addition of one equivalent of asciminib reversed the effect of BRD8833 (**Figure 5a**). This not only demonstrates that BRD8833 potently inhibits the viability of different oncogenic ABL-fusions, but also highlights that they exhibit a unique mode of action that differs from classical occupancy-driven inhibitors such as asciminib.

**Figure 5.**
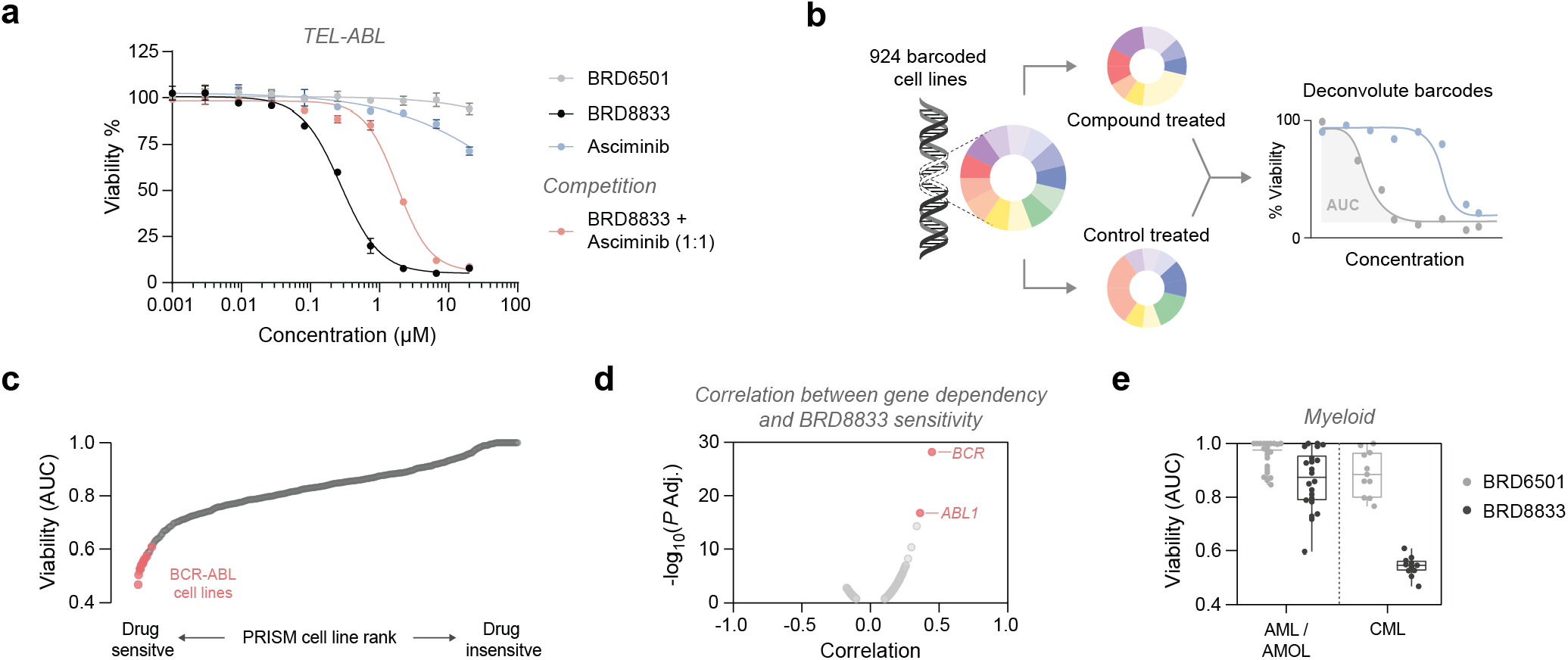
BRD8833 is effective against oncogenic ABL fusions. **a** Cytotoxicity in Ba/F3 cells expressing TEL-ABL. Mean and SD of three independent replicates are shown. **b** Schematic of the PRISM screening. Barcoded cell lines were incubated for 5 days with either BRD8833, its control BRD6501, or imatinib in an 8-point and 3-fold serial dilution. **c** Ranking of PRISM cell lines according to sensitivity for BRD8833. Cell lines annotated BCR-ABL positive by the DepMap portal are highlighted. **d** Correlation between CRISPR knockout gene dependency and BRD8833 sensitivity, calculated at a concentration of 740 nM. **e** Boxplot showing lineage-specific sensitivity to BRD8833 and its control BRD6501. The box extends from the first to third quartiles, the median is depicted as center line, and the whiskers depict the range. PRISM screen was performed in three independent replicates (n = 3).

To evaluate the selectivity of BRD8833 and gain a comprehensive understanding of its activity in CML and other contexts, we conducted PRISM multiplexed screening on 924 barcoded cancer cell lines originating from various tissues with diverse genetic backgrounds (**Figure 5b**).^35^ Crucially, the DepMap portal annotates cell lineages and oncogenic features, enabling targeted analysis of differential responses between cell types, such as those expressing oncogenic ABL fusions.^36^ We treated cell pools with either BRD8833 or the ABL binder control BRD6501 in a 5-day viability assay, using an 8-point dose range (3-fold dilution). Among the tested cell lines, eleven were annotated as BCR-ABL positive and exhibited increased sensitivity to BRD8833, ranking among the top 33 most drug-responsive cell lines (top 3.6% of the 924 tested; **Figure 5c**). We further correlated drug-response with CRISPR gene-dependencies and observed a requirement of *BCR* and *ABL1* for BRD8833 efficacy (**Figure 5d**). In contrast, the ABL binder BRD6501 showed no considerable inhibitory effect on the BCR-ABL positive cell lines (**Figure S7a**). A side-by-side analysis of BRD8833 and imatinib revealed comparable sensitivity profiles, converging on the targeting of cell lines expressing BCR-ABL fusions (**Figure S7b**). Notably, the previously studied cell lines, including K562, KCL22, BV-173, EM-2, and KYO-1, also displayed increased sensitivity to both BRD8833 and imatinib in the PRISM screen (AUC_viability_ < 0.6). Further analysis by lineage revealed that myeloid cell lines, specifically those derived from chronic myeloid leukemia (CML), were more sensitive to BRD8833, while acute myeloid leukemia (AML) cell lines and other lineages showed no clear dependencies (**Figure 5e and Figure S7c**). These findings indicate that BRD8833 exhibits high selectivity for CML cells dependent on ABL oncogenic fusions, with minimal off-target effects on other cell lines.

### BRD8833 displays an orthogonal resistance mechanism to that of asciminib

Resistance development is a common failure mode of many BCR-ABL targeting inhibitors.^6, 37^ Gratifyingly, BRD8833 remained effective against clinically relevant imatinib-resistant E255V and gatekeeper mutant T315I in Ba/F3 cells (**Figure S8a**,**b**). Since asciminib and BRD8833 bind to the same allosteric pocket but with different inhibition mechanisms, we hypothesized that their resistance mechanisms would differ. To identify resistance mutation hotspots, we performed CRISPR-suppressor scanning.^38, 39^ Here, we used the mutagen SpCas9 and a library of guide RNAs (sgRNA) tiling across the coding region of ABL1 to generate a pool of BCR-ABL variants in K562 cells (**Figure 6a**). We applied dose escalation of BRD8833 or asciminib as a selection pressure over 63 days, increasing the dose at days 14 and 21, which ultimately enriched for cells carrying resistance mutations (**Figure S9a**). We treated enriched cells with either asciminib or BRD8833 (**Figure 6b and Figure S9b**) and observed a 1000-fold and 3-fold loss in sensitivity for asciminib and BRD8833, respectively. Interestingly, we identified an orthogonal resistance for BRD8833 and asciminib (**Figure 6c**), where cells subjected to asciminib remained sensitive to BRD8833 and vice versa.

**Figure 6.**
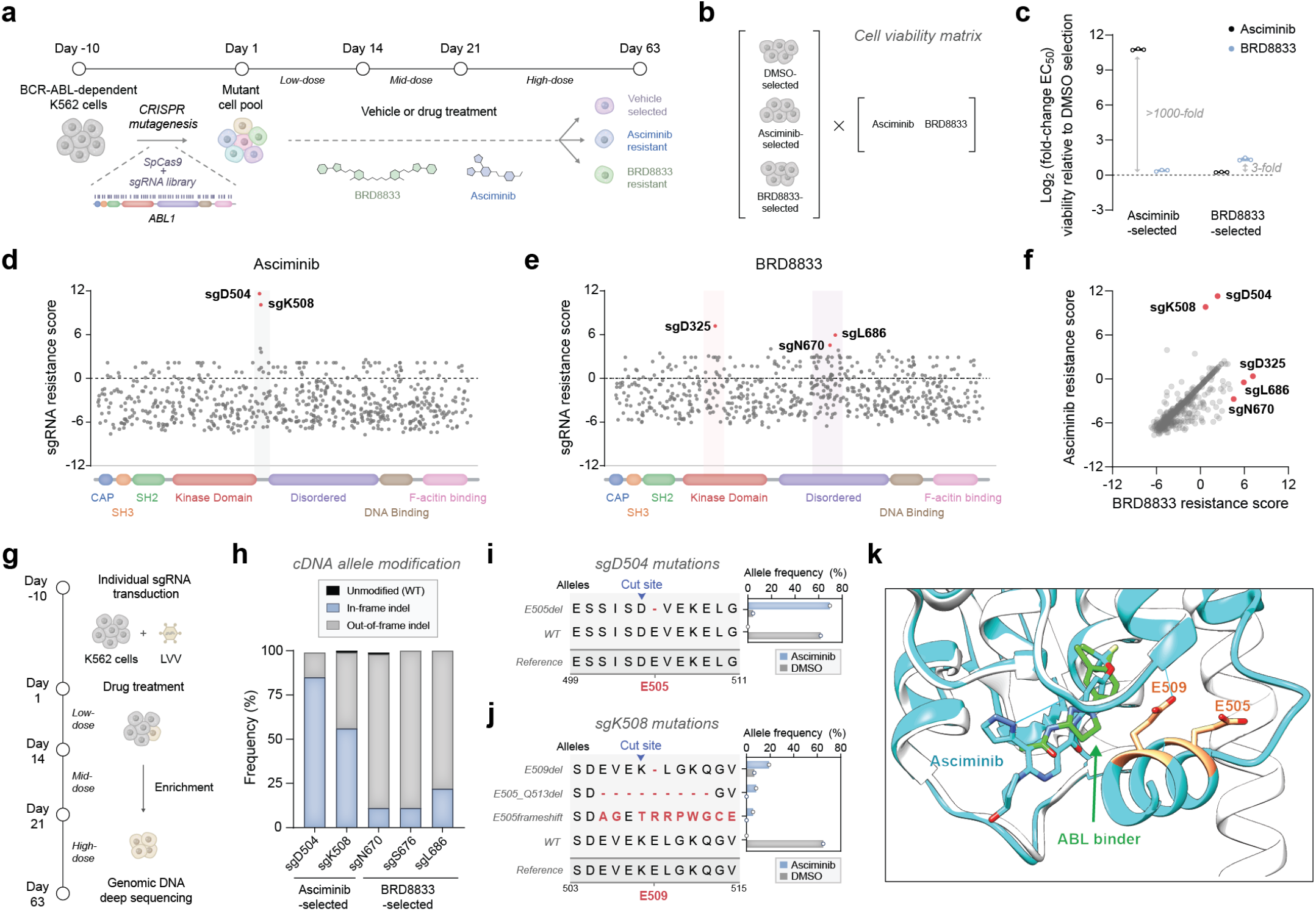
CRISPR-suppressor scanning identifies orthogonal resistance mutations for asciminib and BRD8833. **a** Schematic workflow for the CRISPR-suppressor scanning. **b–c** Cell viability matrix assay to measure drug-sensitivity after 63 days of selection. Resistance is expressed as log_2_-fold change relative to the DMSO selection (**c**). Mean and SD of three independent replicates are shown. **d–e** sgRNA resistance scores in the CRISPR-suppressor scanning. K562 cells were collected after 63 days of treatment with either vehicle, asciminib (**d**) or BRD8833 (**e**) following a dose-escalation schedule. Genomic DNA was extracted and sgRNA abundance was determined by sequencing. Scatter plots show resistance scores, calculated as z-scored log2-fold changes in sgRNA abundance in drug-treated conditions relative to the vehicle (DMSO)-treated population. sgRNA selected for further validation are annotated with their predicted cleavage site and highlighted in red. Datapoints represent the mean resistance scores across three replicate treatments. **f** Scatter plot comparing sgRNA resistance scores following 63 days of dose-escalated selection with asciminib or BRD8833. **g** Schematic for sgRNA validation and analysis of drug-resistant cells. **h** Targeted sequencing of sgRNA target sites and quantification of indel-modified sequences. **i–j** Translation of cDNA allele frequency showing the number of modifications induced by sgD504 (**i**) and sgK508 (**j**). Red residues highlight hotspot mutations E505 and E509. **k** Overlay of co-crystal structures for the myristoyl pocket binding asciminib (cyan, PDB: 5MO4) or the ABL binder in BRD8833 (gray/green, PDB: 6NPU). Mutated residues identified in the CRISPR-suppressor scanning are highlighted in orange on the asciminib-bound structure (cyan).

To further study the divergence in drug resistance, we quantified sgRNA abundance in the pooled screen after 14, 21, and 63 days of selection, and observed a gradual sgRNA enrichment (**Figure 6d,e** and **Figure S9c–h**). After 63 days, asciminib-resistant cells were enriched in sgRNAs targeting residues D504 and K508 (log_2_-fold change >9 compared to vehicle-treated cells) near the drug-binding site at the interface between ABL kinase domain and the disordered region (**Figure 6d**). In contrast, and consistent with the observed weaker resistance, treatment with BRD8833 showed dispersed sgRNA enrichment primarily deep within the disordered region (N670 and L686) or the kinase domain (D325) (**Figure 6e**). Interestingly, further comparing sgRNA enrichment between BRD8833 and asciminib revealed a single sgRNA targeting the S676 residue being enriched in both selections after 21 days (**Figure S9h**), before the sgRNA enrichment diverged by day 63 (**Figure 6f**).

Next, we individually delivered the top enriched sgRNAs in K562 cells to validate the resistance (**Figure 6g**). We subjected the transduced cells to BRD8833 or asciminib treatment, gradually increasing the drug pressure over 63 days (**Figure S10a**). One sgRNA transduction, sgD325, did not recover growth after 63 days of treatment with BRD8833 and was excluded from further analysis. sgRNAs targeting D504 and K508 conferred a strong resistance against asciminib, shifting the EC_50_ from 5 nM to 6500 nM and 900 nM, respectively (**Figure S10b–e**). Confirming the observed orthogonality from the pooled screen, the same cells did not show resistance against BRD8833, maintaining an EC_50_ of 60 nM and 59 nM, respectively. Conversely, sgRNA targeting N670, S676, and L686 emerging from the BRD8833 selection showed ubiquitous resistance against both asciminib and BRD8833 but shifted the EC_50_ by less than 10-fold. We genotyped editing outcomes at the sgRNA target sites to identify the underlying resistance mutations. sgD504 and sgK508 transduced K562 cells treated with asciminib showed predominantly in-frame mutations that result in E505 or E509 deletions near its binding site, where drug resistance frequently emerges (**Figure 6h–j and Figure S10f**,**g**). In contrast, resistance to BRD8833 caused by sgRNA targeting residues N670, S676, and L686 were characterized by large deletions resulting from frameshifts (**Figure 6h and Figure S10h-j**). This could lead to a partial loss of BCR-ABL activity, which explains both the impaired growth rate and weaker drug resistance. The distinct deletion patterns around residues E505 and E509 suggest that the myristoyl binding pocket conformation^40^ plays a crucial role in the orthogonal drug resistance (**Figure 6k**). Residues E505 and E509 are proximal to asciminib, but distal when bound to the ABL binder used in BRD8833. Deleting these residues likely disrupts helical folding or interactions with asciminib, while BRD8833 can effectively bind BCR-ABL regardless of these deletions.

## Discussion

The herein developed BCR-ABL-targeting CIP, BRD8833, introduces a fundamentally new modality in both proximity-mediated pharmacology and modulation of aberrant enzymatic activity by exploiting a natural regulatory mechanism of inhibitory phosphorylation via electric field effects. BRD8833 harnesses the aberrant kinase activity and redirects it against itself to achieve a pharmacologically beneficial effect. This self-targeting leads to phosphorylation in the kinase’s active site, and the high charge density of the phosphate group significantly perturbs protein structure, stability, dynamics, and electrostatic interactions.^41^ MD simulations show that the proximity of the phosphorylation site to the ATP-binding pocket and the DFG/HRD motifs likely invokes multiple inhibitory mechanisms, including perturbation of local BCR-ABL structure, ATP-binding, and kinase-substrate engagement.^42, 43^ Importantly, BRD8833 is one of the first small molecules shown to directly induce targeted phosphorylation in an endogenous cellular context for a pharmacological benefit.

Because most enzymes feature active sites with serine, threonine, or tyrosine-containing loops,^44^ the phosphorylation-based inhibitory mechanism could extend beyond BCR-ABL to other kinases and other enzymes. We found that many therapeutically relevant kinases and even GTPases feature P loop residues that can be phosphorylated (**Figure S5c**). This could provide new opportunities for regulating GTPase activity, a target class that has historically been considered challenging to target with conventional drug modalities.^45^ Although our study employs non-inhibitory ABL binders, we have successfully developed CIPs from existing kinase inhibitors via group-transfer chemistry.^46^ Thus, extant kinase inhibitors can be repurposed to induce inhibitory phosphorylation on a given kinase. For non-kinase targets, the CIP design requires selecting a partnering kinase based on the consensus phosphorylation motif and the kinase abundance in the target cell. Notably, our approach bypasses *de novo* ligand discovery and extensive medicinal chemistry optimization, which can be resource and time-intensive. Here, the catalytic turnover also potentially enables rapid attainment of high potency. Thus, by leveraging extant inhibitors, appropriate CIPs that induce inhibitory phosphorylation or other posttranslational modifications can be developed rapidly and cost-effectively.

Resistance inevitably develops as cancer cells adapt while drugs remain unchanged, pointing to the need for therapies with different resistance profiles^37^─BRD8833 exemplifies this principle. While binding to the same site as asciminib, its distinct mechanism effectively induces apoptosis in asciminib-resistant ABL fusions. CRISPR-suppressor scanning confirmed that BRD8833 can overcome drug resistance mutations against asciminib, showing less than a two-fold reduction in efficacy compared to the ∼1000-fold loss observed for asciminib. BRD8833 was also effective in gatekeeper mutation, but that is not surprising since those mutations occur in the ATP pocket,^6^ which is distant from the myristoyl pocket targeted by BRD8833. A frequently observed resistance mechanism to event-driven pharmacological agents is the alteration in expression levels of the effector enzyme (e.g., E3 ligase complex for PROTACs), which, unlike BCR-ABL, are often non-essential for cancer survival and growth.^47^ Finally, we showed in a PRISM multiplexed cancer cell line screen that BRD8833 does not target non-CML cell lines, comparable to the approved drug imatinib, pointing to the low off-targets.^33^ Beyond their therapeutic promise, CIPs that perturb electric field mechanisms near the enzyme’s active site may also serve as valuable tools to elucidate new aspects of target biology by synthetically inducing native or neophosphorylation. Taken together, we herein demonstrate a rapid, orthogonal strategy to overcome resistance and expand both research and clinical options in oncology.

## ACKNOWLEDGEMENT

This work was supported by the NIH (R01GM132825, R01 GM137606, R01 DK132900, R21AI178690, and R35-GM141930). M.M was supported by the Swiss National Science Foundation (P500PN_214260). Computational resources from ACCESS (project BIO230101) are acknowledged. Part of the computational work was performed on the Shared Computing Cluster, which is administered by Boston University’s Research Computing Services. We thank K. Tran (Broad Institute) for assisting with chemical compound characterization and structural analysis. We thank J. Dreyfuss and H. Pan from the Joslin Diabetes Center Bioinformatics and Biostatistics Core for performing the RNA-seq analysis.

