## Supporting Information for "Proximity-induced rewiring of oncogenic kinase triggers apoptosis"

### Table of contents

#### Supporting Methods


**Experimental Methods** 3

**Chemical Synthesis** 17


#### Supporting Figures


**Figure S1.** CIP **2** is effective against BCR-ABL-dependent K562 cells 49

**Figure S2.** Structure-activity relationship of BCR-ABL CIP 50

**Figure S3.** BRD8833 binds to the myristoyl-binding pocket to induce BCR-ABL phosphorylation 51

**Figure S4.** Analysis of key distances and dihedral angles during molecular dynamics simulations of BCR-ABL 52

**Figure S5.** S/T/Y residues in the P-loop of kinases and GTPases 53

**Figure S6.** Gene expression and gene set enrichment analysis 54

**Figure S7.** PRISM screen demonstrates selective targeting of BCR-ABL dependent cell lines 55

**Figure S8.** BRD8833 targets imatinib-resistant BCR-ABL mutations 56

**Figure S9.** CRISPR-suppressor scanning identifies drug-resistant K562 cell lines 57

**Figure S10.** CRISPR-suppressor scanning reveals different resistance profiles for asciminib and BRD8833 58

**Figure S11.** Uncropped western blots used in Fig. 2f 59

**Figure S12.** Uncropped western blots used in Fig. 4a 60

**Figure S13.** Uncropped western blots used in Fig. 4b 61


#### Supporting Tables


**Table S1.** LgBt and SmBt constructs to measure ternary complex formation 62

**Table S2.** Drug dosing regimen for CRISPR-suppressor scanning 64

**Table S3.** Oligonucleotides used for sgRNA validation 65

**Table S4.** Primers used for indel sequencing 66


#### Notes for Supporting Data


**Data Notes S1.** Global phosphoproteomic data 67

**Data Notes S2.** RNA-Seq data 67

**Data Notes S3.** PRISM screening data 67

**Data Notes S4.** CRISPR-suppressor scanning 67

**Data Notes S5.** Deep sequencing of genomic DNA 67

**Data Notes S6.** Deep sequencing of cDNA 67



#### Supporting References 68

### EXPERIMENTAL METHODS

**Chemical reagents and synthesis.** Imatinib and asciminib were purchased from commercial sources and used without further purification (MedChemExpress, #HY-15463 and #HY-104010). Compounds were used as solutions in 100% DMSO and stored in the freezer at -20°C. Detailed descriptions for the synthesis of all chemical compounds and their characterization are provided in the Chemical Synthesis Methods.

**Cell Culture.** The following cell lines were used: K562 (ATCC, #CCL-243), KCL22 (ATCC, #CCL-3349), BV-173 (DSMZ, #ACC 20), EM-2 (DSMZ, #ACC 135), KYO-1 (DSMZ, #ACC 601), and stable Ba/F3 cells containing a TEL-ABL fusion, an E255V mutation, or a T315I mutation (gifts from J. Griffin and E. Weisberg). Cells were cultured in RPMI media (Gibco, #22400-089) with 10% fetal bovine serum (FBS; Gibco, #16140-071) and 1% penicillin-streptomycin (penstrep; Gibco, #15140-122). HEK293T cells (ATCC, #CRL-3216) were cultured in DMEM + GlutaMAX-I (Gibco, #10564-011) with 10% fetal bovine serum (FBS; Gibco, #16140-071) and 1% penicillin-streptomycin (penstrep; Gibco, #15140-122). Cell cultures were routinely checked for contamination with mycoplasma.

**Cell viability assays.** Compound stock solutions were prepared at a concentration of 10 mM in anhydrous DMSO and dispensed into white-bottom 384-well plates using a Tecan D300e digital dispenser. The compounds were plated either at a maximum concentration of 20 μM, followed by a 12-point, three-fold serial dilution, or at a maximum concentration of 10 μM, followed by a 14-point, two-fold serial dilution. All wells were normalized to a final DMSO concentration of 0.5% (v/v). Cells were seeded at a density of 2,000 cells per well in 25 μL of complete medium (containing 10% (v/v) FBS and 1% (v/v) penicillin/streptomycin) and incubated for 72 hours in a humidified incubator at 37°C with 5% $CO_2$. Cell viability was assessed using the CellTiter-Glo (Promega, G7572) assay, with reagent added at a volume equal to half the total well volume. Luminescence was measured using an Envision plate reader, and percentage viability was calculated relative to DMSO-only controls. Dose-response data were fitted to a four-parameter logistic curve using GraphPad Prism 10. $EC_{50}$ values were determined from at least three independent replicates for all cell lines.

**Protein expression and purification.** *ABL* gene containing the SH3SH2 domain was cloned into the pET23a-His10X vector and expressed with a His-tag. Plasmids were co-transfected with YpoH shuttle vector and transformed into BL21(DE3) competent cells (Thermo Fischer Scientific, #EC0114). The primary culture was inoculated using a single colony grown overnight in 10 ml of TB media containing 50µg/ml of ampicillin (GoldBio, #A-301-5) and spectinomycin (Millipore, #Y000065) at 37°C with shaking at 200 rpm. The secondary culture was prepared by inoculating 1% of the primary culture into 1 L of TB media with ampicillin and spectinomycin until it reached an $OD_{600nm}$ of 0.3. The culture was moved to 18°C and further grown to $OD_{600nm}$ of 0.6 before adding 0.5 mM isopropyl-β-D-1-thiogalactosidease (Thermo Fischer Scientific, #15529019) to induce protein expression. The culture was grown for 16 hours at 200 rpm and harvested by centrifuging at 6000 rpm (Thermo Fischer Scientific, Sorvall LYNX 6000) for 10 minutes at 4°C. The supernatant was discarded, and the cell pellet was resuspended in lysis buffer (20 mM Tris-Cl, pH 7.4, 150 mM NaCl, 1 mM TCEP, and 10% glycerol). All subsequent steps were performed at 4°C in the cold room. Cell lysis was initiated by adding 1 mg/ml of lysozyme, 0.5 mM PMSF (Thermo Fischer Scientific, #36987), and 1X protease inhibitor cocktail (Pierce, #A37989) (one tablet in 1 ml of lysis buffer). The cells were incubated for 1 hour with gentle rocking, and 0.5% Triton X-100 (Sigma, #X100) was added. The mixture was further incubated for 15 minutes, and the cells were sonicated using a 250-watt ultrasonic processor (Fischer Scientific, #SFX250) with a 6 mm diameter probe, applying a cycle of 8 seconds "ON" and 15 seconds "OFF" for 30 minutes. The lysate was centrifuged at 16000 rpm for 1 hour. The cell debris was discarded, and the protein was purified from the clarified lysate by affinity chromatography, incubating it with 1.4 ml of pre-equilibrated His-Pur Nickel nitriloacetic acid (Ni-NTA) resin (Thermo Scientific, #88221) overnight. The resin was washed thrice with 15 column volumes of washing buffer (20 mM Tris-Cl, pH 7.4, 500 mM NaCl, 40 mM imidazole, and 10% glycerol). Histidine-tagged protein was eluted using elution buffer (20 mM Tris-Cl, pH 7.4, 150 mM NaCl, 300 mM imidazole, and 10% glycerol). The collected protein was concentrated using 10 KDa cut-off Amicon spin concentrators (Millipore, #UFC901008), centrifuged at 3500 rpm for 45 min, and subjected to gel filtration chromatography using an AKTA pure system (Cytiva) and a HiLoad Superdex-200 column, 120 ml PG (Cytiva, #28989335) at a flow rate of 1 ml/min with running buffer (20 mM Tris-Cl, pH 7.4, 150 mM NaCl, 10% glycerol). The protein-containing fractions were analyzed using NuPAGE Bis-Tris gels with a 4-12% gradient (Thermo Scientific, #NP0321BOX). The purified protein was quantified using BCA (Pierce, #A55864), snap-frozen in liquid nitrogen, and stored in aliquots at -80°C until future use.

**Differential scanning fluorimetry.** Thermal shift assays were performed using differential scanning fluorimetry (DSF) to assess the ABL thermal stability in the presence or absence of BRD8833 or control ABL binder BRD6501. Reactions were prepared in a final volume of 25 µl in DSF assay buffer (20 mM Tris-Cl-7.5, 150 mM NaCl and 1mM TCEP), containing 4 µM of purified ABL1-kinase domain, 10 µM of test compound or vehicle control, and Protein Thermal Shift™ dye (Thermo Scientific, #4461146) at a final dilution of 1:1000 from the manufacturer's stock. Samples were dispensed into a 384-well optical PCR plate (Applied Biosystems), sealed with optically clear film, and briefly centrifuged to remove bubbles. Thermal denaturation was monitored on a QuantStudio 7 Real-Time PCR System (Applied Biosystems) by increasing the temperature from 25 °C to 95 °C in 0.5 °C increments every 30 seconds. The melting temperature (Tm) was determined using the Protein Thermal Shift™ Software V1.4.

**TR-FRET binding assay.** TR-FRET assay was performed using CoraFluors, following previously reported procedures.[1,2] In brief, anti-GST nanobody (Proteintech, #st-250) was buffer exchanged into reaction buffer (100 mM sodium carbonate buffer, pH 8.5 + 0.05% Tween20 [vol/vol]) using a 7 kDa cutoff centrifugal concentrator (Thermo Scientific, #89877) following the manufacturer's instructions. A 2.5 mM solution of CoraFluor-1-Pfp in dimethylformamide was added at 12–15 fold molar excess to the antibody and left to react for 1 hour at room temperature. Excess reagents and organic solvent were removed, and the buffer was changed to TR-FRET assay buffer (25 mM HEPES pH 7.4, 150 mM NaCl, 1 mM DTT, 0.5 mg $ml^{-1}$ BSA, 0.005% TWEEN-20 [vol/vol]) using a 7 kDa cutoff centrifugal concentrator (Thermo Scientific, #89877).

Compounds were prepared in a 15-point, 2-fold serial dilution in a white 384-well plate (product reference) using a Tecan D300 digital dispenser. BRD6501 and BRD8833 were tested at a maximum concentration of 10 µM, and asciminib was tested at a maximum concentration of 1 µM. Next, TR-FRET assay buffer containing GST-ABL1 (Sino Biological, #11199-H09B, 2 nM), anti-GST nanobody conjugated to CoraFluor-1 (4 nM), and FITC-tracer (VS1231, 670 nM) were added, and the plate was incubated for 2 hours at room temperature. TR-FRET was measured on a Tecan Spark plate reader (SPARKCONTROL software version V2.1, Tecan Group) using an excitation wavelength of 340/50 nm and reading emission at 490/10 nm and 520/10 nm. The delay time between excitation and measuring emission was set to 100 µs, and the signal was measured over 400 µs (integration time). Prism 10 was used to fit the data to a four-parameter dose–response curve. The binding affinities ($K_D$ values) were calculated using the Cheng Prusoff equation (1):

$$K_\mathrm{D} = \frac{\mathrm{EC}_{50}}{1+\frac{[\mathrm{S}]}{K_\mathrm{m}}} \qquad (1)$$

Where $K_m$ is the affinity constant of the tracer, $EC_{50}$ is the concentration at which the test ligand displaces 50% of the tracer, and [S] is the tracer concentration.

**Analytical ultracentrifugation.** Analytical ultracentrifugation (AUC) experiments were performed using a Beckman Optima XL-I analytical ultracentrifuge equipped with an absorbance-based detection system.[3] A two-sector charcoal centerpiece with a 1.2 cm path length and sapphire windows was used. Purified proteins were dialyzed overnight against buffer A (25 mM Tris, pH 7.4, 150 mM NaCl) and samples were prepared by mixing ABL with BRD6501, BRD8833, or vehicle at an equimolar concentration of 1.5 µM while maintaining a DMSO concentration at 2%. The sample cell was loaded with 390 µL of protein solution, while the reference cell contained 400 µL of buffer A. Centrifugation was performed at 40,000 rpm (160,992 × g) at 20°C for 12–14 hours. Radial absorbance scans at 280 nm were collected every 3 minutes throughout the experiment and the resulting data analyzed with 'SedFit analysis' program using a continuous distribution(s) model based on the Lamm equation (2):

$$\frac{\partial \chi}{\partial t} = \frac{1}{r}\left\{\frac{\partial}{\partial r}\left[rD\frac{\partial \chi}{\partial r} - s\omega^2 r^2 \chi\right]\right\} \qquad (2)$$

Where $\chi$ is the concentration, $D$ is the diffusion coefficient and $s$ is the sedimentation coefficient of the particle, $t$ is the time, $r$ is the radius from the center of rotation, and $\omega$ is the angular velocity of the rotor. Parameters including buffer density ($\rho$ = 1.0059), buffer viscosity ($\eta$ = 0.0105443 poise), and partial specific volume ($\upsilon$ =

0.73125) of protein were measured by the Sednterp program. Standard sedimentation coefficients of samples were represented as S20,w referring to sedimentation at 20°C.

**NanoBiT ternary complex formation.** Full-length *BCR-ABL1* (GenScript) was fused N-terminally with *LgBt* and *SmBt* (sequences shown in **Table S1**) and cloned into the pCDNA3.1 vector backbone using NEBuilder HiFi DNA Assembly Master Mix (New England Bio Labs, #E2621L) according to the manufacturer's protocol. The NanoBiT assays were performed according to the manufacturer's protocol (Promega, #N2014) for adherent cells. Briefly, HEK-293T cells were grown overnight in DMEM (Thermo Fischer, #11965118) containing 10% FBS (Gibco, #16140071), washed twice with 1X PBS, trypsinized (Gibco, #25200056), and centrifuged for 5 minutes at 200 g. Cells were resuspended in growth media and counted on a CountessTM3 (Invitrogen, #AMQAZ2000). A transfection mixture was prepared by mixing BCR-ABL fusion constructs (1:3 ratio) with carrier DNA (Promega, #N2641), resulting in a DNA concentration of 10 µg/ml in 1 ml of Opti-MEM I without phenol red (Life Technologies, #11058-021) containing 1 % FBS. 35 µl of FuGENE HD Transfection Reagent (Promega, #E2311) was added to 1 ml of DNA mixture, the solution was mixed by inversion 5–10 times, incubated at room temperature for 30 minutes, and added to 20 ml of HEK-293T cell suspension of 0.2 million cells/ml in Opti-MEM assay solution. The resulting solution was gently mixed by inversion five times, and 95 µl of the final solution was dispensed in each well of the 96-well plate (20,000 cells/well, Corning, #3917). After 22–24 h, compounds or vehicle (DMSO) were added in 10 µl growth media, and 96-well plates were incubated at 37°C in a 5% $CO_2$-humidified atmosphere for 2 h. Compounds were tested in three independent replicates. After incubation, the plates were equilibrated at room temp for 15 minutes. The detection reagent was prepared by mixing 1 volume of Nano-GloR Live Cell Substrate (Promega, #N2011) with 19 volumes of Nano-GloR LCS Dilution Buffer. 50 µl was added to each well and the plate incubated for 2-3 minutes at room temperature before measuring the luminescence with 500 ms integration time was measured on an EnVision multimode plate reader (PerkinElmer). Raw data is processed by subtracting the background values and plotted using GraphPad Prism version 10.1.2.

**Phosphoproteomic analysis.** 50 x $10^6$ K562 cells were treated for 4 hours with DMSO, BRD8833 (1 µM), BRD6501 (2 µM), or asciminib (1 µM) at a density of 1x$10^6$ cells per ml. After compound treatment, cells were washed with ice-cold PBS, pelleted, and stored at -80 ˚C until further use. The cell pellet was then treated (with gentle end-over-end-rotation) for one hour with M-PER (Mammalian Protein Extraction Reagent, ThermoFisher, #78501) supplemented with 1X cOmplete EDTA-free Protease Inhibitor (Roche, #4693159001) and 1X PhosSTOP Phosphatase Inhibitor (Roche, #4906837001). The cell lysates were centrifuged at 16,000 x g for 10 minutes at 4°C, and the total protein concentration in the resulting supernatant was estimated with a Pierce BCA Protein Assay Kit (ThermoFisher, #23225). Immunoprecipitation (IP) of the BCR-ABL protein was performed using the Dynabeads Protein G (ThermoFisher, #10004D) according to manufacturer instructions, using an ABL primary antibody (Cell Signaling Technology, #2862) at a 1:100 dilution (1 μg ABL Ab / 100 μg total lysate) and overnight protein binding at 4 °C on a rotator. Unbound and non-specific proteins were washed out, and bead-bound proteins were eluted with 1X Sample buffer (BioRad, #1610747) using 2-mercaptoethanol and heating at 95°C for 5 minutes. IP eluates were resolved by SDS-PAGE, gel bands excised, and proteins reduced with 1 mM dithiothreitol (DTT) at 60°C for 30 minutes. Samples were alkylated with 5 mM iodoacetamide for 15 minutes at room temperature in the dark and subjected to in-gel trypsin digestion. Briefly, gel pieces were washed, dehydrated with acetonitrile, and dried in a speed-vac (Eppendorf Vacufuge Plus) before rehydration at 4°C with 50 mM ammonium bicarbonate containing 12.5 ng/μL sequencing-grade trypsin (Promega, #V5111). Samples were digested overnight at 37°C. Peptides were then extracted with 50% acetonitrile and 1% formic acid, dried in a speed-vac (Eppendorf Vacufuge Plus), and stored at 4°C until analysis. Phosphoproteomic analysis was performed at the Taplin Biological Mass Spectrometry Facility (Harvard Medical School). In brief, samples were reconstituted in 5–10 μl of HPLC solvent A (2.5% acetonitrile, 0.1% formic acid) and separated using a nano-scale reverse-phase HPLC capillary column (100 μm inner diameter x ~30 cm length) packed with 2.6 μm C18 silica beads. Peptides were loaded via a Thermo EASY-LC system (Thermo Scientific) and eluted using a 130 min gradient (0.0-1.0 min: 3%-5% ACN, 1.0-69.0 min: 5%-25% ACN, 69.0-89.0 min: 25%-40% ACN, 89.0-104.0 min: 40%-60% ACN, 104-120 min: 95% ACN) at a flow rate of 0.3 μL/min. Eluted peptides were subjected to electrospray ionization and analyzed on an Orbitrap Fusion Lumos mass spectrometer (Thermo Scientific). The raw data were analyzed using Proteome Discoverer 2.5 software package (Thermo Scientific). Sequest HT was used to search against human tyrosine-protein kinase Abl (P00519) with a precursor mass tolerance of 10 ppm and fragment mass tolerance of 0.6 Da. A maximum of two missed cleavages and a minimum peptide length of six were selected. Search parameters included dynamic modifications for methionine oxidation (15.995 Da), tyrosine phosphorylation (79.966 Da), N-terminal glutamate cyclization, and acetylation, as well as static cysteine alkylation (+57.021 Da). Target Decoy PSM Validator was applied with a false discovery rate (FDR) of 1%. IMP-ptmRS was used to calculate phosphorylation site probabilities, and phosphopeptide-containing groups were filtered. Labeled and unlabeled peptide precursor ion intensities were used to estimate labeling efficiency. The results were exported and analyzed using GraphPad Prism 9.0.

**Global phosphoproteomic analysis.** K562 cells were adjusted to a density of $1\times10^6$ cells/mL in complete growth media and treated in quadruplicate with vehicle (DMSO), 2 µM control compound BRD6501, or 1 µM BRD8833 for 4 hours. Upon completing the treatment, cells were harvested and washed with ice-cold PBS. For phosphoproteomics, $23\times10^6$ cells were pelleted and lysed in EasyPep Lysis Buffer (Thermo #A45735) with protease and phosphatase inhibitors (Thermo #78444), following the manufacturer's protocol. Protein concentration was determined using the Rapid Gold BCA Kit (Thermo #A53225) and ~500 µg of lysate were reduced with 5 mM TCEP (Thermo #20490) at 60°C for 30 min, alkylated with 10 mM iodoacetamide (Thermo #122270250) before quenching excess reagent with 10 mM DTT (Thermo #20290), and purified using SP3 resin beads (Hydrophilic and Hydrophobic, Thermo #09-981-121; 09-981-123). Next, Trypsin/Lys-C Mix (Promega #V5072) in 200 mM EPPS (pH 8.5) was added at a 1:100 ratio to digest the protein lysate at 37°C for ~16 hours. Resulting peptides were in 30% acetonitrile and labeled with TMTpro 18-plex reagents (Thermo #A52045) at a 1:1 ratio. Upon confirming the labeling efficiency (>95%), the reactions were quenched with 50% hydroxylamine (Thermo #90115). For total proteomic analysis, ~100 µg of pooled sample was cleaned using the SP3 resin protocol described above and fractionated with the Pierce High pH Reversed-phase Peptide Fractionation Kit (Thermo #84868). The resulting 10 fractions were subsequently analyzed by LC-MS/MS on an Orbitrap Eclipse Tribrid instrument as described below. For phosphoproteomic analysis, the remaining samples were dried by vacuum centrifugation, desalted (Peptide Desalting Spin Columns, Thermo #89851), and enriched for phosphotyrosine peptides using 30 µL of P-Tyr-1000 (Cell Signaling #8954) conjugated to protein A agarose (MilliporeSigma #11134515001) in IAP buffer (6-hour incubation at 4°C). After washing, phosphopeptides were eluted in 100 mM formic acid, desalted, and analyzed by LC-MS/MS.

For LC-MS/MS analysis, samples were reconstituted in 2% acetonitrile containing 0.2% formic acid, sonicated, and centrifuged at 10,000g for 1 min. Supernatants were transferred to LC-MS vials (Thermo Fisher, #6PK1655) and loaded onto a Vanquish Neo UHPLC system (Thermo Fisher) coupled to an Orbitrap Eclipse mass spectrometer with a NanoFlex ion source. For total proteomics analysis, samples were separated on a 60 cm column (IonOpticks, #AUR3-60075C18-TS) using a 120-minute gradient (0.0–1.0 min, 3%–5% ACN, 1.0–69.0 min, 5%–25% ACN, 69.0–89.0 min 25%–40% ACN, 89.0–104.0 min 40%–60% ACN, 104–120 min 95% ACN, wash) at a flow rate of 0.3 µL/min. MS1 data were acquired in Orbitrap mode with a resolution of 120,000, AGC target 400000, and maximum injection time 50 ms. Charge states from 2+ to 6+ were included, and a dynamic exclusion time of 60 seconds was used. MS2 scans were isolated with quadruple and fragmented using HCD with a fixed collision energy of 35% and a 0.7 m/z isolation window. The normalized AGC target for MS2 was set to 200%, and fragment ions were detected in the Orbitrap at a resolution of 50,000 with a defined first mass of m/z 120. Phosphoproteomic samples were separated on the 60 cm IonOpticks column (#AUR3-60075C18-TS) using a 180-minute gradient (0.0–6.0 min, 3%–5% ACN, 6.0–121.0 min, 5%–25% ACN, 121.0–152.0 min 25%–40% ACN, 152.0–161.0 min 40%–60% ACN, 161–180 min 95% ACN, wash) at a flow rate of 0.3 µL/min and the same MS acquisition parameters as described above for the total proteomic analysis.

Proteome Discoverer 2.5 (Thermo Fisher) was used to process the raw data, using Sequest HT for peptide identification against the human reference proteome (Uniprot proteome ID: UP000005640_9606) and a contaminant database.[4] The search criteria allowed up to two missed cleavages and included peptides with at least 6 amino acids. Precursor mass tolerance was set at 10 ppm, and fragment mass tolerance was fixed at 0.02 Da. Dynamic modifications included methionine oxidation (+15.995 Da) and phosphorylation (+79.966 Da) on serine, threonine, and tyrosine. N-terminal dynamic protein modifications included acetylation (+42.011 Da), methionine loss (-131.040 Da), and methionine loss together with acetylation (-89.030 Da). Static modifications included cysteine alkylation (+57.021 Da) and conjugation with TMT reagent (+304.207 Da) on peptide N-termini and lysines. Peptide-spectrum match (PSM) was validated using Percolator and a false discovery rate (FDR) of 0.01. IMP-ptmRS node was used for phosphosite localization. TMTpro isotopic impurity corrections were applied and the resulting data was exported as CSV for importing into Perseus,[5] in which reporter ion abundances were $log_2$-transformed and normalized by subtracting each column's median, replicates were grouped according to experimental conditions, and Welch's t-tests were performed to identify phosphopeptides with significantly altered abundances (see **Data S1**). Volcano plots were prepared in GraphPad Prism.

**Molecular Dynamics (MD) simulations.** Since no crystal structures of activated (i.e., DFG-in) BCR-ABL in complex with ATP · $Mg^{2+}$ are currently available in the Protein Data Bank,[6] structural models were generated via AlphaFold3.[7] The two top-ranked predictions were selected for MD simulations. In addition, a "hand-built" model was also constructed by aligning the alpha carbons of 2GQG[8] and 1B38[9], taking the coordinates of BCR-ABL from 2GQG and the coordinates of ATP · $Mg^{2+}$ from 1B38. For each model, four independent MD simulations were performed for four different states; i.e., apo wild-type (WT), apo Y253TP2, holo WT, and holo Y253TP2—using the CHARMM36 force field[10] in OpenMM 8.2.[11,12] Apoenzymatic states were created by removing the ATP · $Mg^{2+}$ ligand from the holoenzymatic structures. In the simulations, the protonation states of titratable residues were assigned using the PDB2PQR software package,[13-16] and the CHARMM-GUI web server[17] was used to solvate and minimize the resulting structures in 150 mM NaCl solutions of TIP3P water.[18,19] The box length of the simulation cell is in the range of 83–85 Å. Every system was equilibrated for 100–200 ns at 303.15 K and 1 bar with a 2 fs timestep, followed by 300 ns of production MD. All bonds involving hydrogen atoms were constrained with the SHAKE algorithm.[20,21] Nonbonded interactions were treated with a van der Waals (vdW) force-switching function employing a switching distance of 10 Å and cutoff distance of 12 Å. Electrostatic interactions were calculated using the Particle-Mesh-Ewald approach, with a real space cutoff of 12 Å and an error tolerance of $5\times10^{-4}$.[22] Temperature was regulated by a Langevin thermostat with a collision frequency of 1 $ps^{-1}$, and pressure was maintained by a Monte Carlo barostat with a coupling frequency of 5 $ps^{-1}$.

Analysis of the resulting trajectories was achieved utilizing the MDAnalysis program.[23,24] Representative snapshots were obtained by *k*-means clustering of each system's two largest principal components. Electrostatic potential calculations of the WT and Y253TP2 apoenzymes were performed using the Adaptive Poisson–Boltzmann Solver[15,16] plugin for PyMOL (Schrödinger INC., Version 3.0) at a temperature of 303.15 K and salt concentration of 150 mM; partial charges were taken from the CHARMM36 force field,[10] and vdW radii were used to define atomic volumes. All three structural models exhibited consistent qualitative behavior. That is, phosphorylation of Y253 substantially alters the P-loop's conformation and leads to the formation of a stable ionic cluster between ATP's triphosphate moiety, Y253TP2's phosphate group, R367, and several sodium ions. Results from simulations starting from the top-ranked AlphaFold3-generated model were used for all figures.

**Immunoblotting.** K562 cells were counted, and 3.5 × $10^6$ cells (at 1 × $10^7$ cells per ml) were taken for each condition. DMSO and solutions of the tested compounds (BRD6501: 2 µM; BRD8833: 1 µM) were prepared in complete media with a final concentration of 0.1% DMSO and were added to the cells. The cells were then incubated for different times depending on the effect to be measured: 4 hours for detecting BCR-ABL autophosphorylation (pY70, pY226, and pY393), 7 hours for downstream signaling (pERK, pSTAT5), and 70 hours for pCRKL signaling and Caspase8 and PARP cleavage. After incubation, the cells were harvested, washed with ice cold 1× PBS, and stored at -80°C until further processing. The cells were lysed using M-PER lysis buffer (ThermoScientific, #78505) containing cocktails of protease (Roche, #4693159001) and phosphatase inhibitors (Roche, #4906837001). The total lysates were spun at 16,000 g for 10 min at 4°C, and the total protein concentration in the supernatant was estimated by BCA (cat# 23225, Thermo Scientific). An equal amount of total lysate (1 µg/µl) was run on the automated simple western (Jess) using all required reagents from ProteinSimple. The results were analyzed using the Compass SW software. Primary antibodies used for detecting the autophosphorylation of BCR-ABL, including pY393 (Cell Signaling, #2865), pY226 (Cell Signaling, #2861), pY70 (Cell Signaling, #3098), and cABL (Santa Cruz Biotechnology, #SC-23), as well as antibodies for detecting downstream signaling, such as pSTAT5 (Cell Signaling, #9351), STAT5 (Cell Signaling, #94205), pERK1/2 (Cell Signaling, #4370S), ERK1/2 (Cell Signaling, #4696S), pCRKL (Cell Signaling, #34940), CRKL (Cell Signaling, #38710), caspase-8 (Cell Signaling, #4790), PARP (Cell Signaling, #9532), and the β-actin (Cell Signaling, #4967, 3700) and GAPDH (Cell Signaling, #2118, 97166) loading controls. All primary antibodies were used at 1:50 dilutions. Secondary antibodies: anti-rabbit-HRP (cat# 042-206, Protein Simple), used without further dilution, and anti-mouse-NIR (cat# 043-821, Protein Simple), used at 1:20 dilution. BCR-ABL phosphosites were mapped to the human ABL1 sequence (UniProt accession: P00519). For antibodies that did not work in automated Simple Western, conventional Western blots were performed. Briefly, cell lysates containing equal amounts of total protein (20 µg) were heated with Laemmli reducing SDS sample buffer at 95–100°C for 5 min. The protein samples were then resolved using NuPAGE gels (4-12%), which were then dry-transferred to nitrocellulose membranes (iBlot2). The membranes were blocked with 5% milk in 1× TBST buffer, followed by incubation overnight at 4°C (in a cold room) on an orbital shaker with a 1:1000 dilution of primary antibodies. The membranes were then washed with 1× TBST buffer and were incubated further with the appropriate secondary antibodies, IRDye800CW-conjugated donkey anti-mouse (1:10,000; LiCor, #926-32212), and IRDye680RD-conjugated donkey anti-rabbit (1:10,000; LiCor, #926-68073), for 1 hour at room temperature. The membranes were again washed with 1× TBST buffer. Fluorescent blots were developed using the LiCor Odyssey CLx Imaging System. Uncropped western blot images are shown in **Figures S11–13**.

**Gene expression analysis.** K562 cells were treated with either DMSO (vehicle), asciminib (100 nM), BRD8833 (1 μM), or BRD6501 (2 μM) for 6, 12, or 24 hours in biological quadruplicate. After the indicated treatment time, the cells were harvested, pelleted, and shipped on dry ice to Genewiz (South Plainfield, NJ) for RNA extraction and RNAseq analysis. Genewiz first isolated total RNA using the RNeasy RNA isolation kit (Qiagen, #74104), and the quality of the sample extraction was analyzed via Nanodrop and Agilent Tapestation. mRNA isolation/library prep was performed by Poly-T oligonucleotide-mediated mRNA extraction through hybridization of the mature mRNA Poly-A tail using NEBNext Ultra II Directional RNA Library Prep Kit for Illumina (New England Biolabs, #E7760S). The overall quality of the library was analyzed using an Agilent Tapestation. The enriched mRNA was reverse transcribed into DNA with subsequent second-strand synthesis. Sequencing was performed by Genewiz using Illumina HiSeq, 2× 150 bp configuration, single-index sequencing platform. Sequencing depth was performed at ~37 M reads per sample, generating ~15 GB of data per sample. The data was downloaded via FTP from Illumina as a fastq file for analysis. Sequencing alignment and data analysis were performed by the Harvard Medical School Joslin Diabetes Center Bioinformatics & Biostatistics core. The aligned sequencing data were further investigated using moderated t-tests, principal component analysis, gene set enrichment analysis (GSEA), expression heatmaps, gene ontology (GO), Reactome, TFT pathway analysis, and KEGG pathway analysis. Weighed p-values, FDR, log fold-change, and fold-change of each treatment compared to the vehicle (DMSO) condition are provided in **Data S2**.

**PRISM multiplexed cancer cell line screening.** This assay was performed in collaboration with the Broad Institute PRISM team as described previously[25,26]. Two PRISM cell line collections were used: PR500, which consists exclusively of adherent cell lines, and PR300+, which includes both suspension and adherent cell lines. Briefly, lentivirally barcoded cells were cultured in pools of 20–25 cell lines in 384-well plates, with each pool consisting of mixed lineages grouped by doubling rate. Cells were treated with either BRD8833 (eight-point, three-fold serial dilution: 0.01 nM to 20 μM), BRD6501, Imatinib (eight-point, three-fold serial dilution: 0.005 nM to 10 μM), or with vehicle control, and incubated for 120 hours. Treatments were performed in three independent replicates. Following incubation, cells were lysed, and genomic DNA was extracted from the pooled cells. The unique DNA barcodes identifying each cell line were amplified by PCR and hybridized to Luminex beads conjugated with complementary antisense barcodes. The beads were then incubated with streptavidin-phycoerythrin to fluorescently label biotin moieties, and fluorescence signals were detected using Luminex FlexMap machines. Data were processed using a previously established standardized R pipeline (https://github.com/broadinstitute/prism_data_processing) to estimate drug sensitivities across different concentrations and fit dose-response curves. The area under the dose-response curve (AUC) was used to quantify and compare drug sensitivity across cell lines. Cell line annotations were obtained from the DepMap platform (https://depmap.org/portal/) to correlate target fusion gene expression with drug sensitivity. Detailed annotations of all cell lines and their respective AUC-based responses to BRD6501 and BRD8833 are provided in **Data S3**.

**sgRNA pooled cloning and CRISPR suppressor screening.** CRISPR suppressor screens were performed in K562 cells, which harbor the *BCR-ABL1* fusion gene and are sensitive to ABL1 knockout, as annotated by the DepMap CRISPR and RNAi database (https://depmap.org/portal/). To generate in situ mutational diversity of BCR-ABL, SpCas9 and a pool of all possible sgRNAs with an NGG PAM site within the *ABL1* coding sequence were used. sgRNAs were filtered to include only those with an off-target score (MIT Specificity Score) greater than 20 to minimize off-target effects. A total of 78 sgRNAs targeting non-essential genes (negative controls) and three sgRNAs targeting essential genes (positive controls) were also included in the pool. A complete list of protospacers targeting *ABL1* and control genes is provided in **Data S4**. The sgRNA library was synthesized as an oligonucleotide pool with corresponding homology arms for Gibson assembly (Twist Bioscience) and cloned into an Esp3I-digested lentiCRISPRv2 vector (Addgene #52961, a gift from Feng Zhang) using NEBuilder HiFi DNA Assembly Master Mix (New England Bio Labs, #E2621L) according to the manufacturer's protocol. The pooled plasmid library was electroporated into Endura DUO cells (Biosearch Technologies, #60242) using a Gene Pulser Xcell Electroporation System (Bio-Rad). To ensure adequate sgRNA coverage, the number of transformed cells was maintained at more than 1,000-fold the number of sgRNAs in the library. Lentiviral particles were produced using pMD2.G (Addgene #12259) and psPAX2 (Addgene #12260) packaging plasmids (both gifts from Didier Trono) and transduced into K562 cells via spinfection (1,800 rcf for 90 min) at a low multiplicity of infection (MOI < 0.3) to ensure integration of a single sgRNA per cell. Cells then underwent puromycin selection for >10 days and were split into three independent replicates for treatment with vehicle (0.1% DMSO), asciminib, or BRD8833 over a total of 63 days (details for the drug-dosing are provided in **Table S2**). Cell growth was recorded every 3–5 days to ensure the selection of drug-resistant cells. At 7, 21, and 63 days post-treatment, cells were harvested for sgRNA enrichment analysis. Genomic DNA was extracted using a QIAamp DNA Blood Mini Kit (Qiagen, #51104) following the manufacturer's protocol. The sgRNA regions were PCR-amplified to incorporate Illumina adapters, and the coverage and distribution of the sgRNA library were assessed using MiSeq (Illumina). Data analysis was performed using previously described Python scripts.[27] Briefly, sgRNAs were mapped to the protein amino acid positions based on the predicted cut site coordinates within the *ABL1* (NP_005148.2) coding sequence. Sequencing reads uniquely matching each sgRNA were converted to reads per million, increased by a pseudocount of 1, log2-transformed, and averaged across replicates for each condition. The values were normalized to the plasmid library. sgRNA resistance scores were then calculated by normalizing to the mean score of the negative control sgRNAs within each condition and subtracting the score in the vehicle condition from that in the compound-treated condition at matching time points. Final values were z-scored based on the mean and standard deviation of the negative control distribution.

**Validation of sgRNAs from the CRISPR-resistance screening.** The six most enriched sgRNAs (sgD325, sgD504, sgK508, sgN670, sgS676, and sgL686) from the pooled CRISPR suppressor screen were cloned into the LentiCRISPRv2 plasmid for validation. The vector was digested with BsmBI (New England Biolabs,

#R07399L) to generate 3’ and 5’ overhangs. Single-stranded protospacer oligonucleotides with complementary overhangs (Eton Bioscience) were phosphorylated using T4 Polynucleotide Kinase (New England Biolabs, #M0201S), annealed, and ligated into the digested vector with the Quick Ligation Kit (New England Biolabs, #M2200S). Oligonucleotide sequences for sgRNA validation are listed in **Table S3**. Lentiviral particles were produced and transduced into K562 cells as described for library preparation. Transduced cells underwent a 63-day drug selection following the outline in **Table S2**. Growth rates were measured every 3–5 days to ensure the selection of drug-resistant cells.

**Genotyping and indel sequencing of drug-resistant variants.** Genomic DNA from drug-selected cells was extracted using the QIAamp DNA Blood Mini Kit (Qiagen, #51104), following the same protocol used for sgRNA distribution analysis. At least 1 µg of genomic DNA was used as the template for the first round of PCR to amplify gene-specific amplicons with common overhangs (20 cycles, NEBNext Ultra II Q5 Master Mix, New England Bio Labs, #M0544L). For cDNA analysis, total mRNA was isolated from drug-selected cells using the RNeasy Mini Kit (Qiagen, #74106). cDNA was synthesized by RT-PCR using the SuperScript III First-Strand Synthesis SuperMix (Invitrogen, #18080-400). The resulting cDNA was then amplified in a first round of PCR, following the same conditions as genomic DNA amplification. PCR products were diluted 1:10 and subjected to a second round of PCR using primers with barcoded Illumina adapters (10 cycles, NEBNext Ultra II Q5 Master Mix). Primer sequences for both PCR rounds are provided in **Table S4**. The final PCR products were purified using SPRIselect bead clean-up (Beckman Coulter Life Sciences, #B23317), isolating fragments of approximately 500 bp according to the manufacturer’s protocol. The concentration of purified amplicons was determined by qPCR using the KAPA Library Quantification Kit (Roche, #07960140001) and adjusted to 2 nM. Final amplicons were sequenced on an Illumina MiSeq using a Nano Kit v2 (Illumina, #MS-103-1003) according to the manufacturer’s instructions. Sequencing reads were aligned to their reference amplicons using CRISPResso2 (ref[28]), and allele frequencies were calculated (see **Data S5,6**).

**CHEMICAL SYNTHESIS**

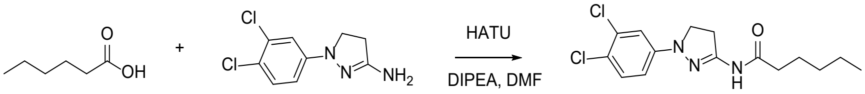

**Compound 1 (VS772): N-(1-(3,4-dichlorophenyl)-4,5-dihydro-1H-pyrazol-3-yl)hexanamide.** The synthesis and characterization of VS772 were previously reported (ref[29]). In brief, hexanoic acid (23 mg, 200 μmol), O-(7-Azabenzotriazol-1-yl)-N,N,N',N'-tetramethyl-uronium hexafluorophosphate (HATU) (100 mg, 260 μmol), and DIPEA (100 uL, 570 μmol) were combined in DMF (2 mL) and stirred at room temperature for 30 min. 1-(3,4-Dichlorophenyl)-4,5-dihydro-1H-pyrazol-3-amine 28 (46 mg) was added, and the reaction mixture was stirred at room temperature overnight. The solvent was removed under reduced pressure, and the residue was purified via flash column chromatography (Hex:EtOAc gradient from 100:0 to 70:30).

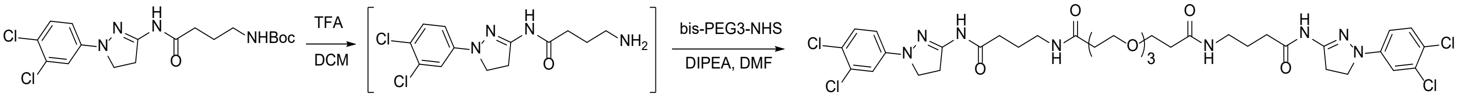

**Compound 2 (VS1143): 4,4'-((3,3'-((oxybis(ethane-2,1-diyl))bis(oxy))bis(propanoyl))bis(azanediyl))bis(N-(1-(3,4-dichlorophenyl)-4,5-dihydro-1H-pyrazol-3-yl)butanamide).** Tert-butyl (4-((1-(3,4-dichlorophenyl)-4,5-dihydro-1H-pyrazol-3-yl)amino)-4-oxobutyl)carbamate (ref[29]) (21 mg, 51 μmol) was dissolved in DCM (2 mL) and treated with TFA (500 μL). The reaction mixture was stirred at room temperature for 30 min and concentrated under reduced pressure. To the oil residue, bis-PEG3-NHS ester (11.1 mg, 25 μmol) in DMF (1 mL) was added, followed by DIPEA (50 μL). The reaction mixture was stirred at room temperature for 30 min, concentrated under reduced pressure, and purified by HPLC, affording the desired product VS1143 as a white powder (12 mg, 57% yield). $^1$H NMR (400 MHz, $CDCl_3$ + $CD_3OD$ (1:1)) δ 7.20 (d, J = 8.9 Hz, 2H), 6.99 (d, J = 2.7 Hz, 2H), 6.70 (dd, J = 9.1, 2.5 Hz, 2H), 3.76 – 3.65 (m, 8H), 3.60 (s, 9H), 3.40 (t, J = 9.8 Hz, 4H), 3.22 (t, J = 6.8 Hz, 4H), 2.42 (t, J = 6.1 Hz, 4H), 2.33 (t, J = 7.5 Hz, 4H), 1.80 ppm (p, J = 7.1 Hz, 4H). $^{13}$C NMR (400 MHz, $CDCl_3$ + $CD_3OD$ (1:1)) δ 172.34, 172.01, 148.96, 146.29, 132.02, 129.91, 120.24, 113.61, 111.70, 69.85, 69.72, 66.74, 38.11, 36.21, 33.12, 32.29, 24.64 ppm. HRMS (ESI-TOF): calculated for $C_{36}H_{47}Cl_4N_8O_7$ (M+H): 842.2244, found: 842.2243.

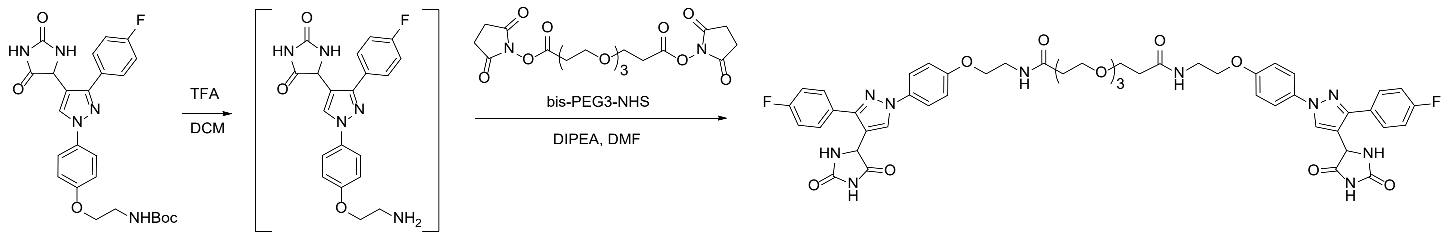


**Compound 3 (VS1524): 3,3'-((oxybis(ethane-2,1-diyl))bis(oxy))bis(N-(2-(4-(4-(2,5-dioxoimidazolidin-4-yl)-3-(4-fluorophenyl)-1H-pyrazol-1-yl)phenoxy)ethyl)propanamide).** *Tert*-butyl (2-(4-(4-(2,5-dioxoimidazolidin-4-yl)-3-(4-fluorophenyl)-1H-pyrazol-1-yl)phenoxy)ethyl)carbamate (ref[29]) (25 mg, 51 µmol) was dissolved in DCM (2 mL) and treated with TFA (500 µL). The reaction mixture was stirred at room temperature for 30 min and concentrated under reduced pressure. To the oil residue, bis-PEG3-NHS ester (11.1 mg, 25 µmol) in DMF (1 mL) was added, followed by DIPEA (50 µL). The reaction mixture was stirred at room temperature for 30 min, concentrated under reduced pressure, and purified by HPLC, affording the desired product VS1524 as a white powder (15 mg, 60% yield). $^1$H NMR (400 MHz, $CDCl_3$ + $CD_3OD$ (1:1)) δ 8.03 (s, 2H), 7.81 – 7.67 (m, 4H), 7.66 – 7.50 (m, 5H), 7.13 (t, J = 8.7 Hz, 4H), 7.03 – 6.92 (m, 4H), 5.21 (d, J = 1.4 Hz, 2H), 4.03 (t, J = 5.4 Hz, 4H), 3.69 (t, J = 6.0 Hz, 4H), 3.62 – 3.47 (m, 12H), 2.45 ppm (t, J = 6.0 Hz, 4H). $^{13}$C NMR (101 MHz, $CDCl_3$ + $CD_3OD$ (1:1)) δ 173.45, 165.00, 162.54, 158.74, 158.48, 152.06, 134.06, 131.15, 131.07, 128.89, 128.35, 121.70, 116.17, 116.11, 115.96, 115.79, 70.80, 70.70, 67.67, 67.39, 54.91, 39.47, 37.06 ppm. HRMS (ESI-TOF): calculated for $C_{50}H_{50}F_2N_{10}O_{11}Na$ (M+Na): 1027.3521, found: 1027.3522.

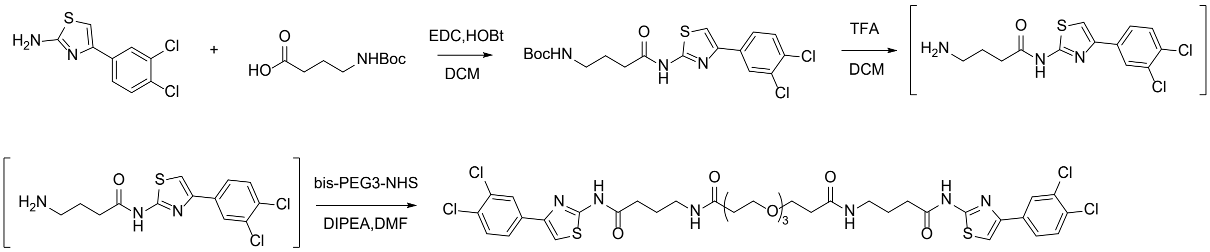

**Compound 4 (VS1189): 4,4'-((3,3'-((oxybis(ethane-2,1-diyl))bis(oxy))bis(propanoyl))bis(azanediyl))bis(N-(4-(3,4-dichlorophenyl)thiazol-2-yl)butanamide).** 4-(3,4-dichlorophenyl)thiazol-2-amine (24 mg, 0.1 mmol) was combined with N-Boc gamma-amino butyric acid (22 mg, 0.11 mmol), 1-Ethyl-3-(3-dimethylaminopropyl)carbodiimide (EDC) (77 mg, 0.4 mmol), and 1-Hydroxybenzotriazole (61 mg, 0.45 mmol) in 3 ml dichloromethane. Reaction was stirred at room temperature overnight (16 h), and the product isolated following previously reported procedures (ref[30]). Tert-butyl (4-((4-(3,4-dichlorophenyl)thiazol-2-yl)amino)-4-oxobutyl)carbamate (43 mg, 100 µmol) was dissolved in DCM (4 mL) and 1 mL TFA was added. The reaction mixture was stirred at room temperature for 30 min and concentrated under reduced pressure. To the oil residue, bis-PEG3-NHS ester (22.2 mg, 50 µmol) in DMF (2 mL) was added, followed by DIPEA (100 µL). The reaction mixture was stirred at room temperature for 30 min, concentrated under reduced pressure, and purified by HPLC, affording the desired product VS1189 as an off-white powder (7.9 mg, 18% yield). $^{1}$H NMR (400 MHz, DMSO-$d_6$) δ 12.21 (s, 2H), 8.11 (d, $J$ = 2.0 Hz, 2H), 7.91 – 7.82 (m, 4H), 7.79 (s, 2H), 7.67 (d, $J$ = 8.4 Hz, 2H), 3.58 (t, $J$ = 6.5 Hz, 4H), 3.48 – 3.42 (m, 8H), 3.08 (q, $J$ = 6.6 Hz, 4H), 2.45 (t, $J$ = 7.4 Hz, 4H), 2.29 (t, $J$ = 6.5 Hz, 4H), 1.73 (p, $J$ = 7.2 Hz, 4H). $^{13}$C NMR (101 MHz, DMSO-$d_6$) 171.36, 170.04, 158.21, 146.11, 134.87, 131.53, 130.98, 129.96, 127.30, 125.65, 109.95, 69.64, 69.51, 66.83, 37.86, 36.18, 32.34, 24.67. HRMS (ESI-TOF): calculated for $C_{36}H_{40}Cl_4N_6O_7S_2$ (M+H): 873.1227, found: 873.1220.

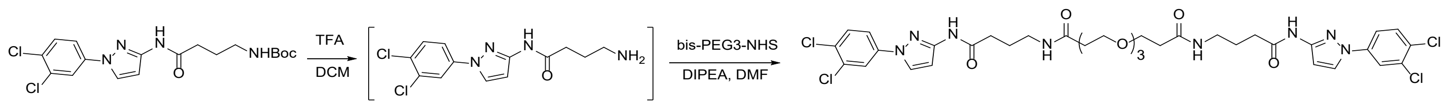


**Compound 5 (VS1144): 4,4'-((3,3'-((oxybis(ethane-2,1-diyl))bis(oxy))bis(propanoyl))bis(azanediyl))bis(N-(1-(3,4-dichlorophenyl)-1H-pyrazol-3-yl)butanamide).** Tert-butyl (4-((1-(3,4-dichlorophenyl)-1H-pyrazol-3-yl)amino)-4-oxobutyl)carbamate (ref[29]) (41 mg, 100 μmol) was dissolved in DCM (4 mL) and treated with TFA (1 mL). The reaction mixture was stirred at room temperature for 30 min and concentrated under reduced pressure. To the oil residue, bis-PEG3-NHS ester (22.2 mg, 50 μmol) in DMF (2 mL) was added, followed by DIPEA (100 μL). The reaction mixture was stirred at room temperature for 30 min, concentrated under reduced pressure, and purified by HPLC, affording the desired product VS1144 as a white powder (20 mg, 48% yield). $^1$H NMR (400 MHz, $CD_3OD$) δ 8.07 (d, *J* = 2.6 Hz, 2H), 7.86 (d, *J* = 2.5 Hz, 2H), 7.57 (dd, *J* = 8.8, 2.6 Hz, 2H), 7.47 (d, *J* = 8.8 Hz, 2H), 6.77 (d, *J* = 2.6 Hz, 2H), 3.63 (t, *J* = 6.1 Hz, 4H), 3.52 (tt, *J* = 5.4, 2.7 Hz, 9H), 3.19 (t, *J* = 6.8 Hz, 5H), 2.35 (t, *J* = 6.2 Hz, 9H), 1.80 ppm (p, *J* = 7.1 Hz, 4H). $^{13}$C NMR (400 MHz, $CD_3OD$) δ174.15, 173.47, 150.69, 140.68, 134.17, 132.24, 129.97, 129.09, 120.81, 118.39, 102.00, 71.44, 71.34, 68.26, 39.75, 37.73, 34.53, 26.41. ppm. HRMS (ESI-TOF): calculated for $C_{36}H_{43}Cl_4N_8O_7$ (M+H): 841.1974, found 841.1971.

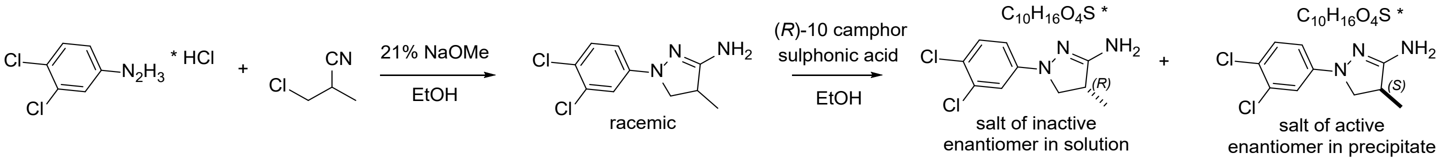


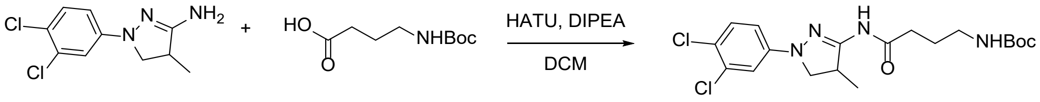


**VS1060: 1-(3,4-dichlorophenyl)-4-methyl-4,5-dihydro-1H-pyrazol-3-amine, separation of enantiomers and linker attachment.** To a suspension of 3,4-dichlorophenylhydrazine hydrochloride (2.1 g, 10 mmol) in EtOH (8 mL) at room temperature, sodium methoxide in MeOH (21 w/w %, 6.4 mL, 24 mmol) was added dropwise. To the mixture, 3-chloro-2-methylpropionitrile (1.1 g, 11 mmol) in 2.5 mL EtOH was added dropwise and the mixture was stirred at 85 °C overnight. The next day, the resulting suspension was evaporated to remove EtOH, water was added, and the yellowish solid was filtered, washed, and dried to provide the desired product (1.75 g, 90% yield). Characterization data of the resulting product matches the data reported for 1-(3,4-dichlorophenyl)-4-methyl-4,5-dihydro-1H-pyrazol-3-amine (ref[31]). The enantiomers of 1-(3,4-dichlorophenyl)-4-methyl-4,5-dihydro-1H-pyrazol-3-amine were separated by recrystallization of diastereomers formed in the reaction with (*R*)-10 camphor sulphonic acid. More specifically, 1-(3,4-dichlorophenyl)-4-methyl-4,5-dihydro-1H-pyrazol-3-amine (1.2 g, 5 mmol) was dissolved in EtOH (12 mL) and treated with (*R*)-10 camphor sulphonic acid (1.2 g, 5 mmol). The mixture was heated at 50 ˚C for 30 min and cooled down to room temperature. After 1h the precipitate was collected via filtration (solid residue contains active (*S*)-enantiomer; absolute stereochemistry assigned based on reported activity data [ref[31]]) and separated from the filtrate (solution contains inactive (*R)*-enantiomers) that is later concentrated under reduced pressure. Salts of separated enantiomers were dissolved in water, neutralized by NaOH, extracted by DCM, washed with brine, and concentrated under reduced pressure affording active and inactive enantiomers of 1-(3,4-dichlorophenyl)-4-methyl-4,5-dihydro-1H-pyrazol-3-amine used in the following steps without further characterization. Coupling of racemic and active/inactive enantiomers 1-(3,4-dichlorophenyl)-4-methyl-4,5-dihydro-1H-pyrazol-3-amine with N-Boc gamma-amino butyric acid was performed using the earlier reported method affording racemic and active/inactive enantiomers of tert-butyl (4-((1-(3,4-dichlorophenyl)-4-methyl-4,5-dihydro-1H-pyrazol-3-yl)amino)-4-oxobutyl)carbamate used in the next steps (ref[30]).

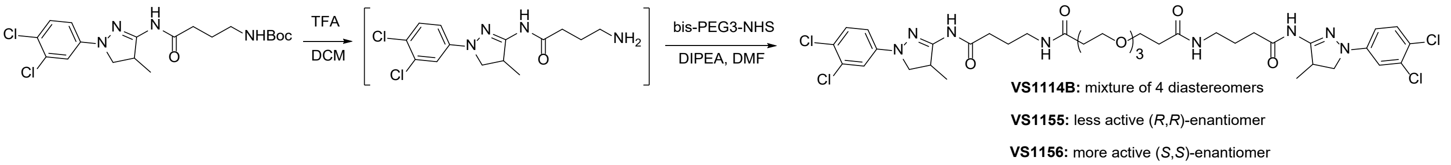


**Compound 6 (VS1114B): 4,4'-((3,3'-((oxybis(ethane-2,1-diyl))bis(oxy))bis(propanoyl))bis(azanediyl)) bis(N-(1-(3,4-dichlorophenyl)-4-methyl-4,5-dihydro-1H-pyrazol-3-yl)butanamide).** Racemic tert-butyl (4-((1-(3,4-dichlorophenyl)-4-methyl-4,5-dihydro-1H-pyrazol-3-yl)amino)-4-oxobutyl)carbamate (42.9 mg, 100 μmol) was dissolved in DCM (4 mL) and treated with TFA (1 mL). The reaction mixture was stirred at room temperature for 30 min and concentrated under reduced pressure. To the oil residue, bis-PEG3-NHS ester (22.2 mg, 50 μmol) in DMF (2 mL) was added, followed by DIPEA (100 μL). The reaction mixture was stirred at room temperature for 30 min, concentrated under reduced pressure, and purified by HPLC, affording the desired product VS1144 as a white powder (25 mg, 57% yield). $^1$H NMR (400 MHz, $CDCl_3$ + $CD_3OD$ (1:1)) δ 7.78 (t, J = 5.8 Hz, 2H), 7.22 (d, J = 8.8 Hz, 2H), 7.02 (d, J = 2.6 Hz, 2H), 6.75 (s, 2H), 3.87 (s, 2H), 3.77 – 3.66 (m, 6H), 3.61 (s, 8H), 3.53 (dd, J = 9.5, 4.3 Hz, 2H), 3.24 (q, J = 6.5 Hz, 4H), 2.44 (t, J = 6.1 Hz, 4H), 2.37 (t, J = 7.5 Hz, 4H), 1.83 (p, J = 7.1 Hz, 4H), 1.25 (d, J = 6.9 Hz, 6H). 13C NMR (101 MHz, $CDCl_3$ + $CD_3OD$ (1:1)) δ 173.43, 172.75, 153.61, 147.29, 133.06, 130.97, 121.42, 114.78, 112.87, 70.90, 70.77, 67.80, 56.75, 39.97, 39.19, 37.25, 34.19, 25.80, 17.71. HRMS (ESI-TOF): calculated for $C_{38}H_{51}Cl_4N_8O_7$ (M+H): 873.2600, found: 873.2598.

**Compound 7 (VS1155): (*R*,*R*)-4,4'-((3,3'-((oxybis(ethane-2,1-diyl))bis(oxy))bis(propanoyl))bis(azanediyl)) bis(N-(1-(3,4-dichlorophenyl)-4-methyl-4,5-dihydro-1H-pyrazol-3-yl)butanamide).** Using **the** inactive enantiomer of tert-butyl (4-((1-(3,4-dichlorophenyl)-4-methyl-4,5-dihydro-1H-pyrazol-3-yl)amino)-4-oxobutyl)carbamate and following the procedure for VS1114B, the desired product VS1155 was obtained (27 mg, 62%). $^1$H and $^{13}$C NMR data matched that reported for VS1114B. HRMS (ESI-TOF): calculated for $C_{38}H_{51}Cl_4N_8O_7$ (M+H): 873.2600, found: 873.2604.

**Compound 8 (VS1156): (*S*,*S*)-4,4'-((3,3'-((oxybis(ethane-2,1-diyl))bis(oxy))bis(propanoyl))bis(azanediyl)) bis(N-(1-(3,4-dichlorophenyl)-4-methyl-4,5-dihydro-1H-pyrazol-3-yl)butanamide).** Using the active enantiomer of tert-butyl (4-((1-(3,4-dichlorophenyl)-4-methyl-4,5-dihydro-1H-pyrazol-3-yl)amino)-4-oxobutyl)carbamate and following the procedure for VS1114B, the desired product VS1156 was obtained (25 mg, 57%). $^1$H and $^{13}$C NMR data matched that reported for VS1114B. HRMS (ESI-TOF): calculated for $C_{38}H_{51}Cl_4N_8O_7$ (M+H): 873.2600, found: 873.2604.

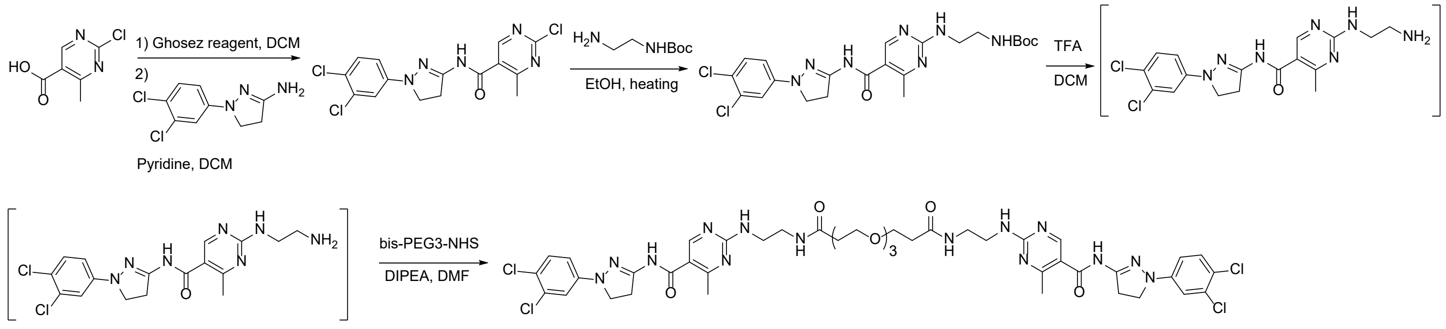


**VS1065: 2-chloro-N-(1-(3,4-dichlorophenyl)-4,5-dihydro-1H-pyrazol-3-yl)-4-methylpyrimidine-5-carboxamide.** 2-chloro-4-methylpyrimidine-5-carboxylic acid (120 mg, 0.7 mmol) was dissolved in DCM (6 mL) and treated with Ghosez reagent (180 µL, 1.4 mmol). The reaction mixture was stirred at room temperature for 1 hour, and 1-(3,4-dichlorophenyl)-4,5-dihydro-1H-pyrazol-3-amine (138 mg, 0.6 mmol) was added, followed by pyridine (180 µL). The reaction mixture was stirred at room temperature overnight, quenched with an aqueous $NaHCO_3$ solution, extracted with DCM, and concentrated under reduced pressure. The residue was purified by flash column chromatography (DCM: EtOAc gradient from 100:0 to 70:30), affording the desired product as a yellowish solid (45 mg, 21% yield). $^1$H NMR (400 MHz, $CDCl_3$ + $CD_3OD$ (1:1)) δ 8.67 (s, 1H), 7.24 (d, J = 8.8 Hz, 1H), 7.05 (d, J = 2.7 Hz, 1H), 6.77 (dd, J = 8.9, 2.7 Hz, 1H), 3.81 (t, J = 10.0 Hz, 2H), 3.56 (t, J = 10.0 Hz, 2H), 2.67 ppm (s, 3H). $^{13}$C NMR (101 MHz, $CDCl_3$ + $CD_3OD$ (1:1)) δ 170.66, 164.35, 161.94, 158.35, 149.44, 147.01, 133.16, 131.02, 128.25, 121.72, 114.79, 112.86, 49.31, 33.30, 22.89 ppm. HRMS (ESI-TOF): calculated for $C_{15}H_{13}Cl_3N_5O$ (M+H): 386.0151, found: 386.0148.

**VS1067: tert-butyl (2-((5-((1-(3,4-dichlorophenyl)-4,5-dihydro-1H-pyrazol-3-yl)carbamoyl)-4-methylpyrimidin-2-yl)amino)ethyl)carbamate.** 2-chloro-N-(1-(3,4-dichlorophenyl)-4,5-dihydro-1H-pyrazol-3-yl)-4-methylpyrimidine-5-carboxamide (VS1065, 22 mg, 60 µmol) and tert-butyl (2-aminoethyl)carbamate were dissolved in ethanol (0.5 mL) and heated at 70 ºC overnight. The next day, the solvent was removed under reduced pressure, and the residue was purified via flash column chromatography (DCM: MeOH gradient from 100:0 up to 95:5), affording the desired product as a yellowish solid (22 mg, 73% yield). $^1$H NMR (400 MHz, $CD_3OD$) δ 8.43 (s, 1H), 7.27 (d, J = 8.9 Hz, 1H), 7.12 (d, J = 2.7 Hz, 1H), 6.83 (dd, J = 8.9, 2.7 Hz, 1H), 3.78 (t, J = 10.1 Hz, 2H), 3.62 – 3.43 (m, 4H), 3.27 (t, J = 6.1 Hz, 2H), 2.51 (s, 3H), 1.43 (s, 9H). $^{13}$C NMR (101 MHz, $CD_3OD$) δ 167.20, 163.47, 158.89, 158.65, 151.19, 148.15, 133.43, 131.44, 121.41, 118.46, 115.07, 113.36, 80.14, 49.49, 42.29, 40.99, 33.77, 28.75, 23.55. HRMS (ESI-TOF): calculated for $C_{22}H_{28}Cl_2N_7O_3$ (M+H): 508.1625, found: 508.1628.

**Compound 9 (VS1142): 2,2'-((4,16-dioxo-7,10,13-trioxa-3,17-diazanonadecane-1,19-diyl)bis(azanediyl)) bis(N-(1-(3,4-dichlorophenyl)-4,5-dihydro-1H-pyrazol-3-yl)-4-methylpyrimidine-5-carboxamide).** Tert-butyl (2-((5-((1-(3,4-dichlorophenyl)-4,5-dihydro-1H-pyrazol-3-yl)carbamoyl)-4-methylpyrimidin-2-

yl)amino)ethyl)carbamate (VS1067) (22 mg, 43 µmol) was dissolved in DCM (1 mL) and treated with TFA (0.25 mL). The reaction mixture was stirred at room temperature for 30 min and concentrated under reduced pressure. To the oil residue, bis-PEG3-NHS ester (9.5 mg, 22 µmol) in DMF (1 mL) was added, followed by DIPEA (50 µL). The reaction mixture was stirred at room temperature for 30 min, concentrated under reduced pressure, and purified by HPLC, affording the desired product VS1142 as a white powder (10 mg, 23% yield). $^{1}H$ NMR (400 MHz, $CDCl_3$ + $CD_3OD$ (1:1)) δ 8.41 (d, J = 4.5 Hz, 3H), 7.59 (s, 3H), 7.24 (d, J = 8.8 Hz, 2H), 7.06 (d, J = 2.7 Hz, 2H), 6.77 (dd, J = 8.8, 2.7 Hz, 2H), 3.78 (t, J = 9.9 Hz, 4H), 3.71 (t, J = 6.1 Hz, 4H), 3.61 (hept, J = 2.8 Hz, 8H), 3.58 – 3.50 (m, 8H), 3.44 (dd, J = 6.7, 5.1 Hz, 4H), 2.52 (s, 6H), 2.44 ppm (t, J = 6.1 Hz, 4H). $^{13}C$ NMR (400 MHz, $CDCl_3$ + $CD_3OD$ (1:1)) δ 173.64, 166.41, 162.67, 158.25, 150.38, 147.26, 133.11, 130.98, 121.44, 114.75, 112.81, 70.85, 70.76, 67.72, 41.41, 39.65, 37.26, 33.39 ppm. HRMS (ESI-TOF): calculated for $C_{44}H_{53}Cl_4N_{14}O_7$ (M+H): 1031.2941, found: 1031.2946.

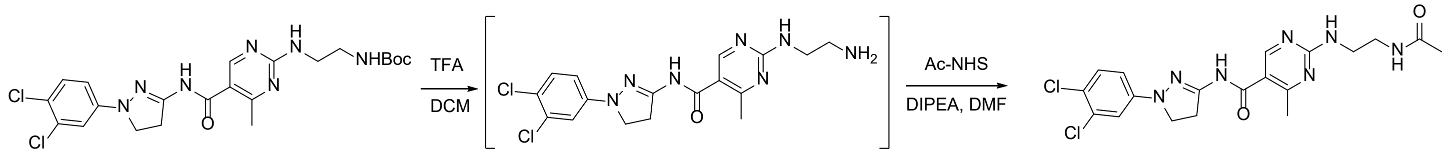

**BRD6501: 2-((2-acetamidoethyl)amino)-N-(1-(3,4-dichlorophenyl)-4,5-dihydro-1H-pyrazol-3-yl)-4-methylpyrimidine-5-carboxamide.** Tert-butyl (2-((5-((1-(3,4-dichlorophenyl)-4,5-dihydro-1H-pyrazol-3-yl)carbamoyl)-4-methylpyrimidin-2-yl)amino) ethyl)carbamate (VS1067) (22 mg, 43 µmol) was dissolved in DCM (1 mL) and treated with TFA (0.25 mL). The reaction mixture was stirred at room temperature for 30 min and concentrated under reduced pressure. To the oil residue, acetic acid N-hydroxysuccinimide ester (7.9 mg, 50 µmol) in DMF (1 mL) was added, followed by DIPEA (50 µL). The reaction mixture was stirred at room temperature for 30 min, concentrated under reduced pressure, and purified by HPLC, affording the desired product VS1148 as a white powder (12 mg, 62% yield). $^{1}H$ NMR (400 MHz, $CDCl_3$ + $CD_3OD$ (1:1)) δ 8.41 (s, 1H), 7.61 (s, 1H), 7.23 (d, J = 8.8 Hz, 1H), 7.06 (d, J = 2.6 Hz, 1H), 6.77 (dd, J = 8.9, 2.6 Hz, 1H), 3.77 (t, J = 9.9 Hz, 2H), 3.60 – 3.46 (m, 4H), 3.39 (t, J = 6.0 Hz, 2H), 2.51 (s, 3H), 1.93 ppm (s, 3H). $^{13}C$ NMR (400 MHz, $CDCl_3$ + $CD_3OD$ (1:1)) δ 173.16, 166.51, 162.67, 158.28, 150.42, 147.32, 133.12, 131.00, 121.41, 114.76, 112.86, 41.38, 39.80, 33.42, 22.67. HRMS (ESI-TOF): calculated for $C_{19}H_{22}Cl_2N_7O_2$ (M+H): 450.1207, found: 450.1207.

**Synthesis of compounds with different linker lengths (PEG0–3):**

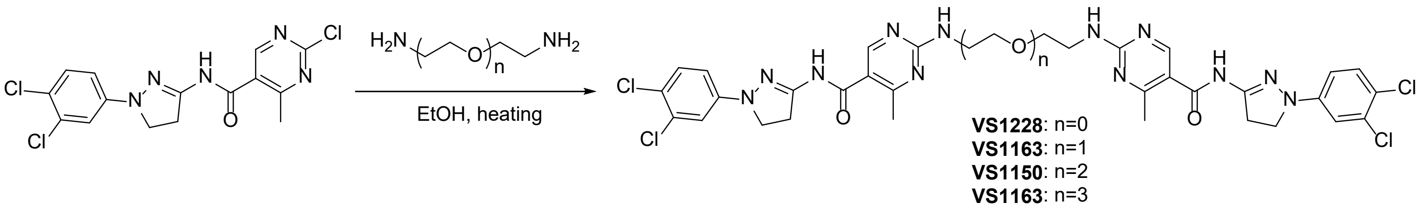

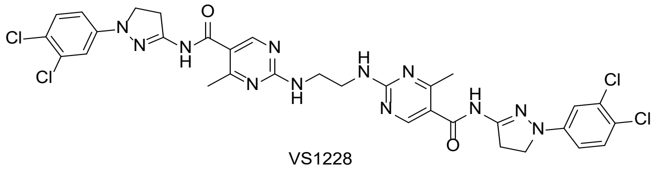

VS1228

**2,2'-(ethane-1,2-diylbis(azanediyl))bis(N-(1-(3,4-dichlorophenyl)-4,5-dihydro-1H-pyrazol-3-yl)-4-methylpyrimidine-5-carboxamide) (VS1228).** 2-chloro-N-(1-(3,4-dichlorophenyl)-4,5-dihydro-1H-pyrazol-3-yl)-4-methylpyrimidine-5-carboxamide (VS1065, 20 mg, 52 µmol) and ethylenediamine (2 mg, 33 µmol) were dissolved in 1 mL EtOH, and DIPEA was added (20 µL). The reaction was heated at 70 ºC overnight. The solvent was removed under reduced pressure, and the oil residue was purified via HPLC, affording the desired product VS1228 as a white solid (6.6 mg, 35% yield). $^1$H NMR (400 MHz, DMSO-*d₆*) δ 10.97 (s, 2H), 8.56 – 8.38 (m, 2H), 7.80 – 7.61 (m, 2H), 7.41 (d, *J* = 8.8 Hz, 2H), 7.08 (d, *J* = 2.6 Hz, 2H), 6.85 (dd, *J* = 9.0, 2.7 Hz, 2H), 3.75 (t, *J* = 9.8 Hz, 4H), 3.60 – 3.39 (m, 8H), 2.42 (s, 6H). $^{13}$C NMR (101 MHz, DMSO-$d_6$) δ 167.54, 164.89, 161.78, 158.09, 150.97, 146.55, 131.33, 130.66, 118.41, 116.22, 113.21, 112.35, 47.92, 32.75, 23.56, 23.06. HRMS (ESI-TOF): calculated for $C_{32}H_{30}Cl_4N_{12}O_2$ (M+H): 755.1442, found: 757.1404.

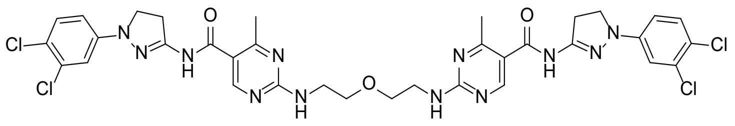

VS1163

**2,2'-((oxybis(ethane-2,1-diyl))bis(azanediyl))bis(N-(1-(3,4-dichlorophenyl)-4,5-dihydro-1H-pyrazol-3-yl)-4-methylpyrimidine-5-carboxamide) (VS1163).** 2-chloro-N-(1-(3,4-dichlorophenyl)-4,5-dihydro-1H-pyrazol-3-yl)-4-methylpyrimidine-5-carboxamide (VS1065, 20 mg, 52 μmol) and 2,2'-oxybis(ethan-1-amine) (2.7 mg, 26 μmol) were dissolved in 1 mL EtOH, and DIPEA was added (20 μL). The reaction was heated at 70 ºC overnight. The solvent was removed under reduced pressure and oil residue was purified via HPLC, affording the desired product VS1163 as a yellowish solid (8 mg, 38% yield). $^1$H NMR (400 MHz, DMSO-$d_6$) δ 10.95 (s, 2H), 8.47 (s, 2H), 7.70 (s, 2H), 7.41 (d, J = 8.9 Hz, 2H), 7.08 (d, J = 2.6 Hz, 2H), 6.84 (dd, J = 8.9, 2.7 Hz, 2H), 3.75 (t, J = 9.8 Hz, 4H), 3.63 – 3.42 (m, 12H), 2.43 (s, 6H). $^{13}$C NMR (101 MHz, DMSO-$d_6$) δ 165.38, 162.24, 158.73, 151.43, 147.03, 131.82, 131.12, 118.90, 113.69, 112.82, 69.10, 55.38, 48.39, 33.23, 31.16 HRMS (ESI-TOF): calculated for $C_{34}H_{35}Cl_4N_{12}O_3$ (M+H): 801.1674, found: 801.1676.

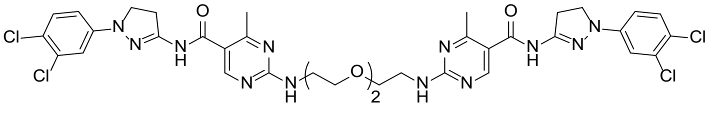

VS1150

**BRD8833 (VS1150): 2,2'-(((ethane-1,2-diylbis(oxy))bis(ethane-2,1-diyl))bis(azanediyl))bis(N-(1-(3,4-dichlorophenyl)-4,5-dihydro-1H-pyrazol-3-yl)-4-methylpyrimidine-5-carboxamide).** 2-chloro-N-(1-(3,4-dichlorophenyl)-4,5-dihydro-1H-pyrazol-3-yl)-4-methylpyrimidine-5-carboxamide (VS1065, 40 mg, 104 µmol) and 2,2'-(ethane-1,2-diylbis(oxy))bis(ethan-1-amine) (7.5 mg, 51 µmol) were dissolved in 1 mL EtOH, and DIPEA was added (20 µL). The reaction was heated at 70 ºC overnight. The solvent was removed under reduced pressure, and the oil residue was purified via HPLC, affording the desired product VS1150 as a yellowish solid (20 mg, 44% yield). $^{1}$H NMR (400 MHz, DMSO-$d_6$) δ 8.46 (d, J = 15.5 Hz, 2H), 7.63 (d, J = 18.0 Hz, 2H), 7.41 (d, J = 8.8 Hz, 2H), 7.08 (d, J = 2.7 Hz, 2H), 6.84 (dd, J = 8.9, 2.7 Hz, 2H), 3.75 (t, J = 9.8 Hz, 4H), 3.63 – 3.39 (m, 16H), 2.51 (p, J = 1.8 Hz, 6H), 2.43 ppm (d, J = 11.0 Hz, 4H). $^{13}$C NMR (101 MHz, DMSO-$d_6$) δ 167.61, 164.91, 161.74, 158.14, 151.01, 146.55, 131.36, 130.68, 118.42, 116.43, 113.22, 112.36, 69.61, 68.77, 47.93, 40.40, 32.78, 30.74 ppm. HRMS (ESI-TOF): calculated for $C_{36}H_{39}Cl_4N_{12}O_4$ (M+H): 845.1936, found: 845.1937.

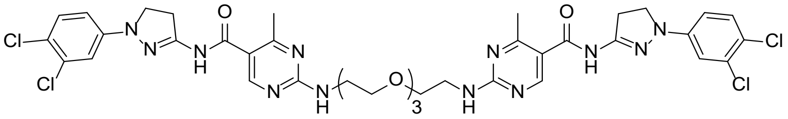

VS1186

**2,2'-((((oxybis(ethane-2,1-diyl))bis(oxy))bis(ethane-2,1-diyl))bis(azanediyl))bis(N-(1-(3,4-dichlorophenyl)-4,5-dihydro-1H-pyrazol-3-yl)-4-methylpyrimidine-5-carboxamide) (VS1186).** 2-chloro-N-(1-(3,4-dichlorophenyl)-4,5-dihydro-1H-pyrazol-3-yl)-4-methylpyrimidine-5-carboxamide (VS1065, 15 mg, 39 µmol) and 2,2'-((oxybis(ethane-2,1-diyl))bis(oxy))bis(ethan-1-amine) (3.8 mg, 20 µmol) were dissolved in 1 mL EtOH, and DIPEA was added (20 µL). The reaction was heated at 70 ºC overnight. The solvent was removed under reduced pressure and the oil residue was purified via HPLC, affording the desired product VS1186 as a yellowish solid (12 mg, 68% yield). $^{1}$H NMR (400 MHz, DMSO-$d_6$) δ 8.47 (s, 2H), 7.59 (s, 2H), 7.40 (d, $J$ = 8.9 Hz, 2H), 7.17 – 7.00 (m, 2H), 6.84 (d, $J$ = 9.0 Hz, 2H), 3.75 (t, $J$ = 9.9 Hz, 4H), 3.51 (m, 20H), 2.43 (s, 4H), 2.08 (s, 6H). $^{13}$C NMR (101 MHz, DMSO-d6) $^{13}$C NMR (101 MHz, DMSO-$d_6$) δ 167.54, 164.88, 161.72, 158.15, 150.93, 146.53, 131.32, 130.62, 118.40, 118.04, 113.20, 112.32, 69.76, 69.59, 68.72, 54.89, 47.89, 40.38, 40.15, 39.94, 39.73, 39.52, 39.31, 39.10, 38.89, 32.73. HRMS (ESI-TOF): calculated for $C_{38}H_{43}Cl_4N_{12}O_5$ (M+H): 889.2198, found: 889.2198.

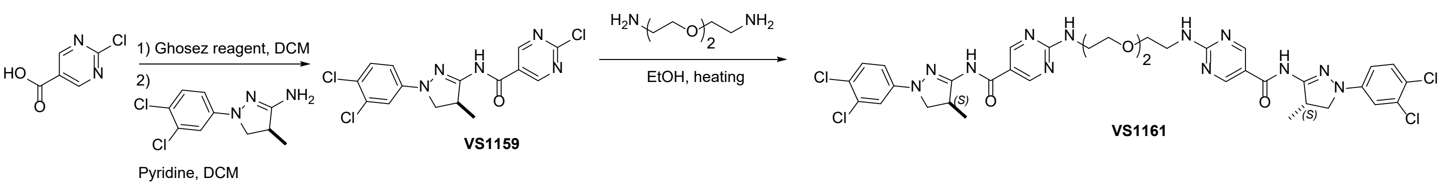


**(*S*)-2-chloro-N-(1-(3,4-dichlorophenyl)-4-methyl-4,5-dihydro-1H-pyrazol-3-yl)pyrimidine-5-carboxamide (VS1159).** 2-chloropyrimidine-5-carboxylic acid (180 mg, 1.1 mmol) was dissolved in DCM (10 mL) and treated with Ghosez reagent (180 µL, 2.1 mmol). The reaction mixture was stirred at room temperature for 1 hour and (*S*)-1-(3,4-dichlorophenyl)-4-methyl-4,5-dihydro-1H-pyrazol-3-amine (VS1060) (225 mg, 0.9 mmol) was added, followed by pyridine (270 µL). The reaction mixture was stirred at room temperature overnight, quenched with an aqueous $NaHCO_3$ solution, extracted with DCM, and concentrated under reduced pressure. The residue was purified by flash column chromatography (DCM: EtOAc gradient from 100:0 to 70:30) affording the desired product as a yellowish solid (14 mg, 4.1% yield). $^{1}$H NMR (400 MHz, $CDCl_3$ + $CD_3OD$ (1:1)) δ 9.14 (s, 2H), 7.27 (d, J = 8.8 Hz, 1H), 7.09 (d, J = 2.6 Hz, 1H), 6.81 (dd, J = 8.8, 2.7 Hz, 1H), 4.08 (dtd, J = 14.2, 7.0, 3.5 Hz, 1H), 3.83 (t, J = 10.0 Hz, 1H), 3.65 (dd, J = 9.5, 4.5 Hz, 1H), 1.34 ppm (d, J = 7.0 Hz, 3H). $^{13}$C NMR (101 MHz, $CDCl_3$ + $CD_3OD$ (1:1)) δ 164.01, 161.75, 159.89, 153.02, 146.72, 133.12, 130.97, 127.01, 121.82, 114.79, 112.82, 56.90, 39.91, 17.78. HRMS (ESI-TOF): calculated for $C_{15}H_{13}Cl_3N_5O$ (M+H): 386.0151, found: 386.0145.

**BRD8834 (VS1161): 2,2'-(((ethane-1,2-diylbis(oxy))bis(ethane-2,1-diyl))bis(azanediyl))bis(N-((S)-1-(3,4-dichlorophenyl)-4-methyl-4,5-dihydro-1H-pyrazol-3-yl)pyrimidine-5-carboxamide).** (*S*)-2-chloro-N-(1-(3,4-dichlorophenyl)-4-methyl-4,5-dihydro-1H-pyrazol-3-yl)pyrimidine-5-carboxamide (VS1159, 12 mg, 31 µmol) and 2,2'-(ethane-1,2-diylbis(oxy))bis(ethan-1-amine) (2 mg, 14 µmol) were dissolved in 0.5 mL EtOH, and DIPEA was added (10 µL). The reaction was heated at 70 ºC overnight. The solvent was removed under reduced pressure, and the oil residue was purified via HPLC, affording the desired product VS1161 as a yellowish solid (7 mg, 59% yield). $^{1}$H NMR (400 MHz, $CDCl_3$ + $CD_3OD$ (1:1)) δ 8.81 (s, 4H), 7.52 (s, 5H), 7.25 (d, *J* = 8.8 Hz, 2H), 7.06 (d, *J* = 2.7 Hz, 2H), 6.78 (dd, *J* = 8.9, 2.7 Hz, 2H), 3.84 – 3.74 (m, 2H), 3.72 – 3.65 (m, 8H), 3.59 (dd, *J* = 9.5, 4.6 Hz, 2H), 3.33 (s, 8H), 1.31 (d, *J* = 7.0 Hz, 6H). $^{13}$C NMR (101 MHz, DMSO-$d_6$) δ 163.17, 162.91, 158.94, 154.37, 146.75, 131.73, 131.00, 119.22, 115.74, 113.81, 112.86, 69.92, 68.94, 55.77, 40.92, 17.35. HRMS (ESI-TOF): calculated for $C_{36}H_{39}Cl_4N_{12}O_4$ (M+H): 845.1936, found: 845.1935.

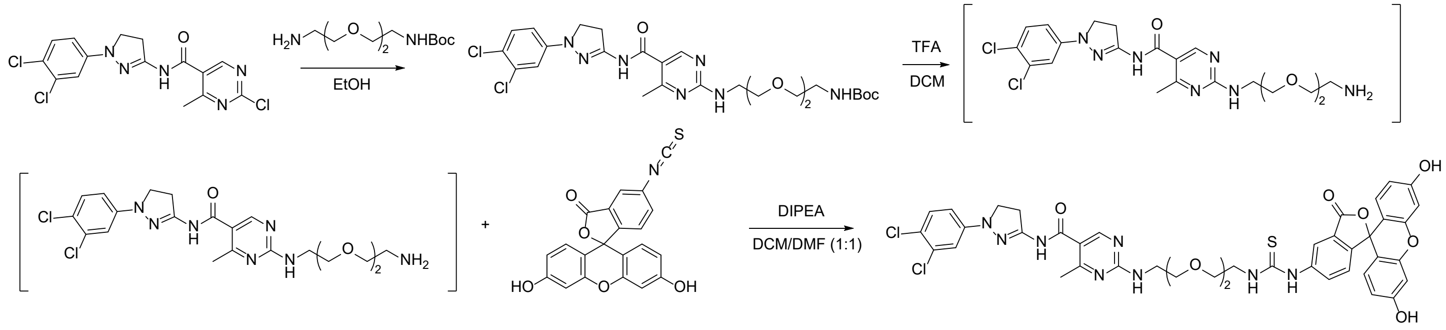

**TR-FRET tracer (VS1231): N-(1-(3,4-dichlorophenyl)-4,5-dihydro-1H-pyrazol-3-yl)-2-((2-(2-(2-(3-(3',6'-dihydroxy-3-oxo-3H-spiro[isobenzofuran-1,9'-xanthen]-5-yl)thioureido)ethoxy)ethoxy)ethyl)amino) pyrimidine-5-carboxamide:** Tert-butyl (2-(2-(2-aminoethoxy)ethoxy)ethyl)carbamate (30 mg, 0.12 mmol) is added to 2-chloro-N-(1-(3,4-dichlorophenyl)-4,5-dihydro-1H-pyrazol-3-yl)pyrimidine-5-carboxamide (VS1065) (45 mg, 0.12 mmol) in 1 mL EtOH. The reaction was heated at 70ºC overnight, and the solvent was removed under reduced pressure. The resulting oil was purified by flash chromatography (DCM:MeOH). The resulting tert-butyl (2-(2-((5-((1-(3,4-dichlorophenyl)-4,5-dihydro-1H-pyrazol-3-yl)carbamoyl)pyrimidin-2-yl)amino)ethoxy) -3-methoxypropyl)carbamate (40 mg, 0.07 mmol) was dissolved in 3 mL DCM, and 0.75 mL TFA was added. After 30 min, the reaction mixture was concentrated under reduced pressure, and the residue was dissolved in 1 mL of a 1:1 mixture of DCM and DMF. To this, 3',6'-dihydroxy-5-isothiocyanato-3H-spiro[isobenzofuran-1,9'-xanthen]-3-one (FITC-isothiocyanate) (30 mg, 0.077 mmol) and 100 µL of DIPEA were added. The reaction was stirred for 1 h at room temperature, the solvent removed under reduced pressure, and the residue purified via HPLC, affording the desired product VS1231 as an orange solid (8.3 mg, 8% yield). $^{1}$H NMR (400 MHz, $CDCl_3$ + $CD_3OD$ (1:1)) δ 8.43 (s, 1H), 8.12 (d, *J* = 2.0 Hz, 1H), 7.88 (d, *J* = 8.6 Hz, 1H), 7.25 (d, *J* = 8.8 Hz, 1H), 7.16 (d, *J* = 8.3 Hz, 1H), 7.07 (d, *J* = 2.7 Hz, 1H), 6.79 (dd, *J* = 8.9, 2.7 Hz, 1H), 6.72 – 6.63 (m, 4H), 6.54 (dd, *J* = 8.7, 2.4 Hz, 2H), 3.88 (s, 1H), 3.83 – 3.73 (m, 5H), 3.73 – 3.60 (m, 8H), 3.55 (t, *J* = 9.9 Hz, 2H), 2.53 (s, 3H). $^{13}$C NMR (101 MHz, $CDCl_3$ + $CD_3OD$ (1:1)) δ 181.43, 169.91, 168.90, 165.60, 161.69, 159.83, 157.53, 152.91, 149.59, 146.51, 140.90, 132.35, 130.94, 129.06, 127.66, 124.53, 120.64, 118.60, 117.24, 116.69, 113.97, 112.47, 112.10, 110.15, 102.57, 70.14, 70.08, 69.39, 44.23, 40.75, 39.68, 32.64, 29.48, 22.82. HRMS (ESI-TOF): calculated for $C_{42}H_{38}Cl_2N_8O_8S$ (M+H): 885.1982, found: 885.1969.

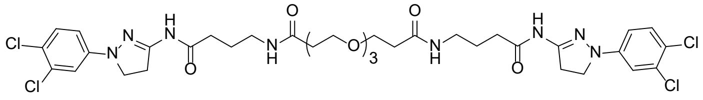


Compound 2 (VS1143)

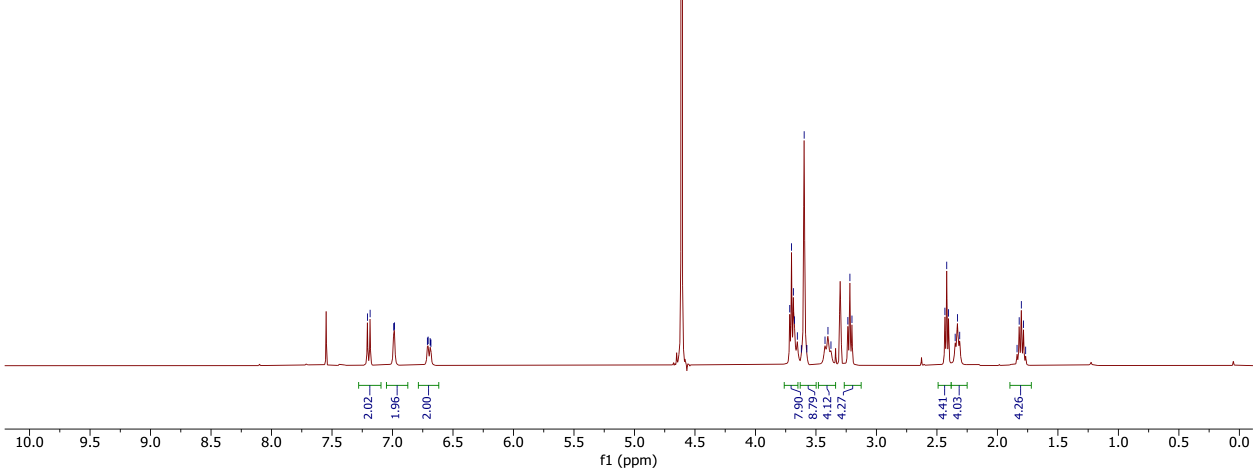


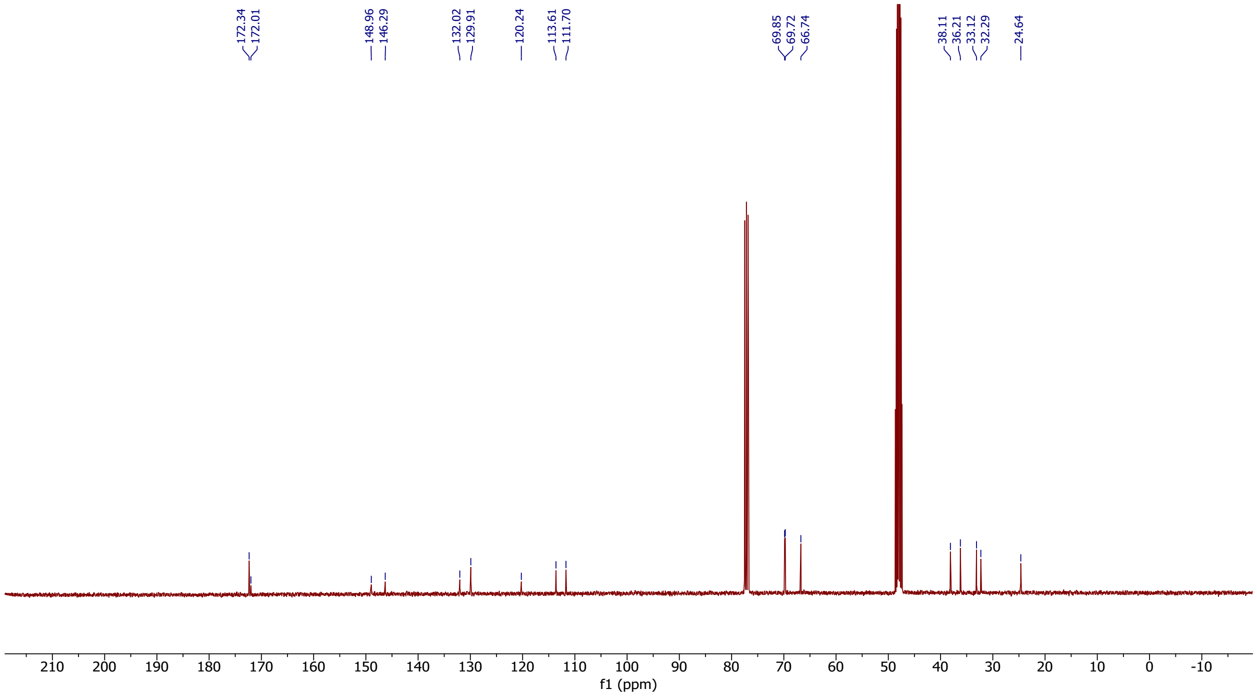

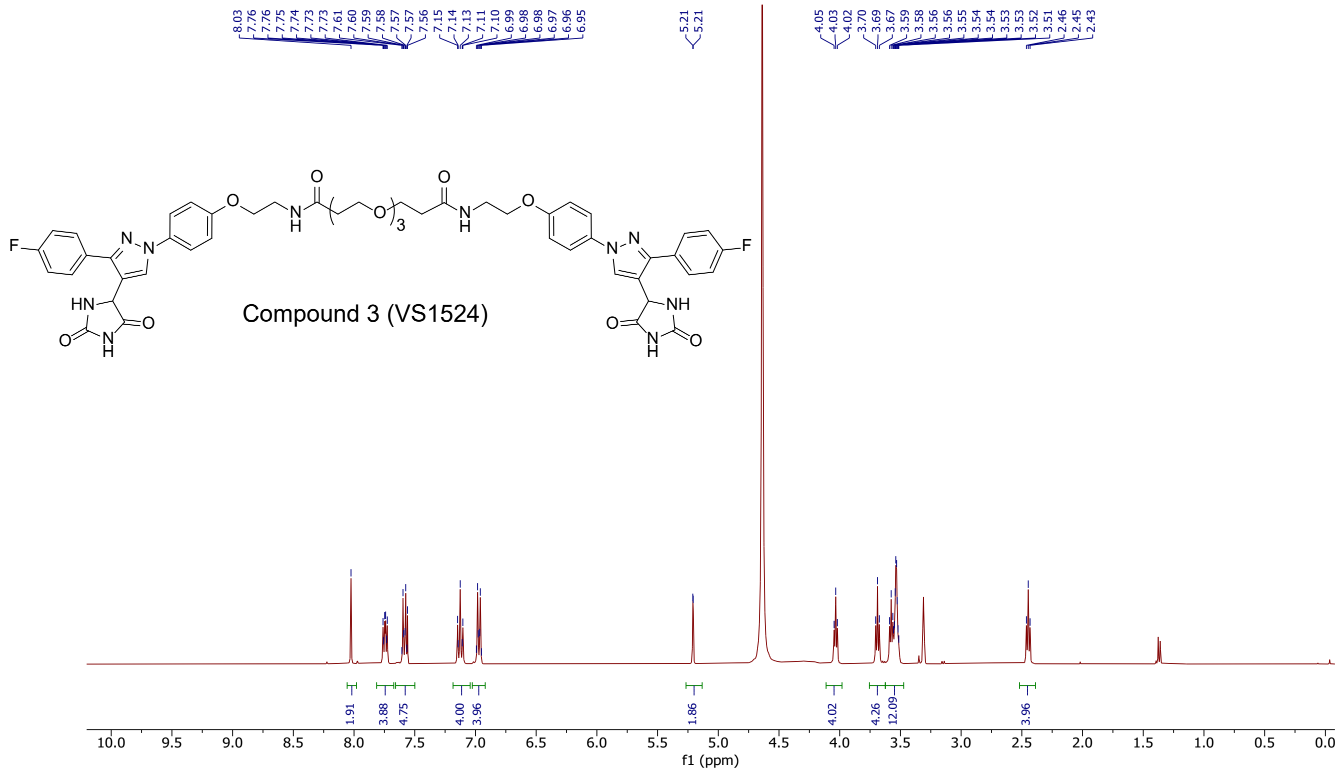
8.03 7.76 7.76 7.75 7.74 7.73 7.73 7.61 7.60 7.59 7.58 7.57 7.57 7.56 7.15 7.14 7.13 7.11 7.10 6.99 6.98 6.98 6.97 6.96 6.95
5.21 5.21
4.05 4.03 4.02 3.70 3.69 3.67 3.59 3.58 3.56 3.56 3.55 3.54 3.54 3.53 3.53 3.52 3.51 2.46 2.45 2.43
Compound 3 (VS1524)
1.91 3.88 4.75 4.00 3.96 1.86 4.02 4.26 12.09 3.96
10.0 9.5 9.0 8.5 8.0 7.5 7.0 6.5 6.0 5.5 5.0 4.5 4.0 3.5 3.0 2.5 2.0 1.5 1.0 0.5 0.0
f1 (ppm)

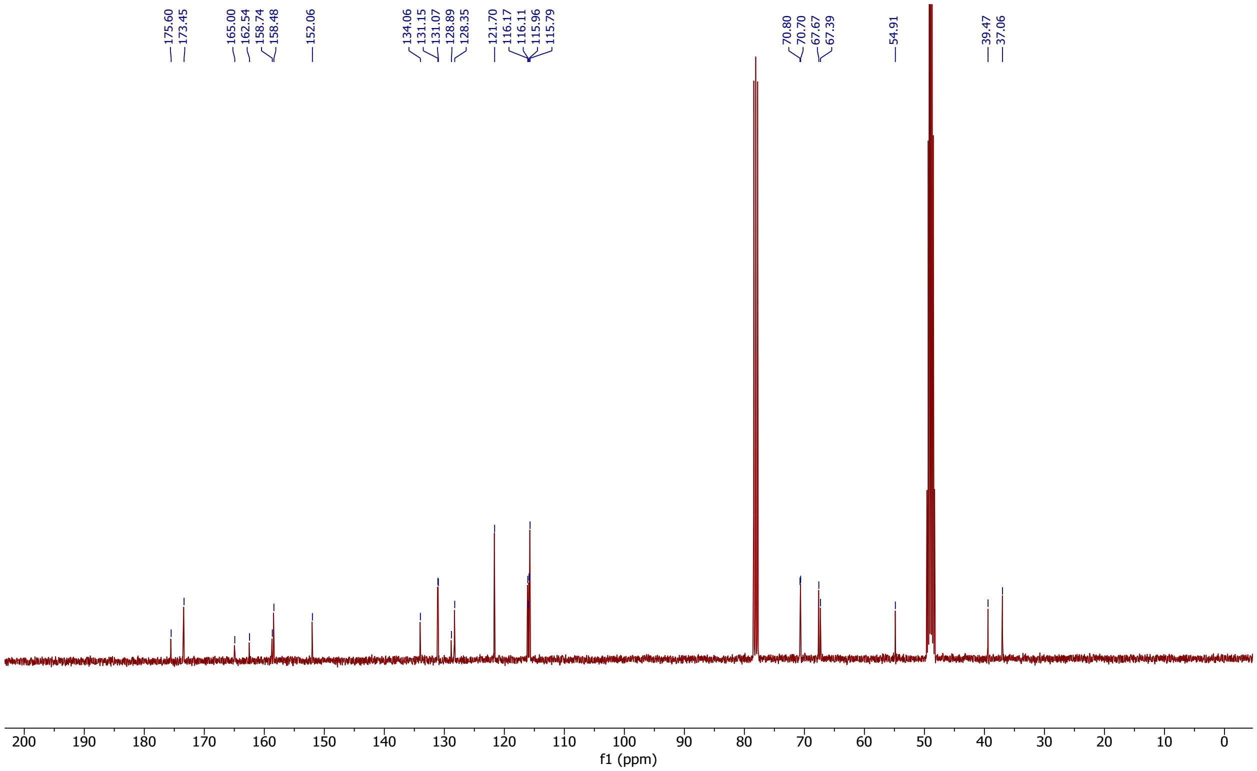
175.60 173.45
165.00 162.54 158.74 158.48
152.06
134.06 131.15 131.07 128.89 128.35
121.70 116.17 116.11 115.96 115.79
70.80 70.70 67.67 67.39
54.91
39.47 37.06
200 190 180 170 160 150 140 130 120 110 100 90 80 70 60 50 40 30 20 10 0
f1 (ppm)

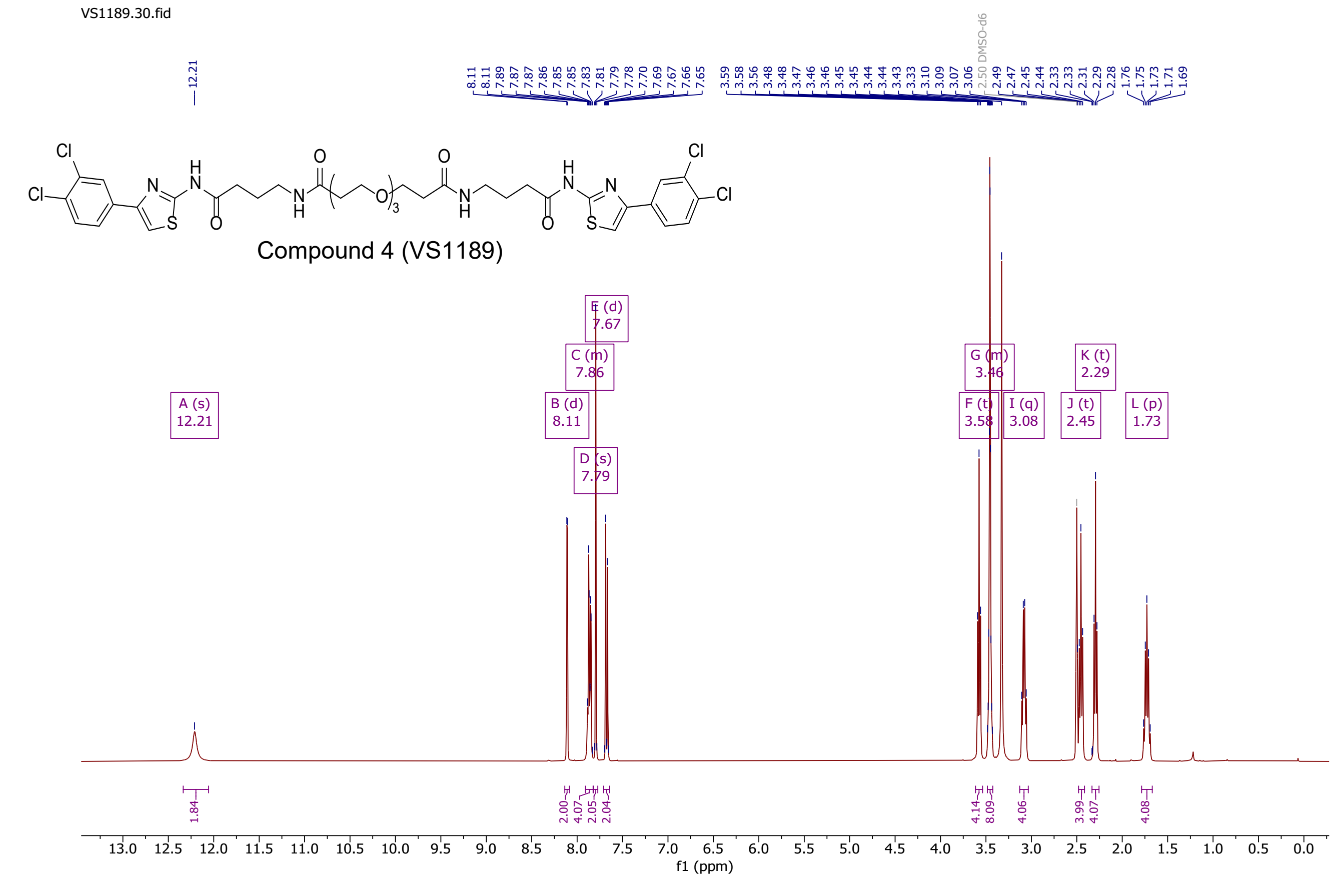
VS1189.30.fid
12.21
8.11 8.11 7.89 7.87 7.87 7.86 7.85 7.85 7.83 7.81 7.79 7.78 7.70 7.69 7.67 7.66 7.65
3.59 3.58 3.56 3.48 3.48 3.47 3.46 3.46 3.45 3.45 3.44 3.44 3.43 3.33 3.10 3.09 3.07 3.06 2.50 DMSO-d6 2.49 2.47 2.45 2.44 2.33 2.31 2.29 2.28 1.76 1.75 1.73 1.71 1.69
Compound 4 (VS1189)
A (s) 12.21
B (d) 8.11
C (m) 7.86
D (s) 7.79
E (d) 7.67
F (t) 3.58
G (m) 3.46
I (q) 3.08
J (t) 2.45
K (t) 2.29
L (p) 1.73
1.84
2.00 4.07 2.05 2.04
4.14 8.09 4.06 3.99 4.07 4.08
f1 (ppm)

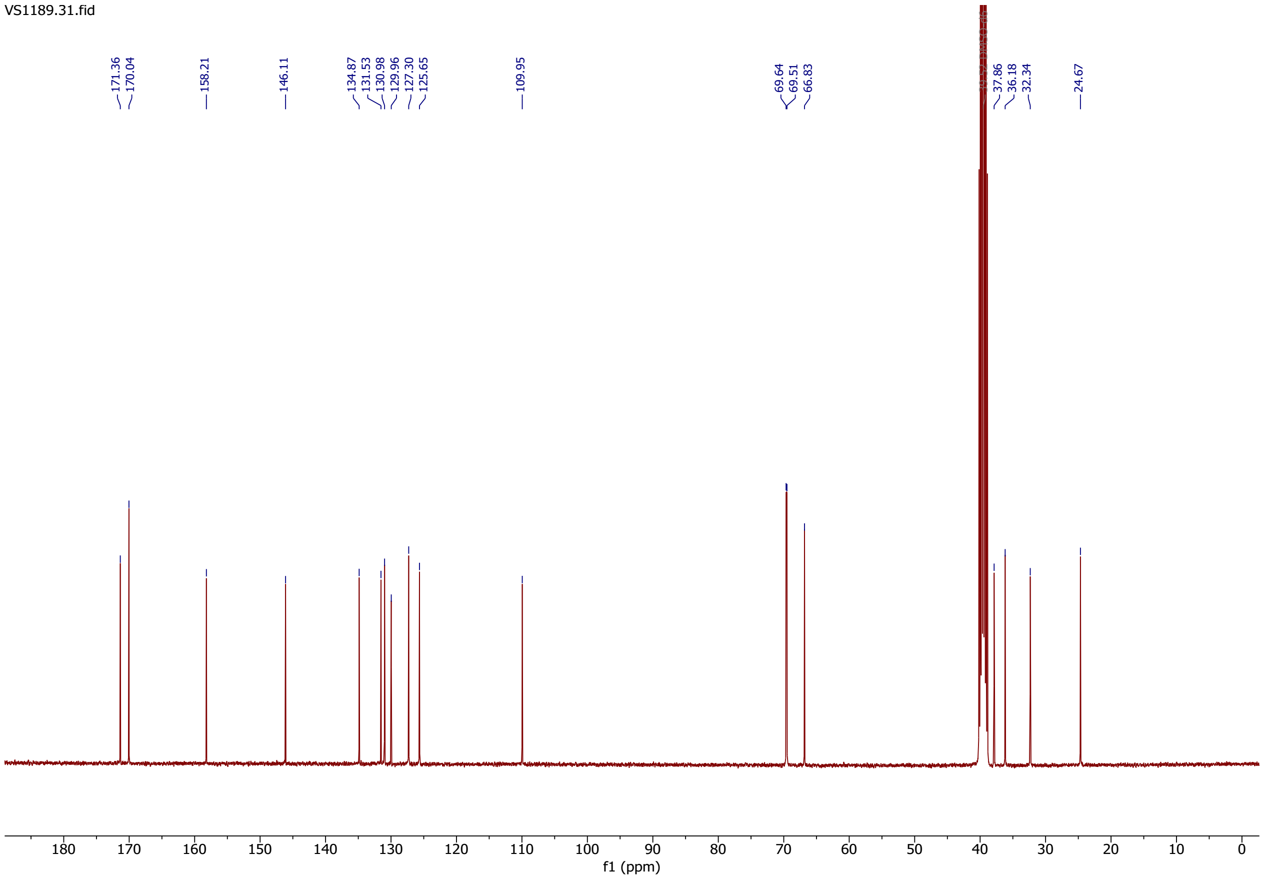
VS1189.31.fid
171.36 170.04 158.21 146.11 134.87 131.53 130.98 129.96 127.30 125.65 109.95 69.64 69.51 66.83 37.86 36.18 32.34 24.67
f1 (ppm)

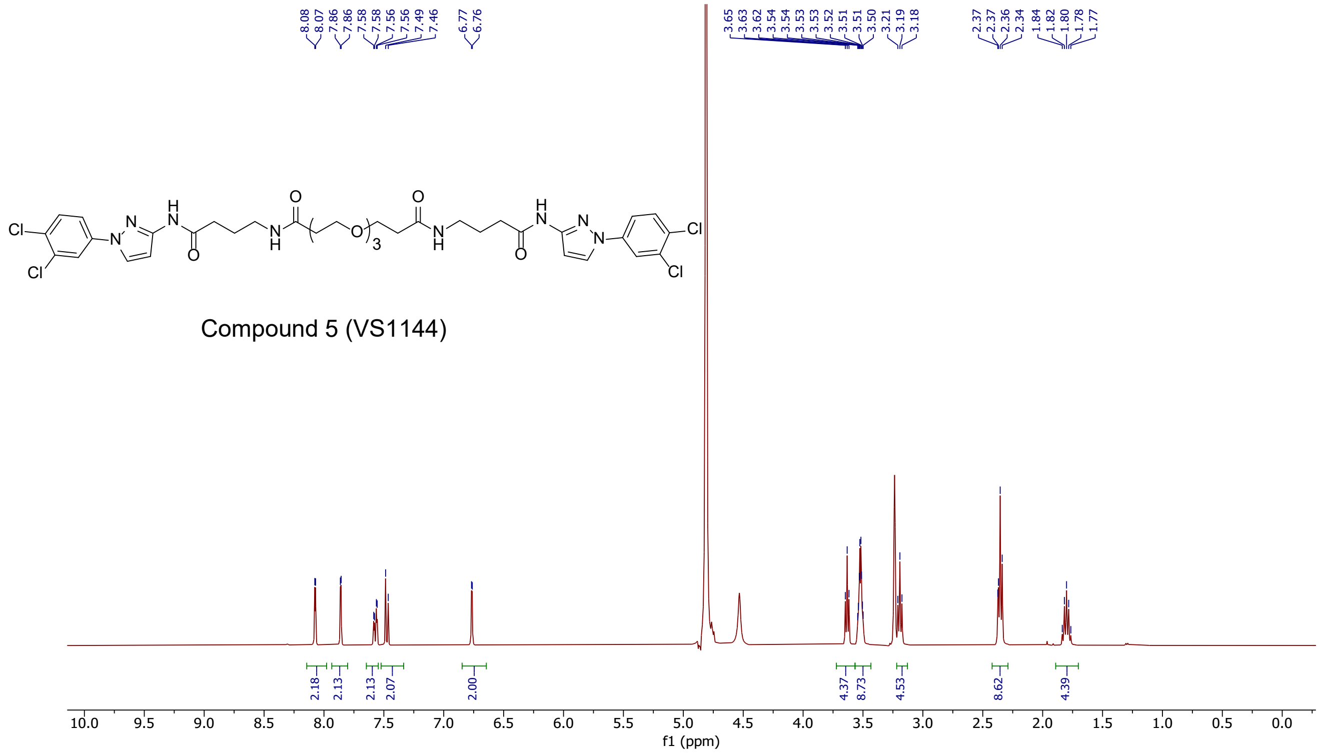
8.08 8.07 7.86 7.86 7.58 7.58 7.56 7.56 7.49 7.46 6.77 6.76
3.65 3.63 3.62 3.54 3.54 3.53 3.53 3.52 3.51 3.51 3.50 3.21 3.19 3.18
2.37 2.37 2.36 2.34 1.84 1.82 1.80 1.78 1.77
Compound 5 (VS1144)
2.18 2.13 2.13 2.07 2.00 4.37 8.73 4.53 8.62 4.39
10.0 9.5 9.0 8.5 8.0 7.5 7.0 6.5 6.0 5.5 5.0 4.5 4.0 3.5 3.0 2.5 2.0 1.5 1.0 0.5 0.0
f1 (ppm)

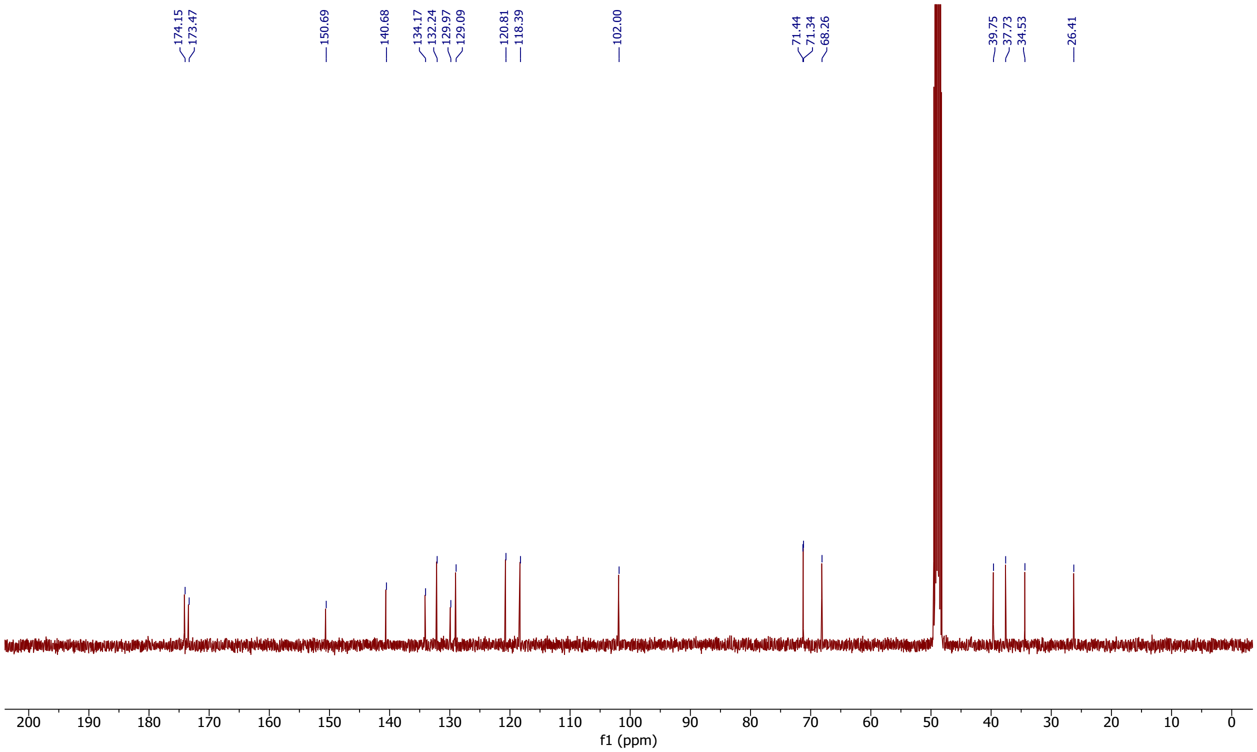
174.15 173.47 150.69 140.68 134.17 132.24 129.97 129.09 120.81 118.39 102.00
71.44 71.34 68.26 39.75 37.73 34.53 26.41
200 190 180 170 160 150 140 130 120 110 100 90 80 70 60 50 40 30 20 10 0
f1 (ppm)

Compound 6 (VS1114B)
Racemic mixture

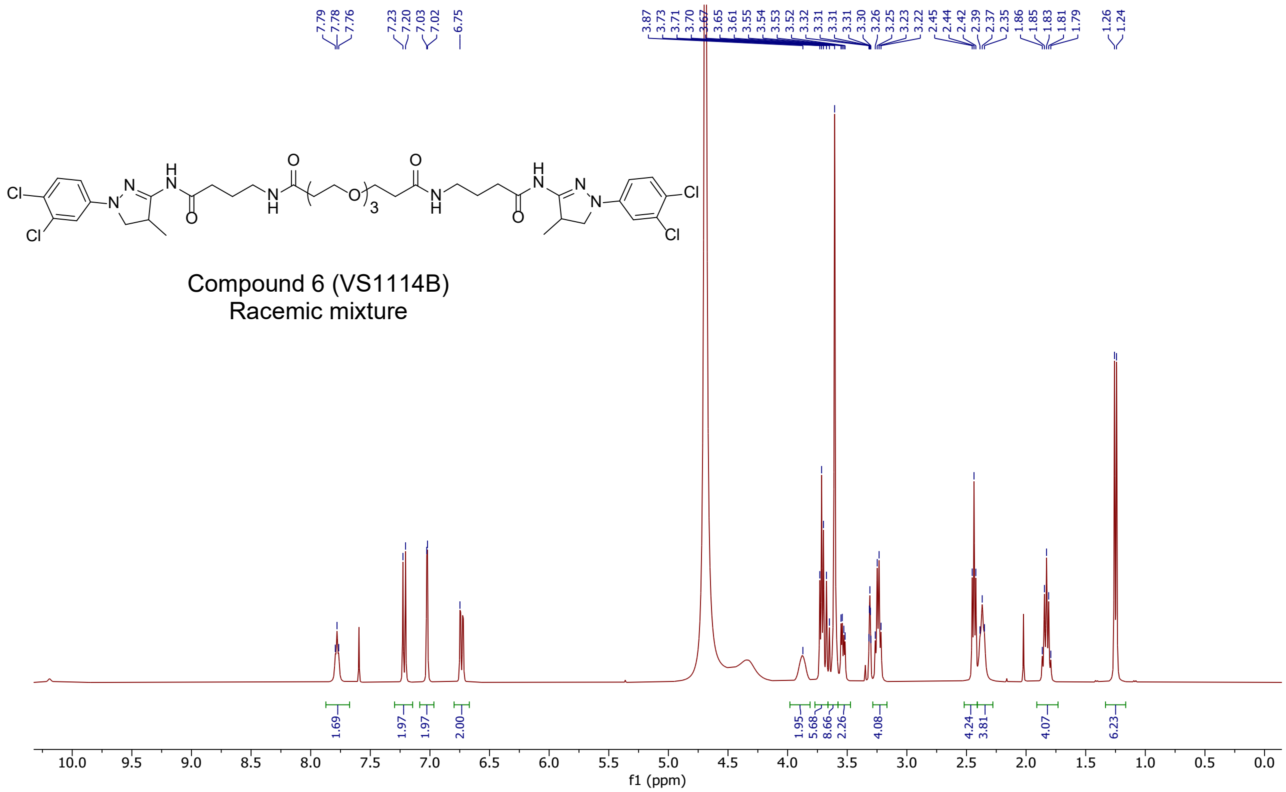


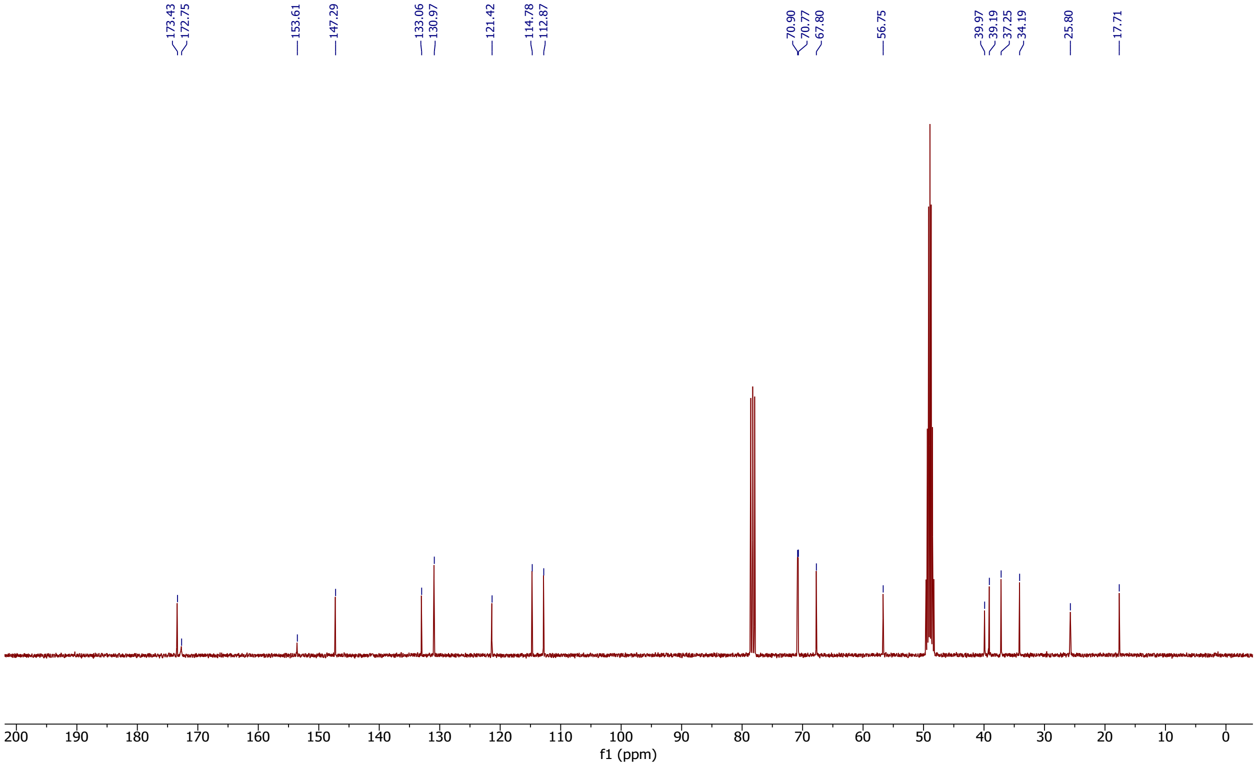

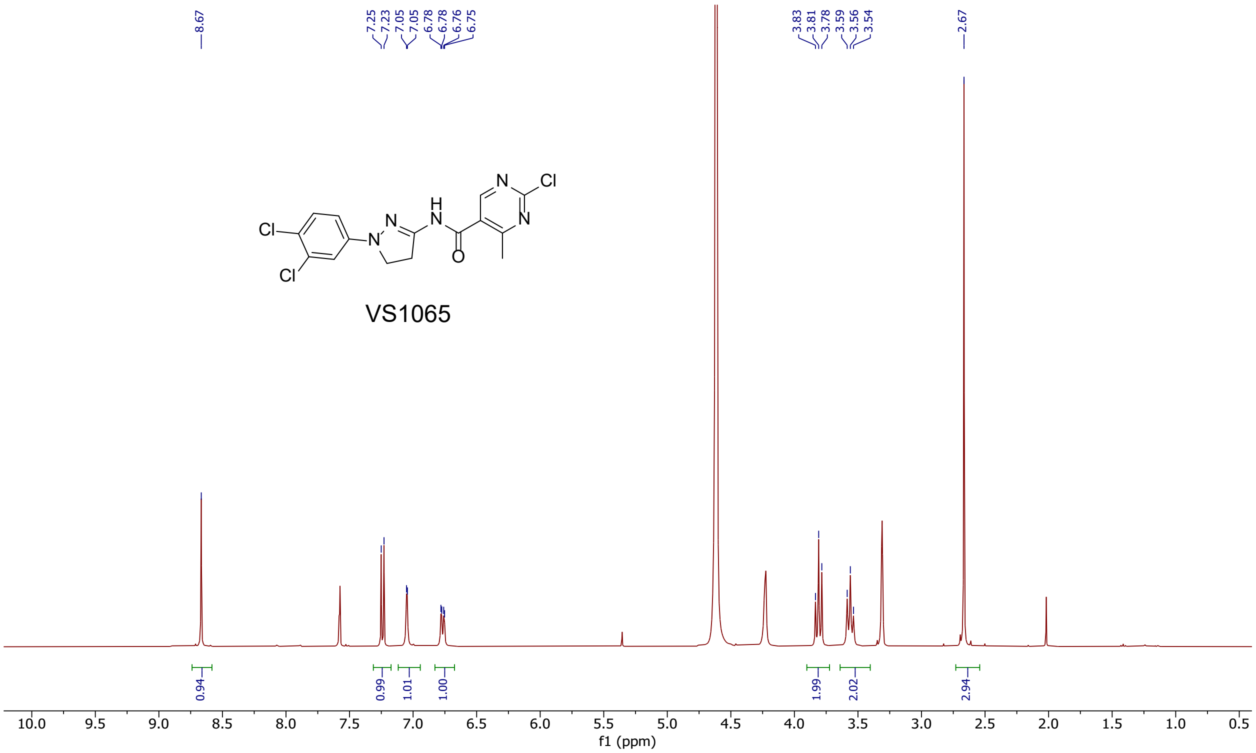
8.67
7.25 7.23 7.05 7.05 6.78 6.78 6.76 6.75
3.83 3.81 3.78 3.59 3.56 3.54
2.67
Cl
Cl
N
N
H
N
O
N
Cl
N
VS1065
0.94
0.99
1.01
1.00
1.99
2.02
2.94
10.0 9.5 9.0 8.5 8.0 7.5 7.0 6.5 6.0 5.5 5.0 4.5 4.0 3.5 3.0 2.5 2.0 1.5 1.0 0.5
f1 (ppm)

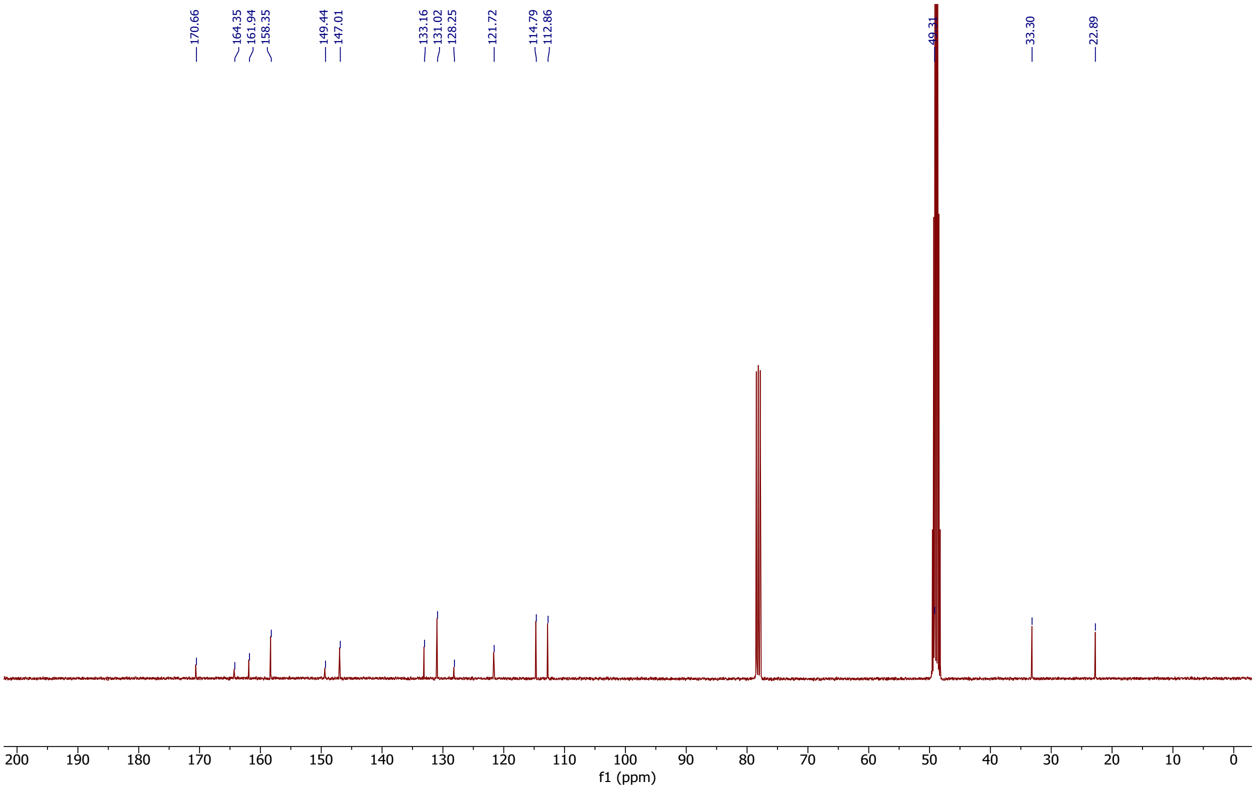
170.66
164.35 161.94 158.35
149.44 147.01
133.16 131.02 128.25
121.72
114.79 112.86
49.31
33.30
22.89
200 190 180 170 160 150 140 130 120 110 100 90 80 70 60 50 40 30 20 10 0
f1 (ppm)

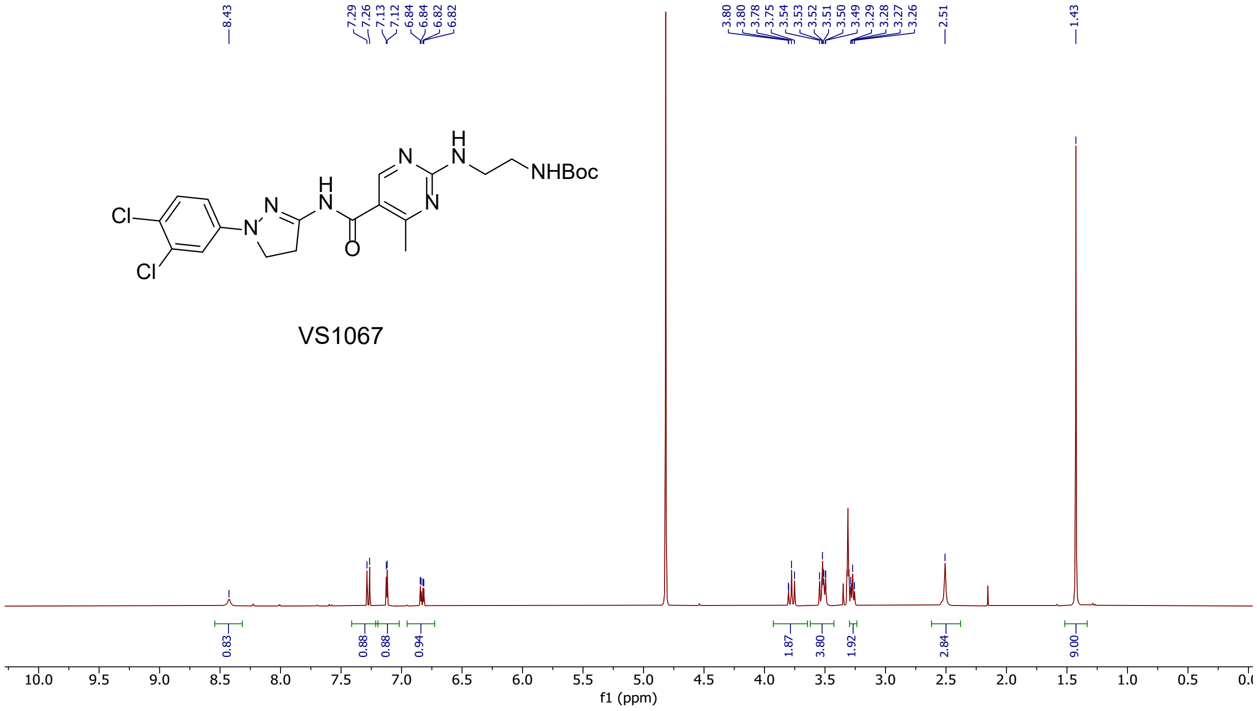
8.43
7.29
7.26
7.13
7.12
6.84
6.84
6.82
6.82
3.80
3.80
3.78
3.75
3.54
3.53
3.52
3.51
3.50
3.49
3.29
3.28
3.27
3.26
2.51
1.43
Cl
Cl
N
N
H
N
O
N
N
H
N
NHBoc
VS1067
0.83
0.88
0.88
0.94
1.87
3.80
1.92
2.84
9.00
f1 (ppm)

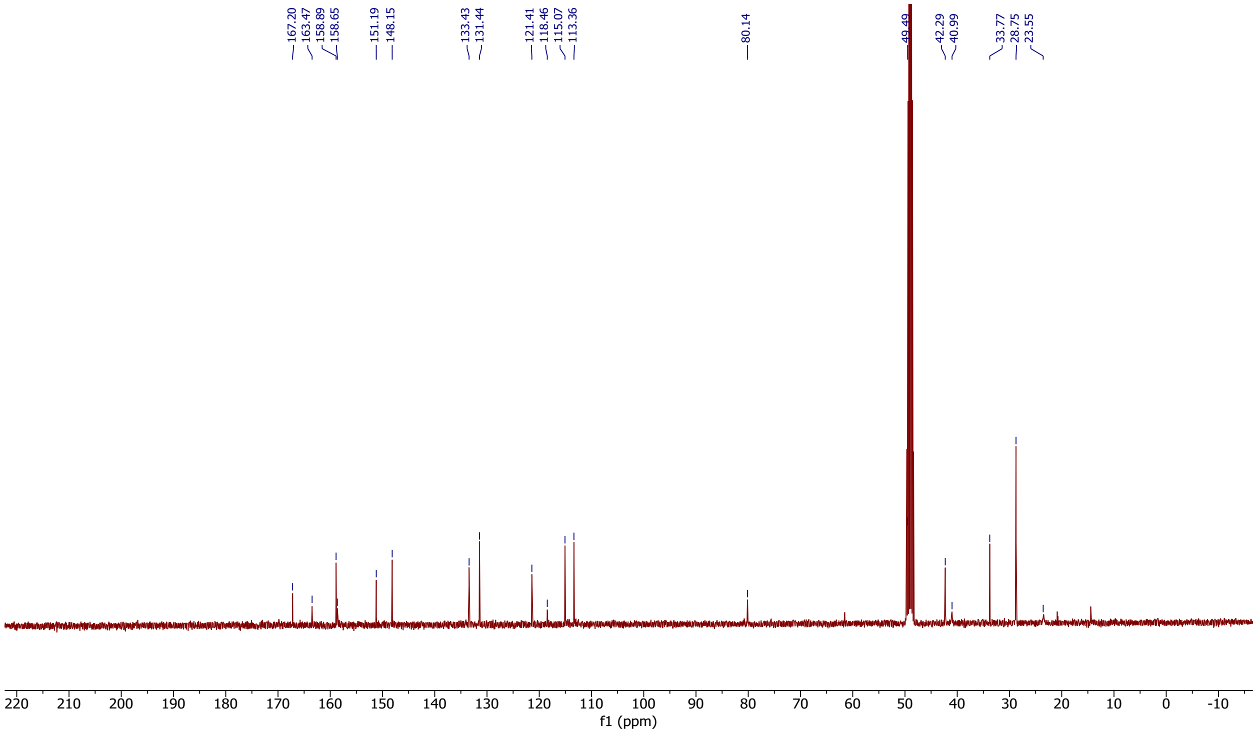
167.20
163.47
158.89
158.65
151.19
148.15
133.43
131.44
121.41
118.46
115.07
113.36
80.14
49.49
42.29
40.99
33.77
28.75
23.55
f1 (ppm)

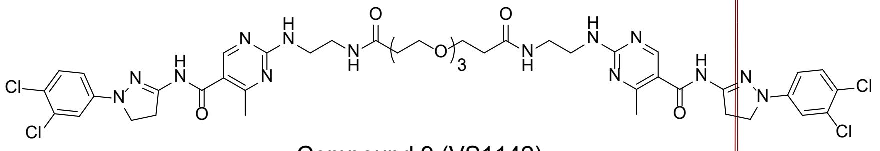


Compound 9 (VS1142)

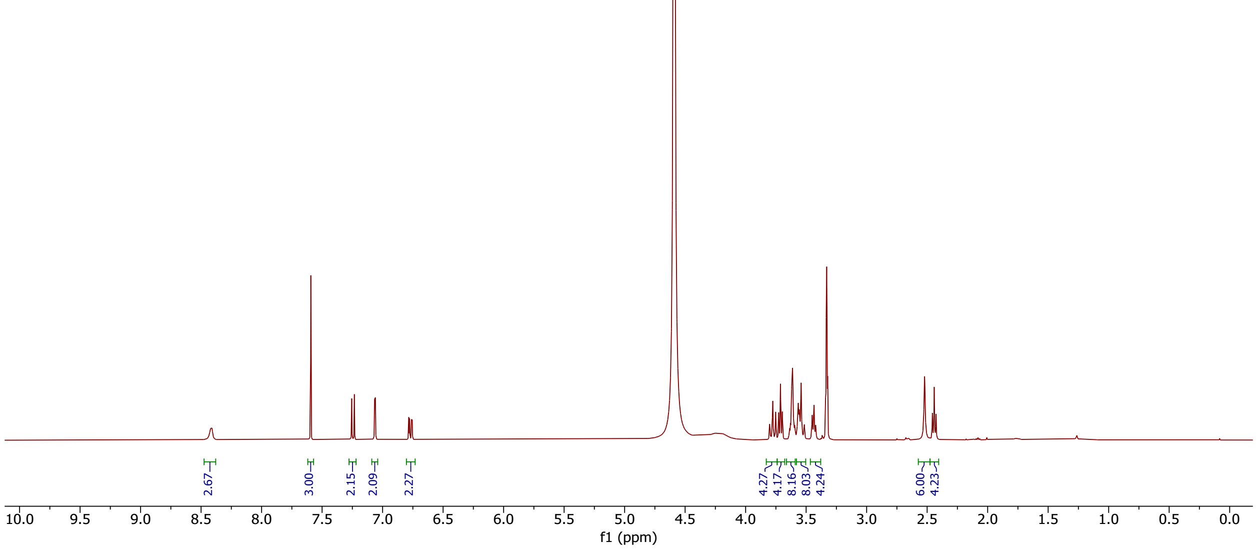


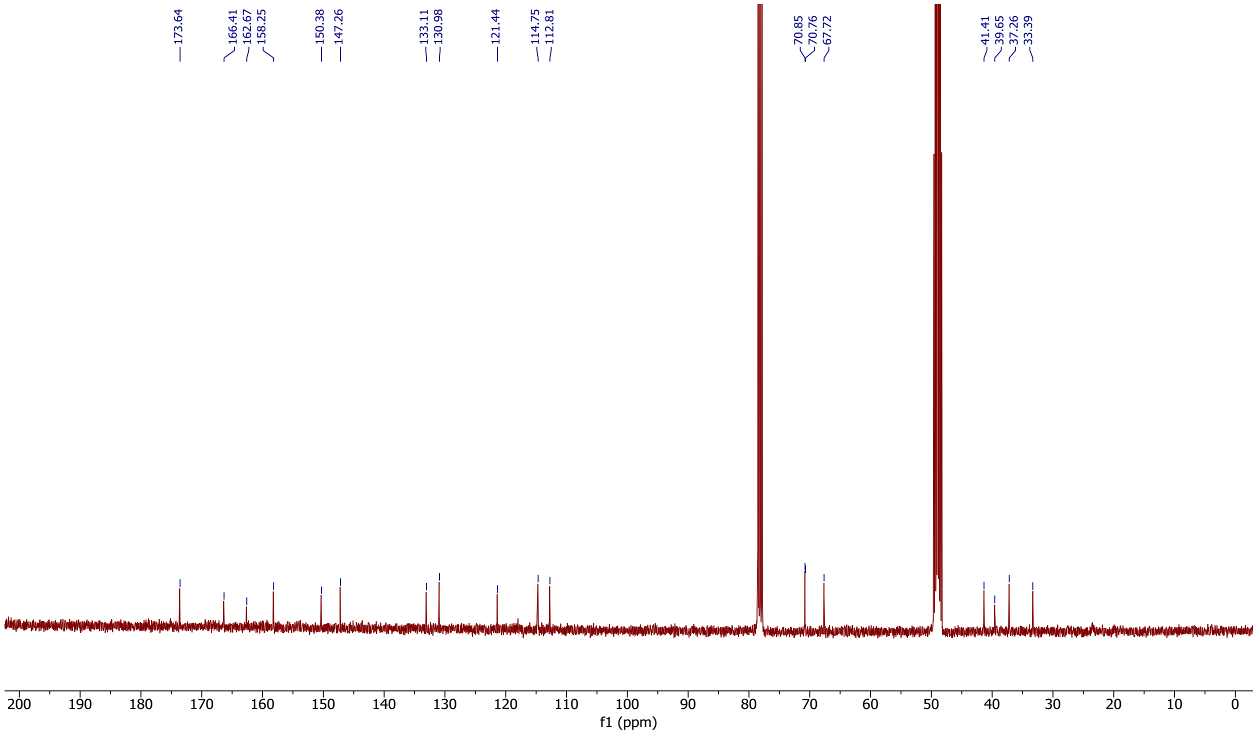

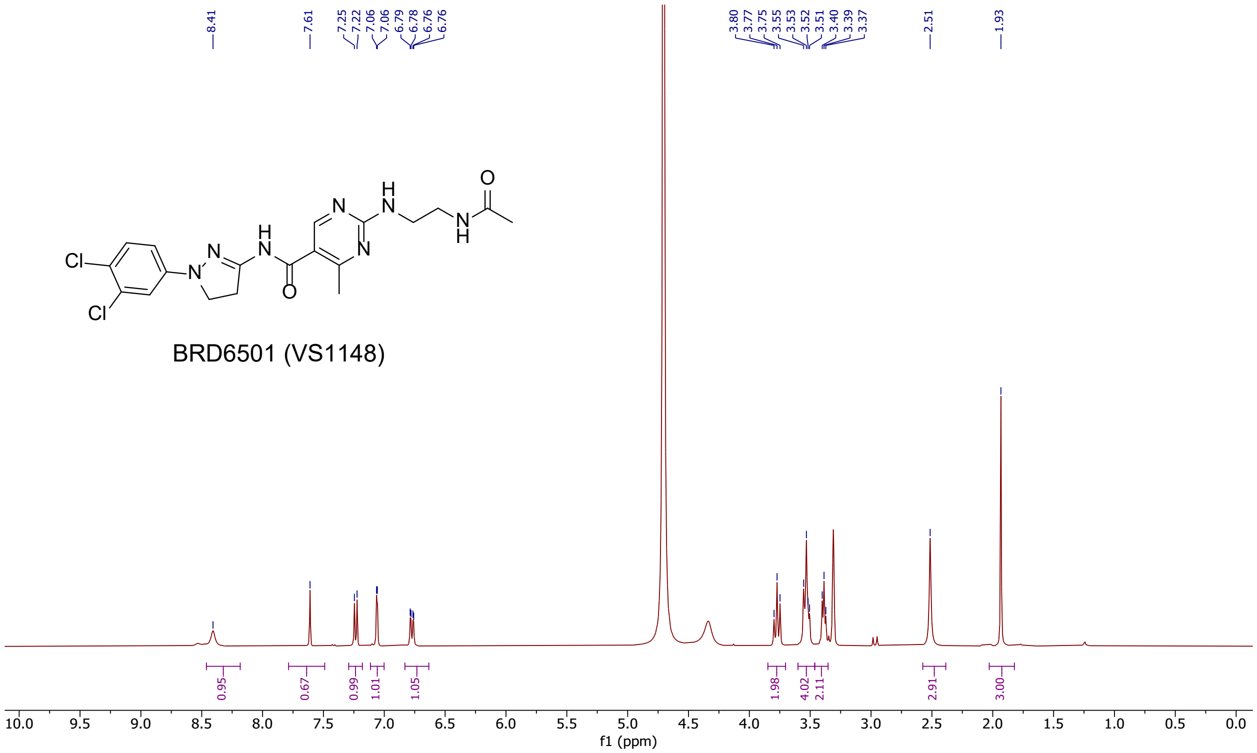

BRD6501 (VS1148)
8.41
7.61
7.25
7.22
7.06
7.06
6.79
6.78
6.76
6.76
3.80
3.77
3.75
3.55
3.53
3.52
3.51
3.40
3.39
3.37
2.51
1.93
0.95
0.67
0.99
1.01
1.05
1.98
4.02
2.11
2.91
3.00
f1 (ppm)


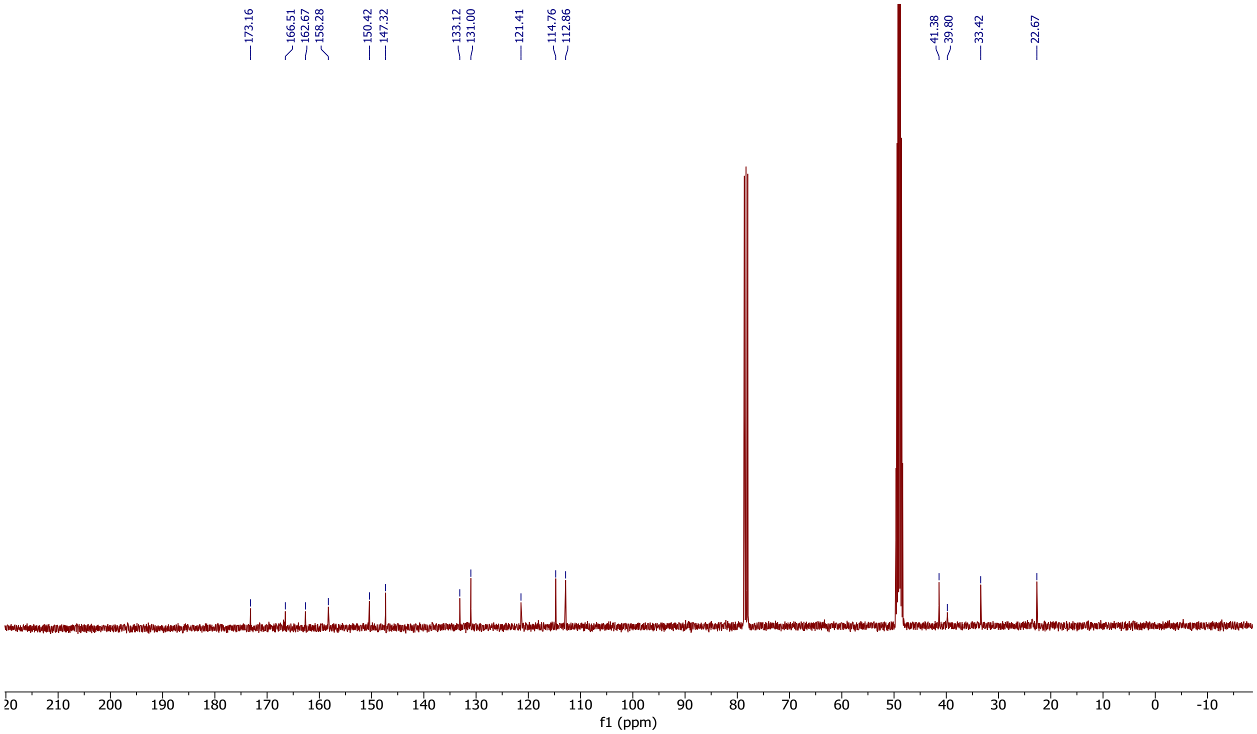

173.16
166.51
162.67
158.28
150.42
147.32
133.12
131.00
121.41
114.76
112.86
41.38
39.80
33.42
22.67
f1 (ppm)

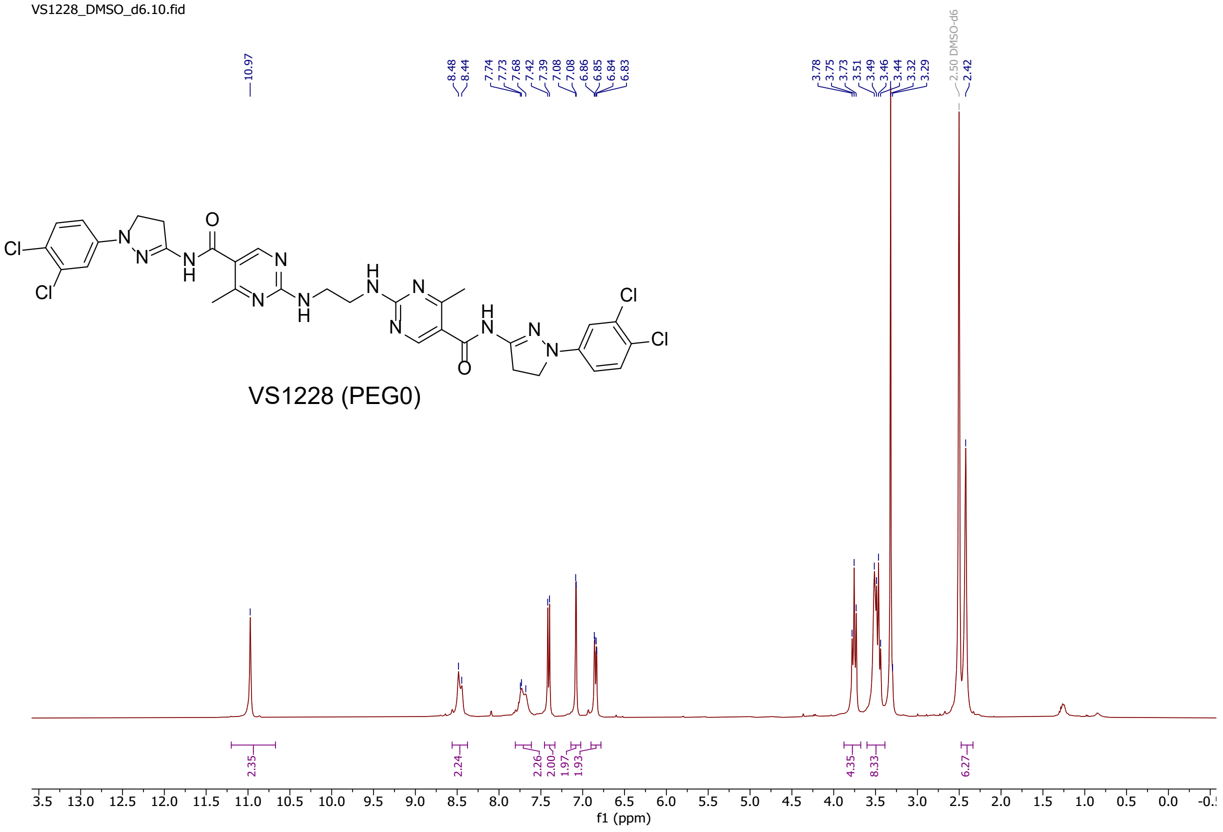
VS1228_DMSO_d6.10.fid
10.97
8.48
8.44
7.74
7.73
7.68
7.42
7.39
7.08
7.08
6.86
6.85
6.84
6.83
3.78
3.75
3.73
3.51
3.49
3.46
3.44
3.32
3.29
2.50 DMSO-d6
2.42
VS1228 (PEG0)
2.35
2.24
2.26
2.00
1.97
1.93
4.35
8.33
6.27
f1 (ppm)

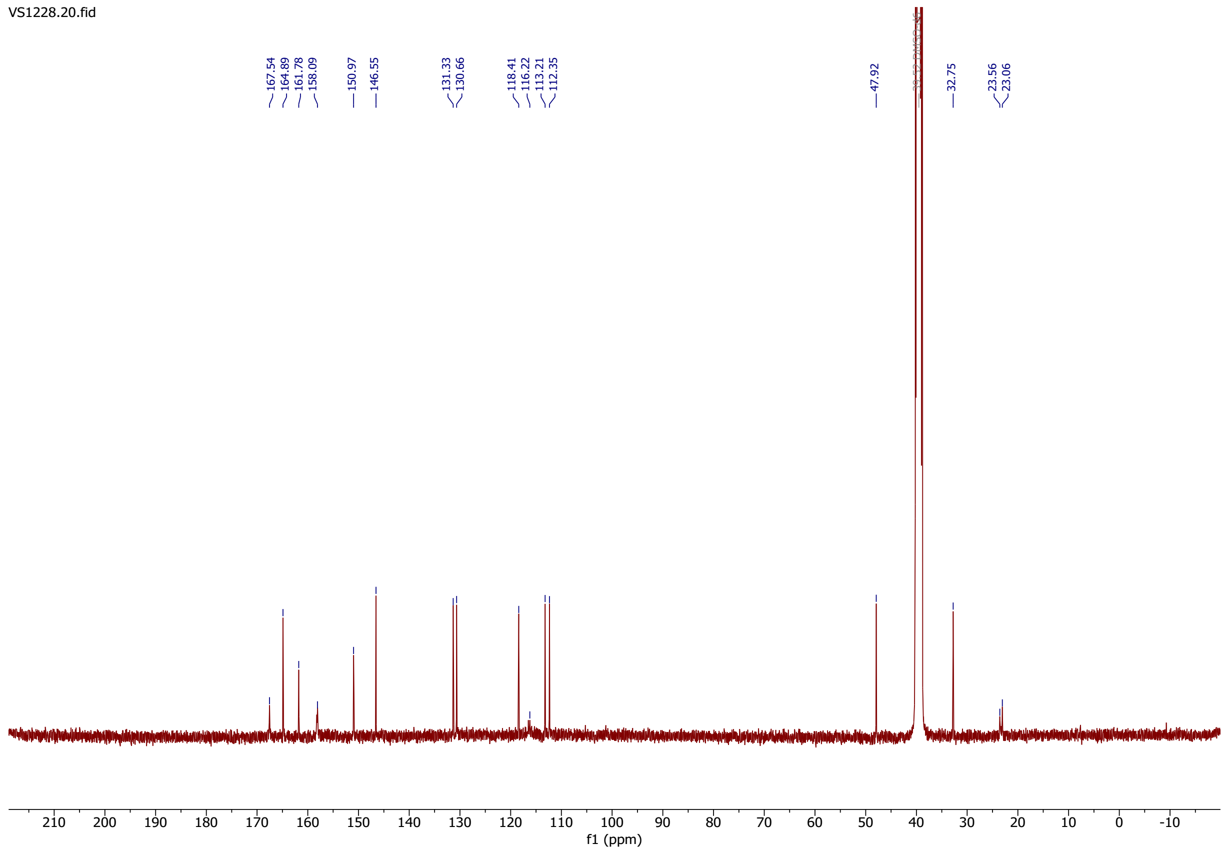
VS1228.20.fid
167.54
164.89
161.78
158.09
150.97
146.55
131.33
130.66
118.41
116.22
113.21
112.35
47.92
32.75
23.56
23.06
f1 (ppm)

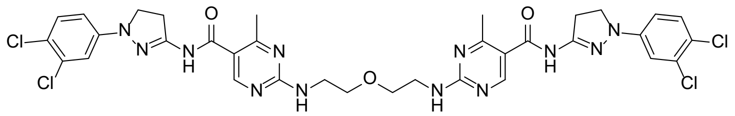

VS1163 (PEG1)

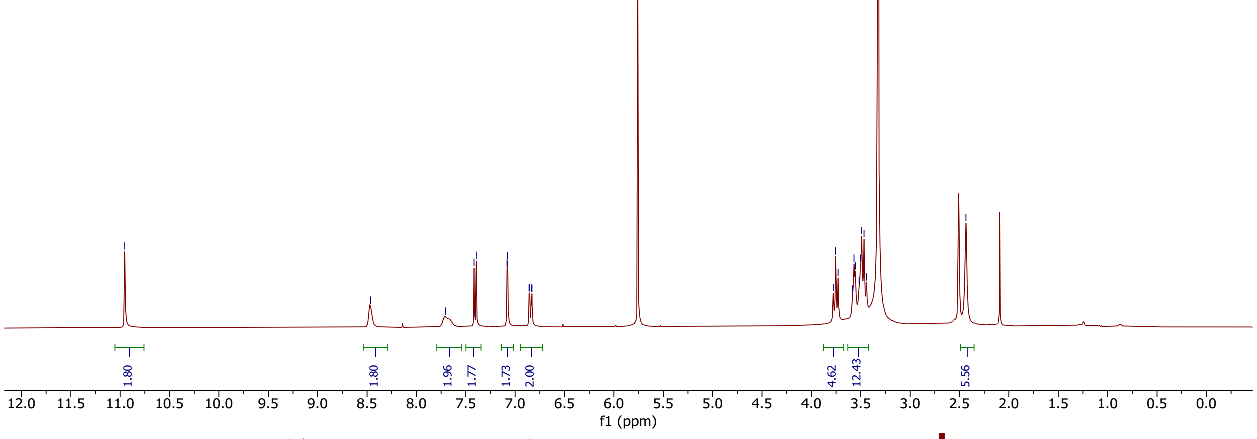


10.95
8.47
7.70
7.42
7.41
7.39
7.08
7.08
6.86
6.85
6.84
6.83
3.78
3.75
3.73
3.58
3.57
3.55
3.52
3.50
3.49
3.47
3.44
2.43
1.80
1.80
1.96
1.77
1.73
2.00
4.62
12.43
5.56
f1 (ppm)


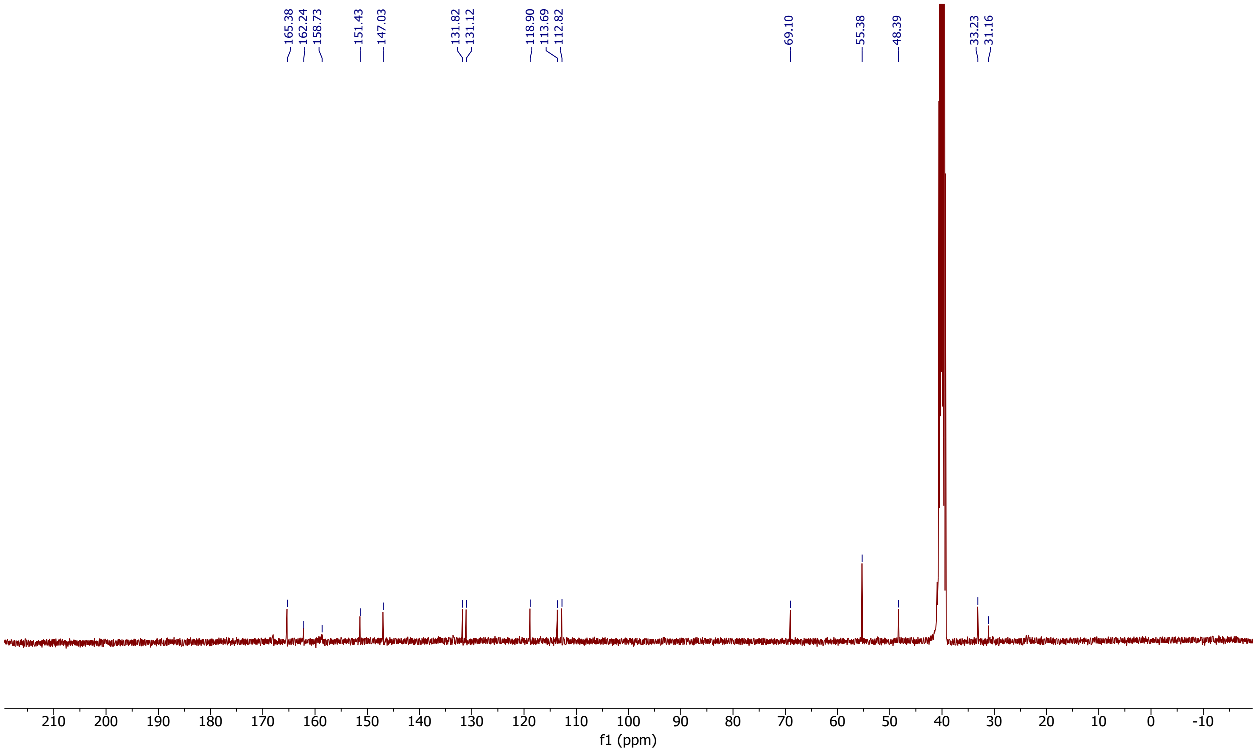


165.38
162.24
158.73
151.43
147.03
131.82
131.12
118.90
113.69
112.82
69.10
55.38
48.39
33.23
31.16
f1 (ppm)

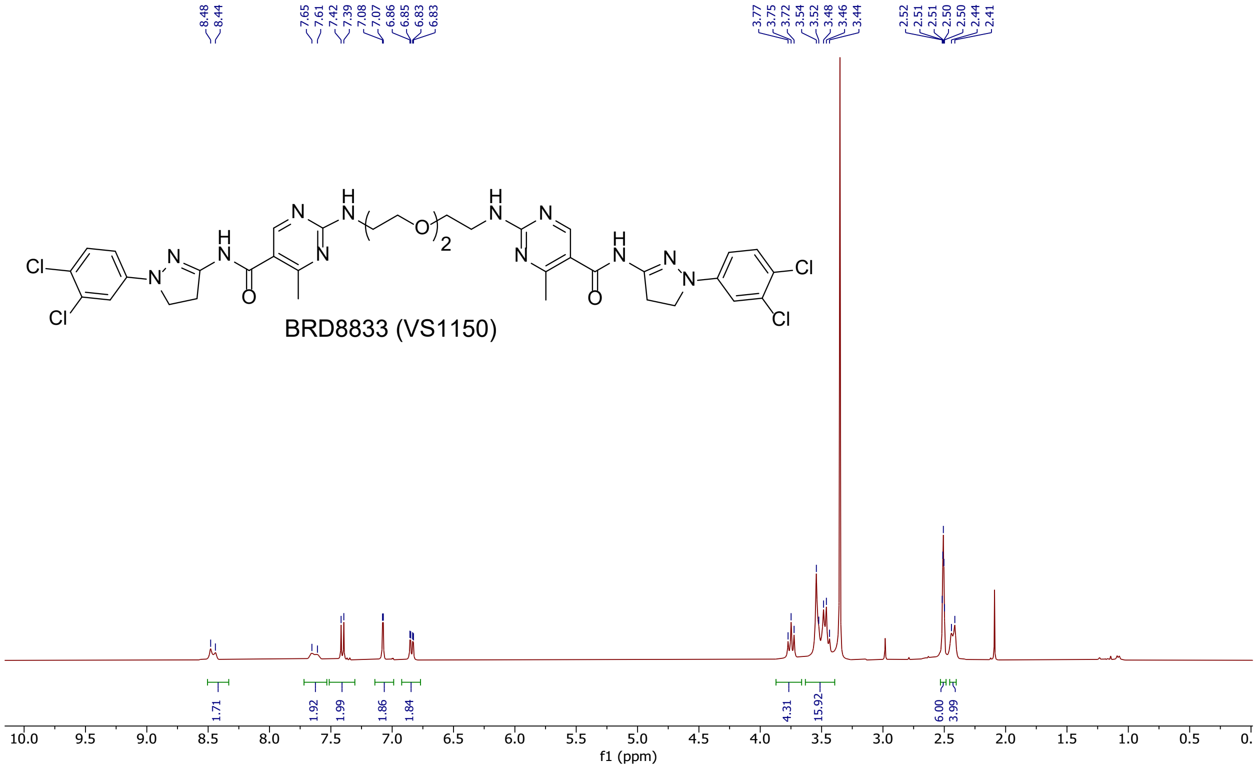
BRD8833 (VS1150)
Cl
Cl
N
H
O
8.48 8.44 7.65 7.61 7.42 7.39 7.08 7.07 6.86 6.85 6.83 6.83
3.77 3.75 3.72 3.54 3.52 3.48 3.46 3.44
2.52 2.51 2.51 2.50 2.50 2.44 2.41
1.71 1.92 1.99 1.86 1.84 4.31 15.92 6.00 3.99
f1 (ppm)

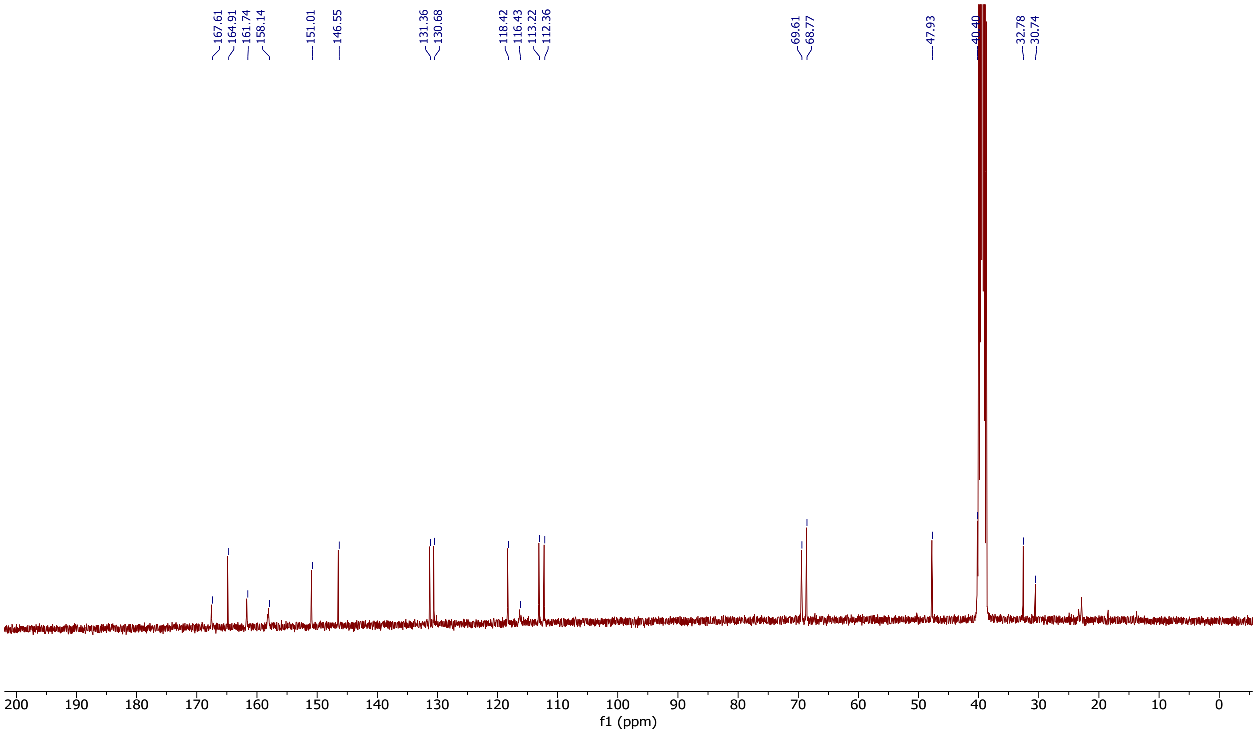
167.61 164.91 161.74 158.14 151.01 146.55 131.36 130.68 118.42 116.43 113.22 112.36 69.61 68.77 47.93 40.40 32.78 30.74
f1 (ppm)

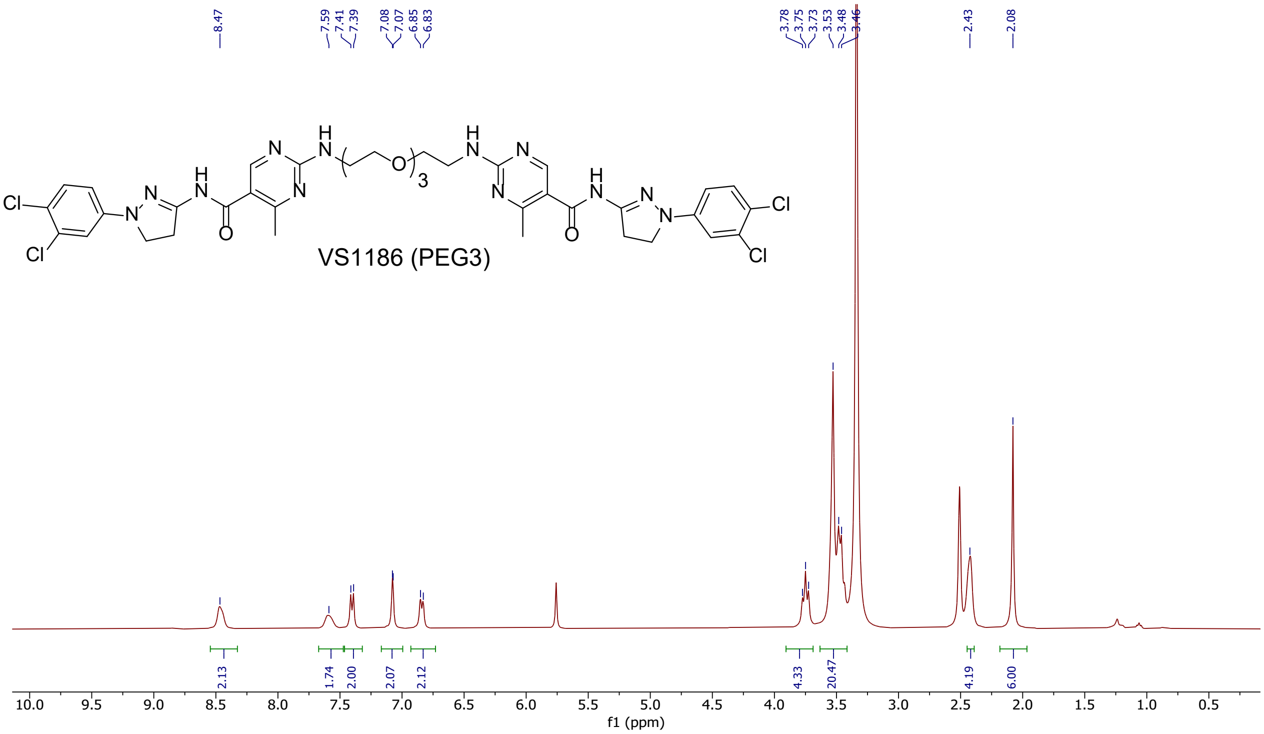

8.47
7.59
7.41
7.39
7.08
7.07
6.85
6.83
3.78
3.75
3.73
3.53
3.48
3.46
2.43
2.08
2.13
1.74
2.00
2.07
2.12
4.33
20.47
4.19
6.00
10.0 9.5 9.0 8.5 8.0 7.5 7.0 6.5 6.0 5.5 5.0 4.5 4.0 3.5 3.0 2.5 2.0 1.5 1.0 0.5
f1 (ppm)


VS1186 (PEG3)

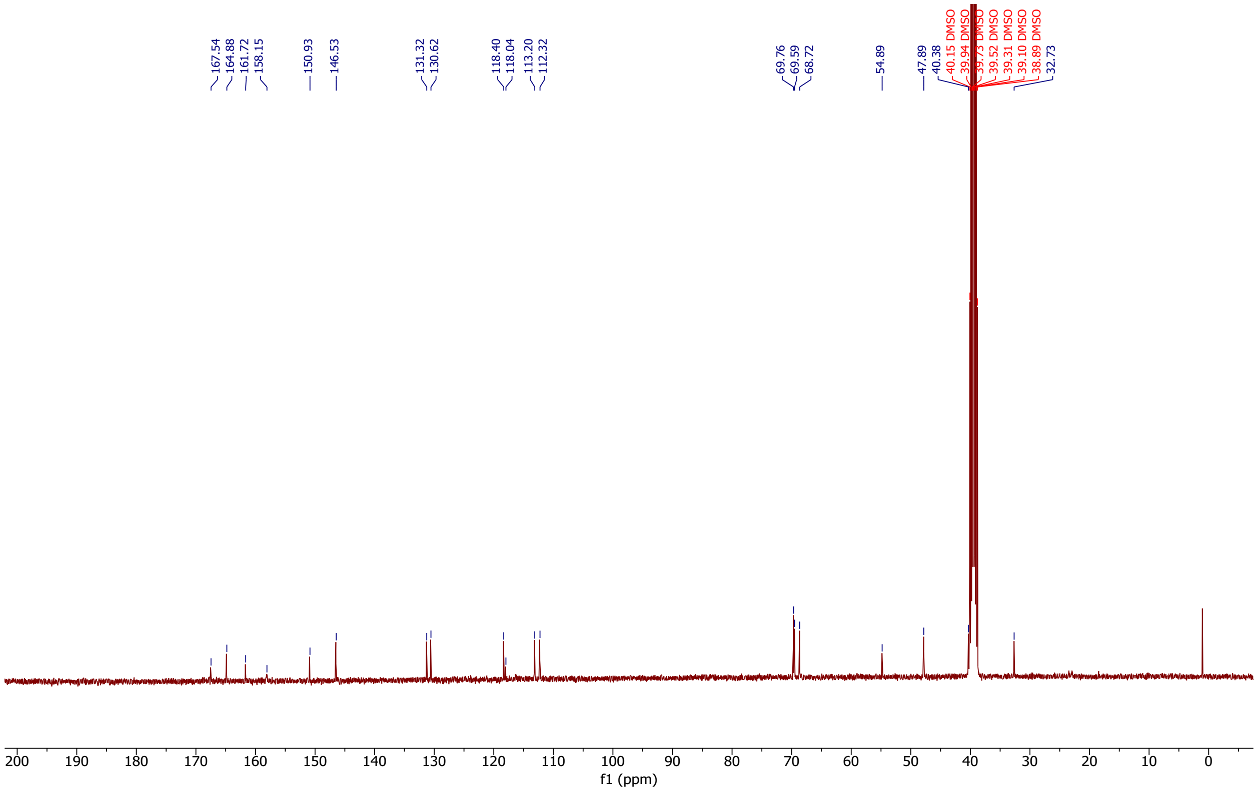

167.54
164.88
161.72
158.15
150.93
146.53
131.32
130.62
118.40
118.04
113.20
112.32
69.76
69.59
68.72
54.89
47.89
40.38
40.15 DMSO
39.94 DMSO
39.73 DMSO
39.52 DMSO
39.31 DMSO
39.10 DMSO
38.89 DMSO
32.73
200 190 180 170 160 150 140 130 120 110 100 90 80 70 60 50 40 30 20 10 0
f1 (ppm)

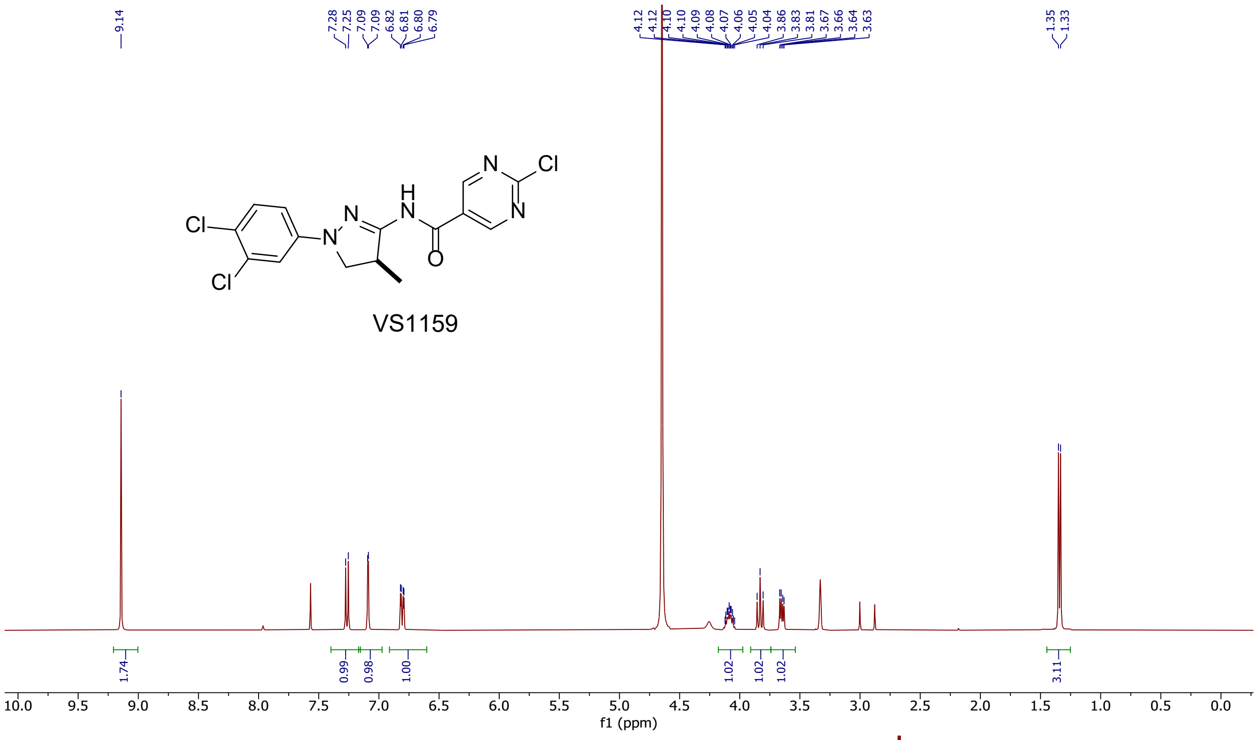
9.14
7.28 7.25 7.09 7.09 6.82 6.81 6.80 6.79
4.12 4.12 4.10 4.10 4.09 4.08 4.07 4.06 4.05 4.04 3.86 3.83 3.81 3.67 3.66 3.64 3.63
1.35 1.33
VS1159
1.74 0.99 0.98 1.00 1.02 1.02 1.02 3.11
f1 (ppm)

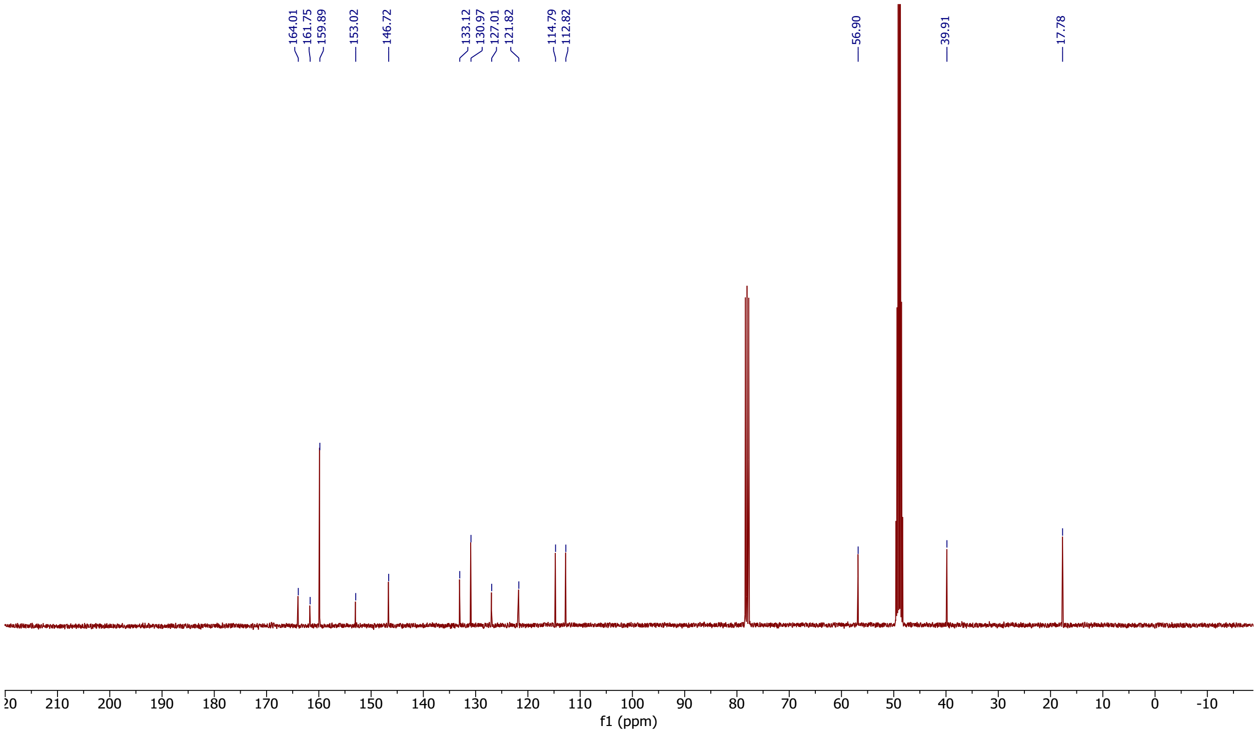
164.01 161.75 159.89 153.02 146.72 133.12 130.97 127.01 121.82 114.79 112.82 56.90 39.91 17.78
f1 (ppm)

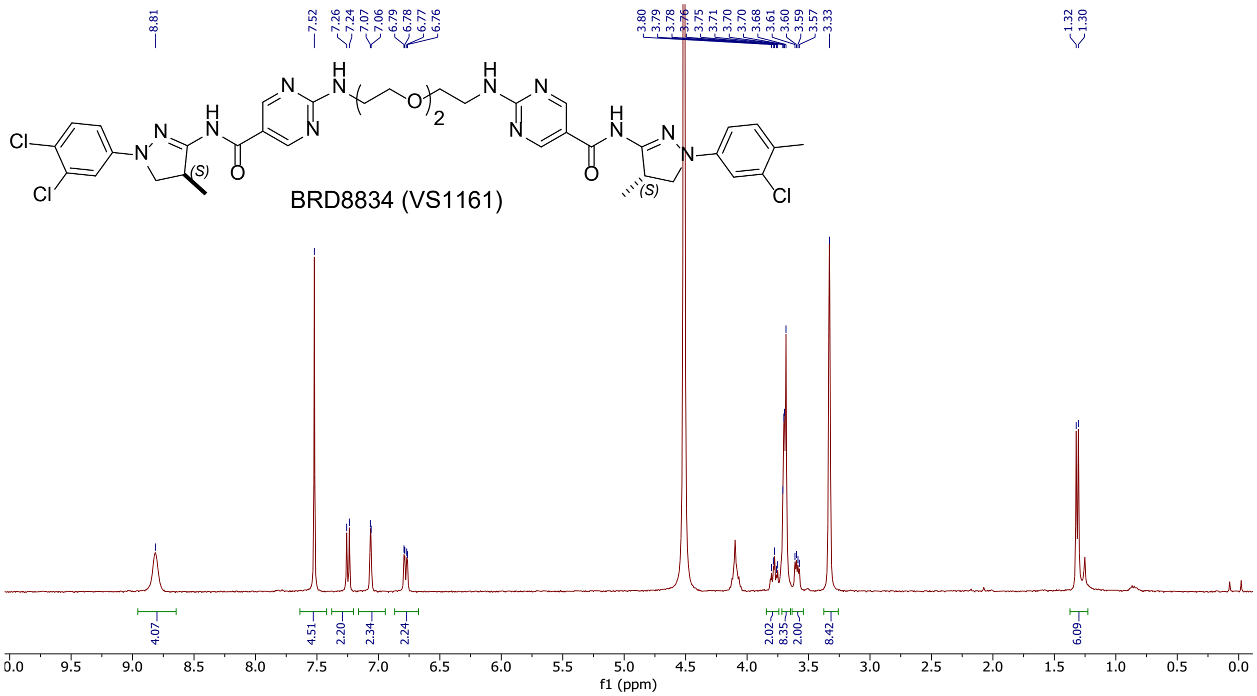
8.81
7.52 7.26 7.24 7.07 7.06 6.79 6.78 6.77 6.76
3.80 3.79 3.78 3.76 3.75 3.71 3.70 3.70 3.68 3.61 3.60 3.59 3.57 3.33
1.32 1.30
BRD8834 (VS1161)
4.07 4.51 2.20 2.34 2.24 2.02 8.35 2.00 8.42 6.09
f1 (ppm)

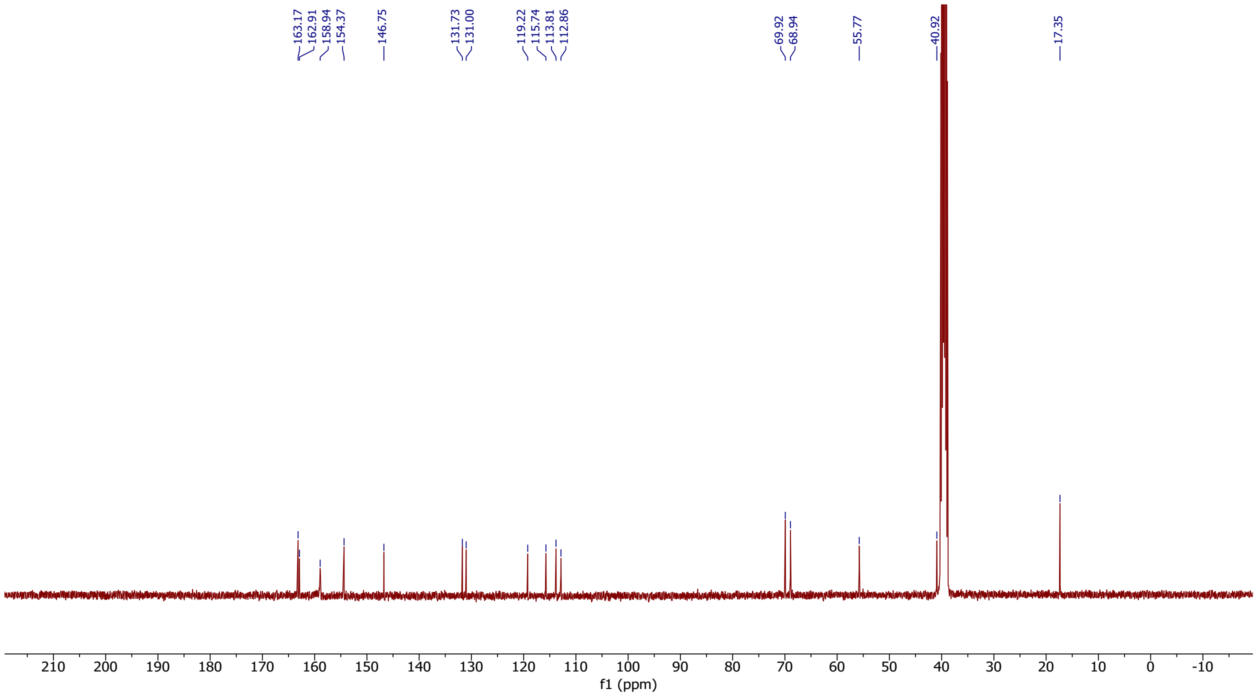
163.17 162.91 158.94 154.37 146.75 131.73 131.00 119.22 115.74 113.81 112.86 69.92 68.94 55.77 40.92 17.35
f1 (ppm)

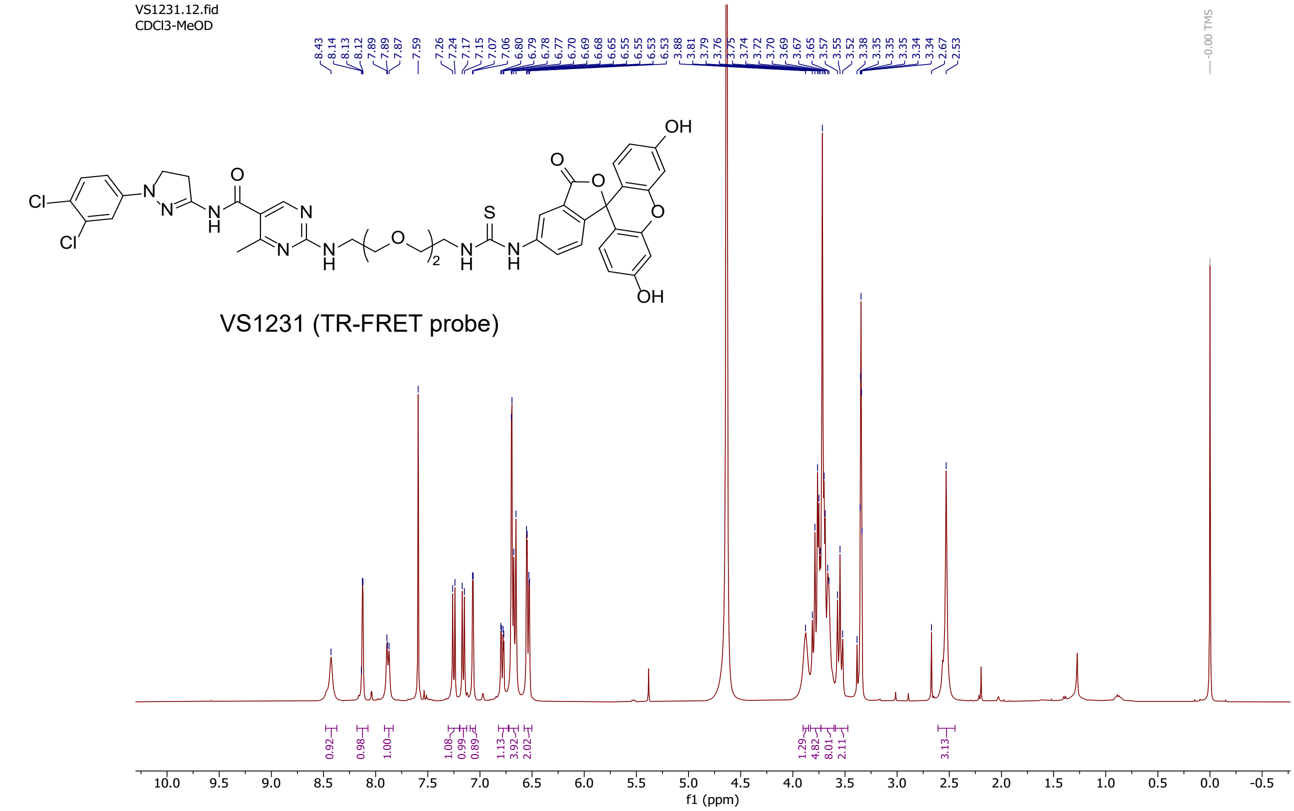
VS1231.12.fid
CDCl3-MeOD
8.43
8.14
8.13
8.12
7.89
7.89
7.87
7.59
7.26
7.24
7.17
7.15
7.07
7.06
6.80
6.79
6.78
6.77
6.70
6.69
6.68
6.65
6.55
6.55
6.53
6.53
3.88
3.81
3.79
3.76
3.75
3.74
3.72
3.70
3.69
3.67
3.65
3.57
3.55
3.52
3.38
3.35
3.35
3.35
3.34
3.34
2.67
2.53
0.00 TMS
VS1231 (TR-FRET probe)
0.92
0.98
1.00
1.08
0.99
0.89
1.13
3.92
2.02
1.29
4.82
8.01
2.11
3.13
f1 (ppm)

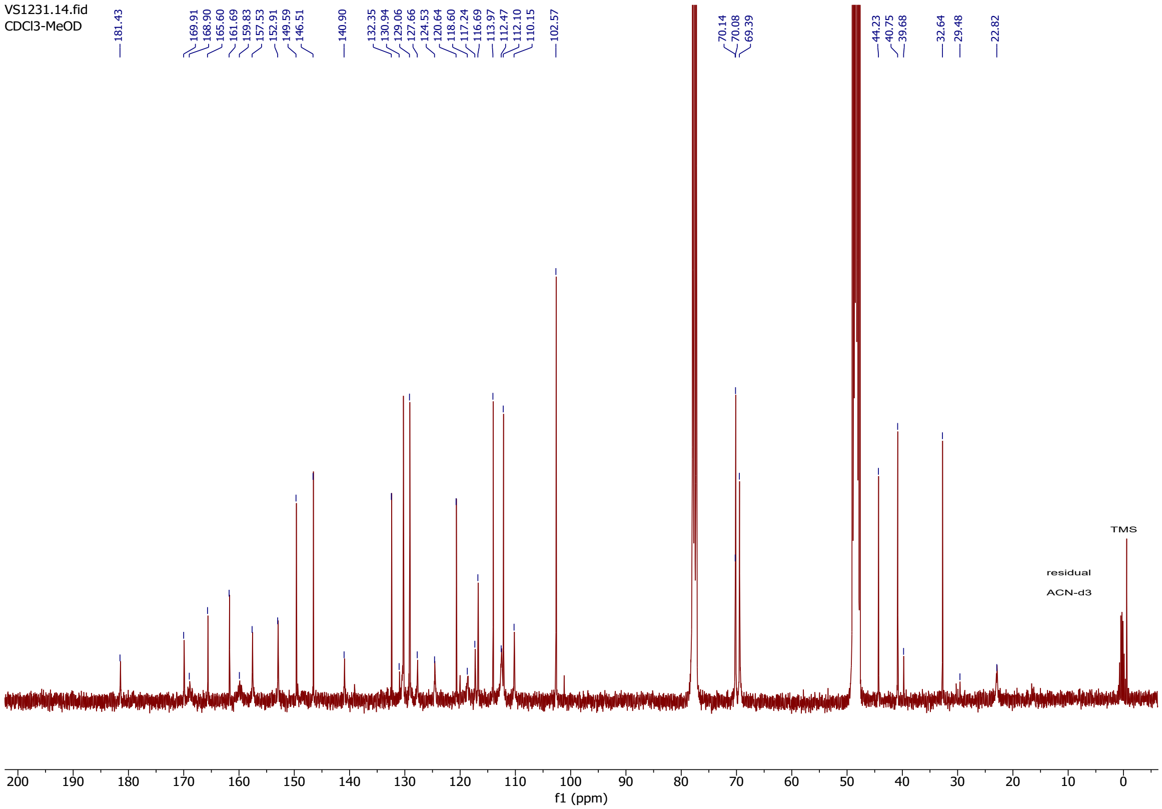
VS1231.14.fid
CDCl3-MeOD
181.43
169.91
168.90
165.60
161.69
159.83
157.53
152.91
149.59
146.51
140.90
132.35
130.94
129.06
127.66
124.53
120.64
118.60
117.24
116.69
113.97
112.47
112.10
110.15
102.57
70.14
70.08
69.39
44.23
40.75
39.68
32.64
29.48
22.82
TMS
residual
ACN-d3
f1 (ppm)

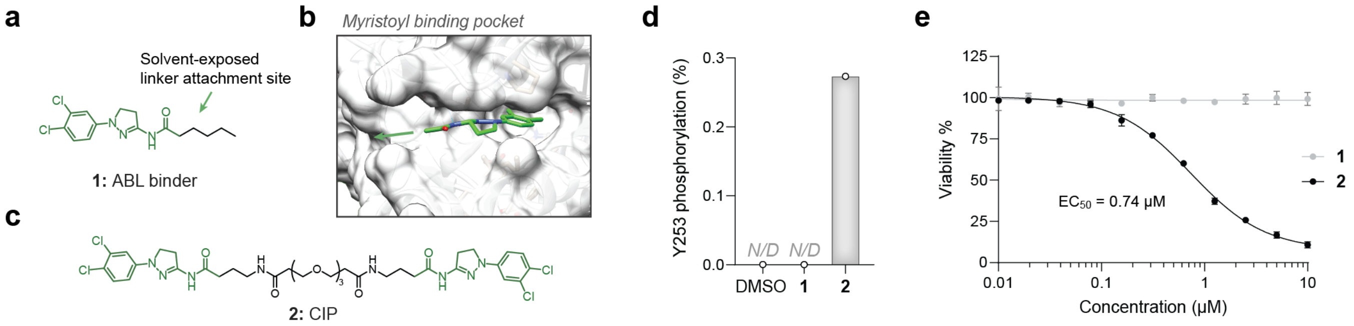


**Figure S1. CIP 2 is effective against BCR-ABL-dependent K562 cells. a** Chemical structure of ABL activator **1**. **b** Co-crystal structure of the activator binding to the allosteric myristoyl pocket of BCR-ABL (PDB: 6NPU). Arrows depict the solvent-exposed exit vector for linker attachment. **c** Chemical structure of BCR-ABL CIP **2**. **d** Analysis of biochemical ABL phosphorylation on Y253 by LC-MS/MS. The experiment was performed once (n = 1). **e** Dose response of **2** and its control **1** for cytotoxicity in K562. Mean and SD of three independent replicates are shown.

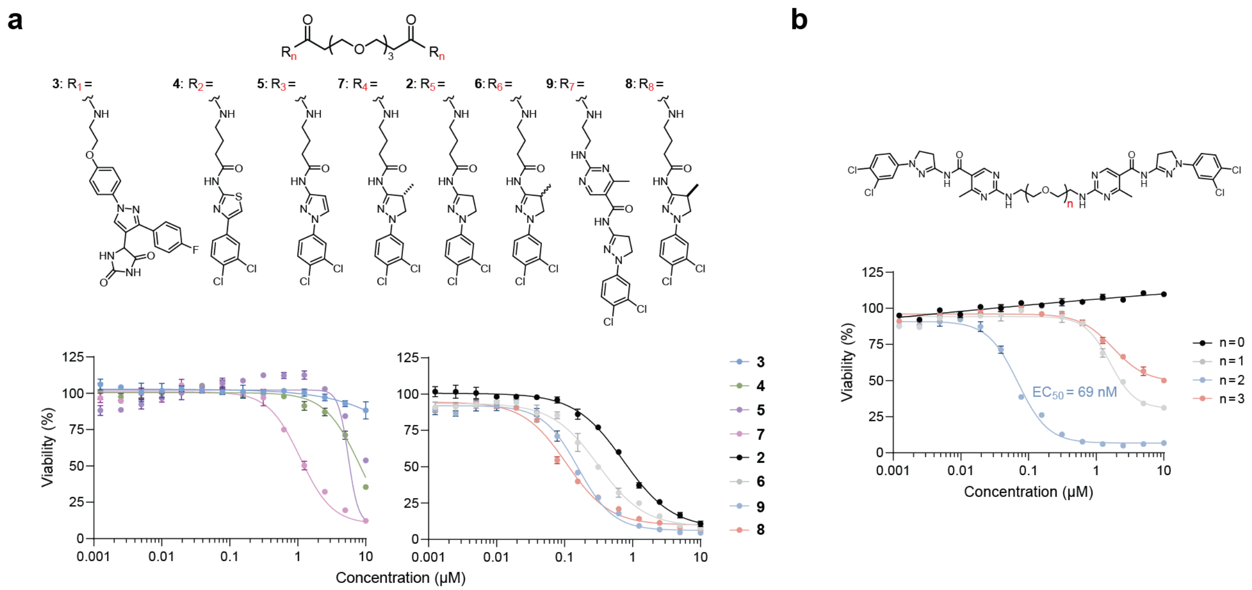


**Figure S2. Structure-activity relationship of BCR-ABL CIP. a–b** Optimization of the ABL binder (**a**) and linker (**b**) for cytotoxicity in K562 cells. Mean and SD of three independent replicates are shown.

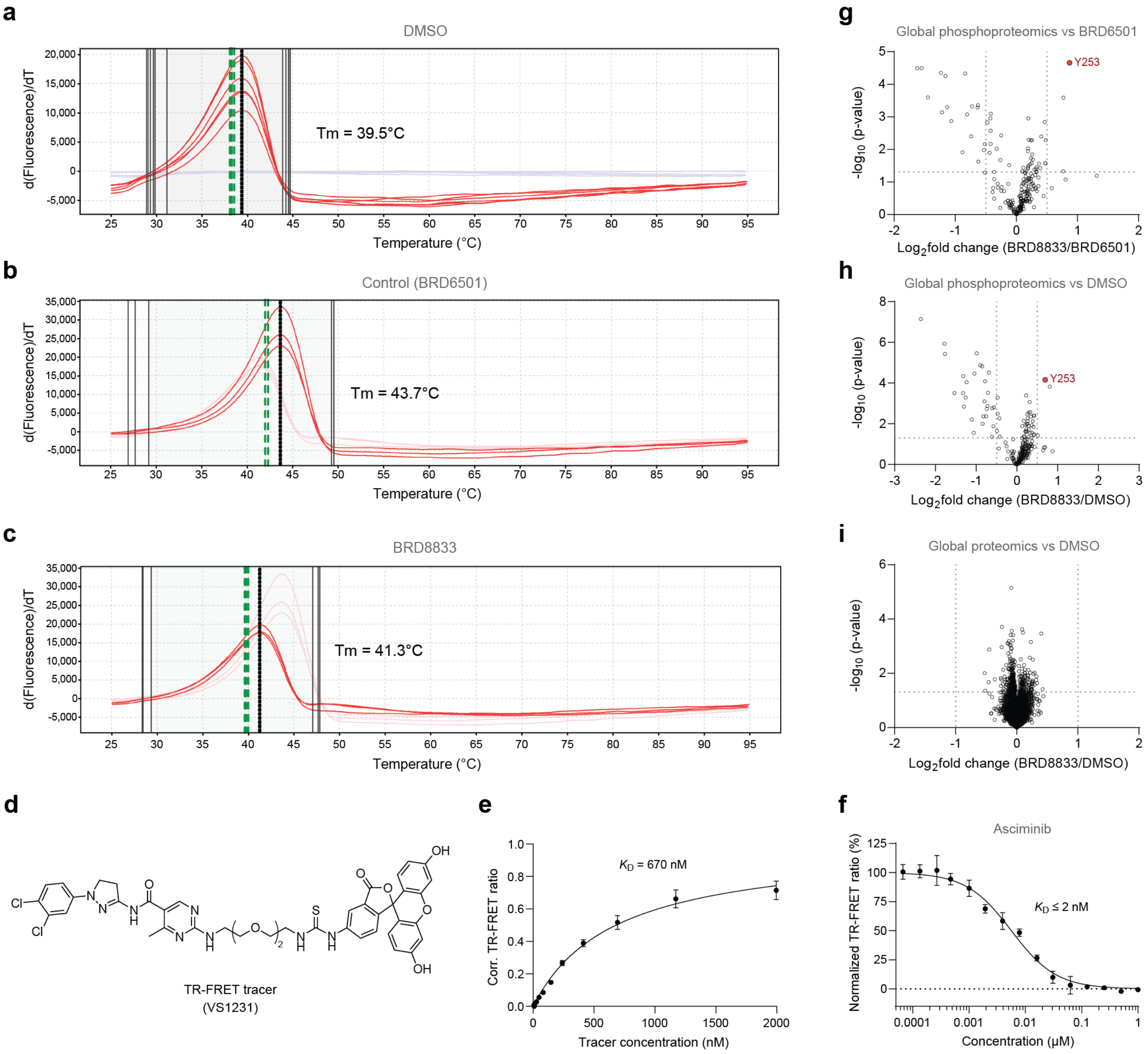


**Figure S3. BRD8833 binds to the myristoyl-binding pocket to induce BCR-ABL phosphorylation. a–c** Differential scanning fluorimetry assay to measure binding of control ABL binder (**b**) and BRD8833 (**c**) to the ABL kinase. Measurements were performed in three independent replicates (n = 3). **d–e** Structure (**d**) and binding curve (**e**) of TR-FRET tracer targeting the allosteric myristoyl pocket of the ABL kinase. **f** TR-FRET assay to measure dose-dependent displacement of the tracer by asciminib. Mean and SD of three independent replicates are shown. **g–i** Global phosphoproteomic and proteomic profiling in K562 cells. Volcano plot showing the $\log_2$-fold change of phosphopeptides normalized to the control ABL binder BRD6501 (**g**) or DMSO (**h**), using cutoffs of $\log_2$-fold change ≥0.5 and p-value ≤0.05 (Welch's t-test). For global proteomic analysis (**i**), a cutoff of $\log_2$-fold change ≥1 and p-value ≤0.05 was used. Proteomic and phosphoproteomic analysis was performed in four independent biological replicates (n = 4).

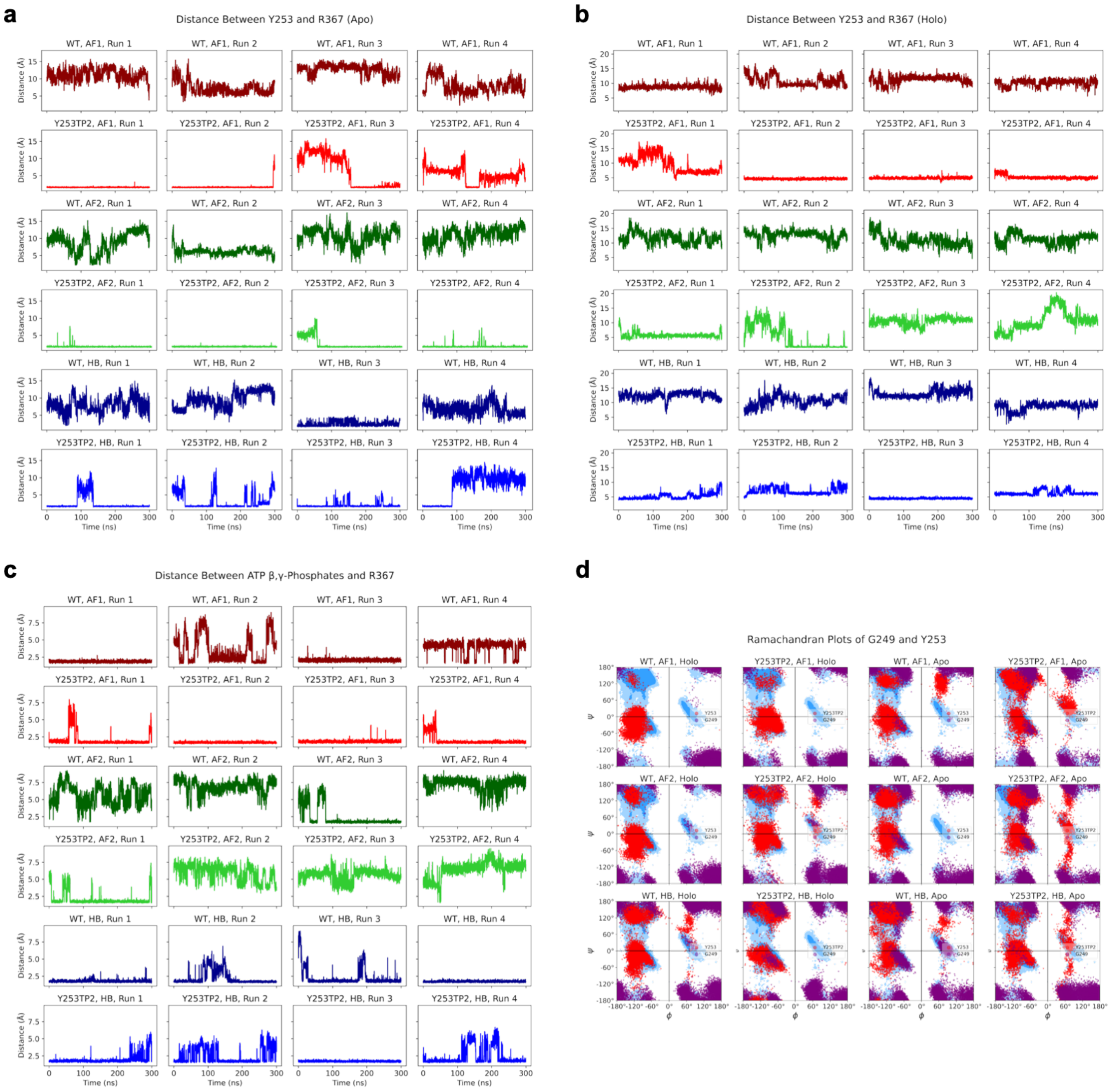


**Figure S4. Analysis of key distances and dihedral angles during molecular dynamics simulations of BCR-ABL. a–c** Time evolution of key distances: (**a**) minimum distance between Y253 and R367 in the apo state, (**b**) minimum distance between Y253 and R367 in the holo state, and (**c**) minimum distance between ATP β,γ-phosphates and R367. Results are shown for three different starting structures: the top-ranked AlphaFold3 model (AF1), the second-ranked AlphaFold3 model (AF2), and the "handbuilt" model (HB, see Supporting Methods). **d** Ramachandran plot of G249 and Y253 for every state and starting structure showing conformational change of the P-loop upon Y253 phosphorylation. These two residues were found to exhibit the largest changes in the backbone $\phi,\psi$ angles relative to the starting structure among the P-loop residues. Contours in the background indicate the "allowed" (blue) and the "marginally allowed" (light blue) regions.

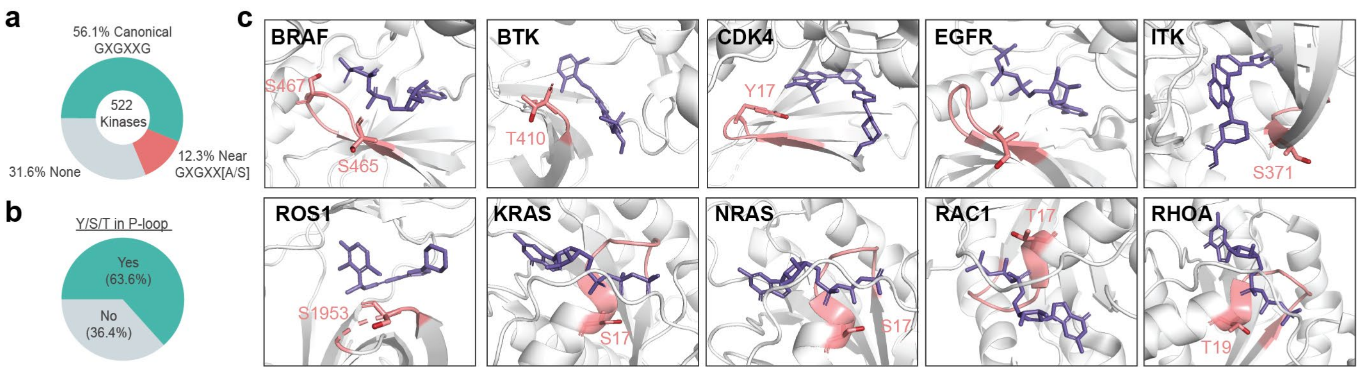


**Figure S5. S/T/Y residues in the P-loop of kinases and GTPases. a** Distribution of the 522 kinases in the human kinome containing either a canonical, a near, or no P loop motif. **b** Distribution of kinase P loop motifs containing at least one tyrosine, serine, or threonine residue. **c** Structures of relevant kinases (BRAF [PDB:6V2W], BTK [PDB:6NFI], CDK4 [PDB:7SJ3], EGFR [PDB:2ITX], ITK [PDB:4HCT], ROS1 [PDB:3ZBF]) or GTPases (KRAS [PDB:6GJ8], NRAS [PDB:5UHV], RAC1 [PDB:3TH5], RHOA [PDB:1KMQ]) with P-loop residues that can be phosphorylated. P-loops and S/T/Y residues that can be phosphorylated are colored in pink, and ATP-competitive inhibitors or GTP analogs are shown in purple.

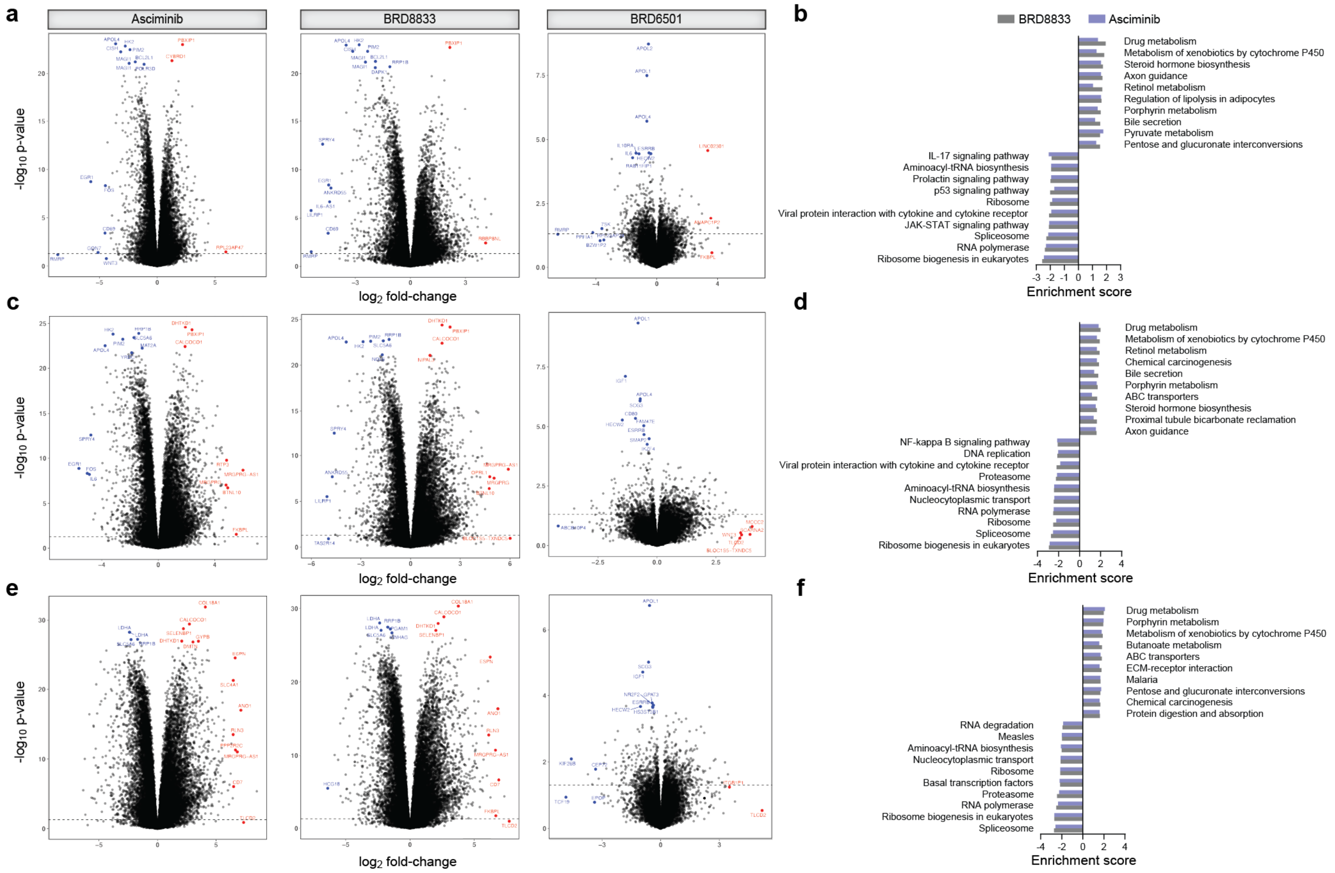


**Figure S6. Gene expression and gene set enrichment analysis.** Volcano plots showing gene expression, and gene set enrichment analysis vs. DMSO after 6 h (**a–b**), 12 h (**c–d**), and 24 h (**e–f**) treatment with asciminib, BRD8833, or control ABL binder BRD6501. A p-value cutoff of < 0.01 is used in the volcano plot. Genes with the lowest p-values or the highest $\log_2$-fold change are labeled. Each datapoint represents the average of four independent biological replicates (n = 4).

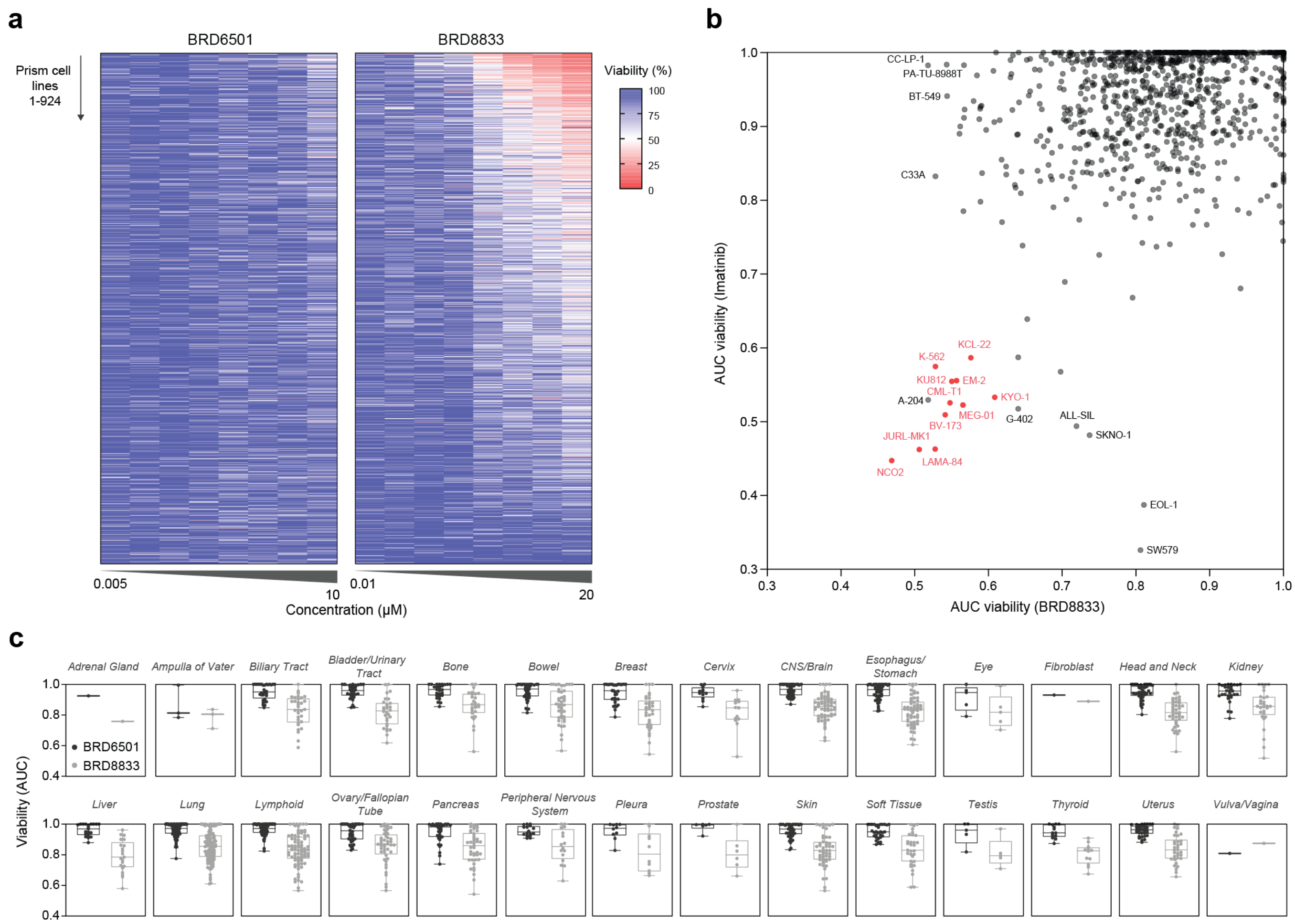


**Figure S7. PRISM screen demonstrates selective targeting of BCR-ABL dependent cell lines. a** Heatmap showing the sensitivity of cell lines in the PRISM screen after 5 days of treatment with control ABL binder BRD6501 or BRD8833. **b** Scatterplot comparing the cell line sensitivity to imatinib (y-axis) and BRD8833 (x-axis). BCR-ABL positive cell lines are highlighted in red, and all cell lines with AUC < 0.55 in either imatinib- or BRD8833-treated selection are annotated. **c** Boxplot showing drug sensitivity grouped by cell lineages. The box extends from the first to the third quartiles, the median is depicted as the center line, and the whiskers depict the range. Drug treatment was performed in three independent replicates (n = 3).

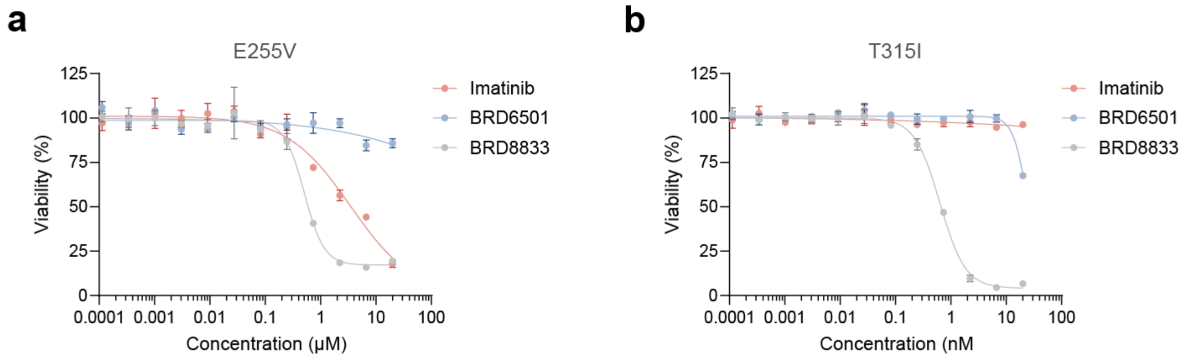


**Figure S8. BRD8833 targets imatinib-resistant BCR-ABL mutations.** Cytotoxicity in Ba/f3 stable cell lines expressing BCR-ABL carrying E255V (**a**) or gatekeeper T315I mutations (**b**). Mean and SD of three independent replicates are shown.

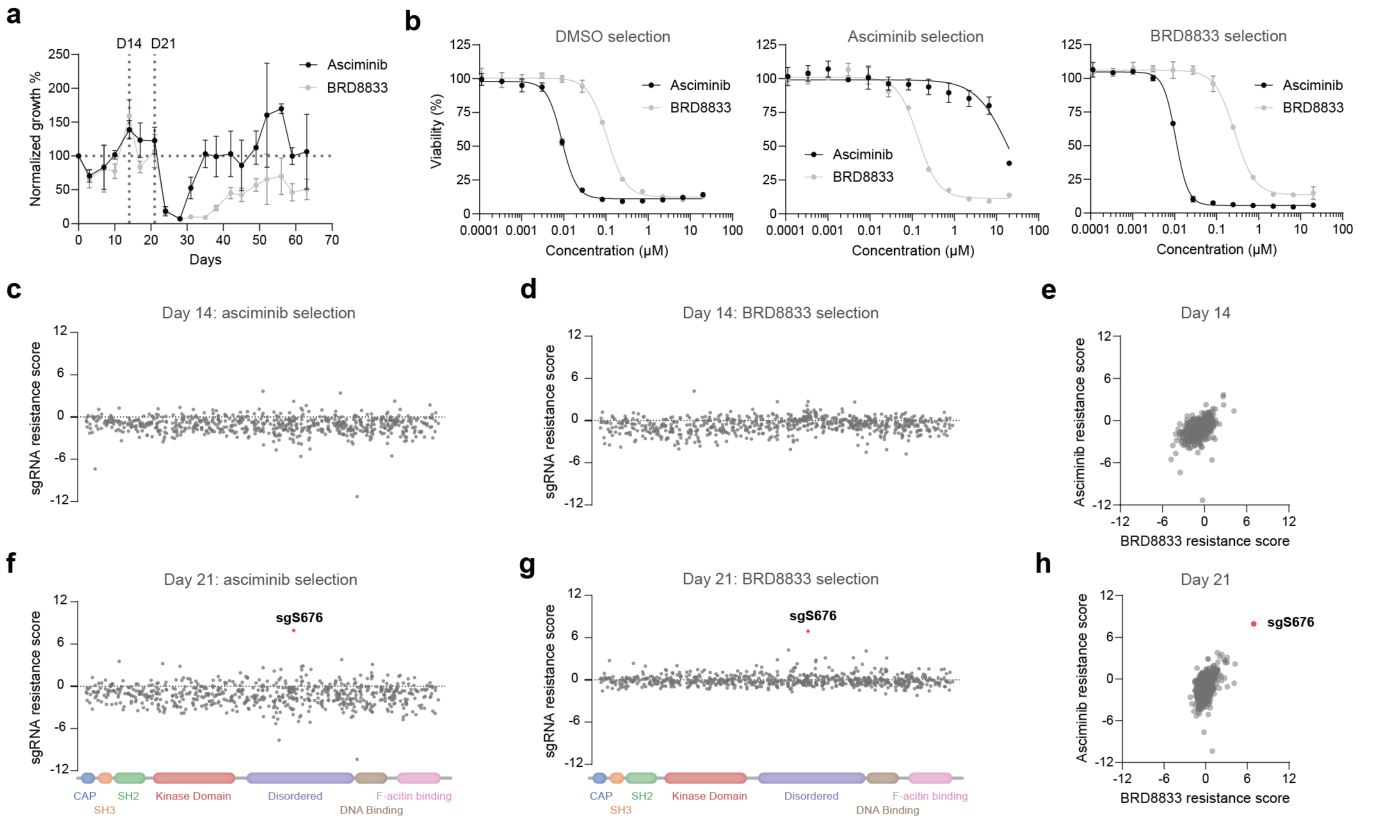


**Figure S9. CRISPR-suppressor scanning identifies mutant K562 cell lines. a** Growth rates during the selection process. Drug dose was increased, and cell samples were harvested for analysis of sgRNA enrichment after D14 and D21 of selection. Mean and SD for three independent selections are shown. **b** Dose-response for cytotoxicity in K562 cells subjected to a 63-day treatment with vehicle (DMSO), asciminib, or BRD8833. Mean and SD for three independent replicates are shown. **c–h** sgRNA enrichment after 14 days (**c–e**) or 21 days (**f–h**) of drug selection. Resistance scores were calculated as z-scored $\log_2$-fold changes in sgRNA abundance in drug-treated conditions relative to the vehicle (DMSO)-treated population. The scatter plots compare enrichment after 14 days (**e**) and 21 days (**h**) of selection with asciminib or BRD8833.

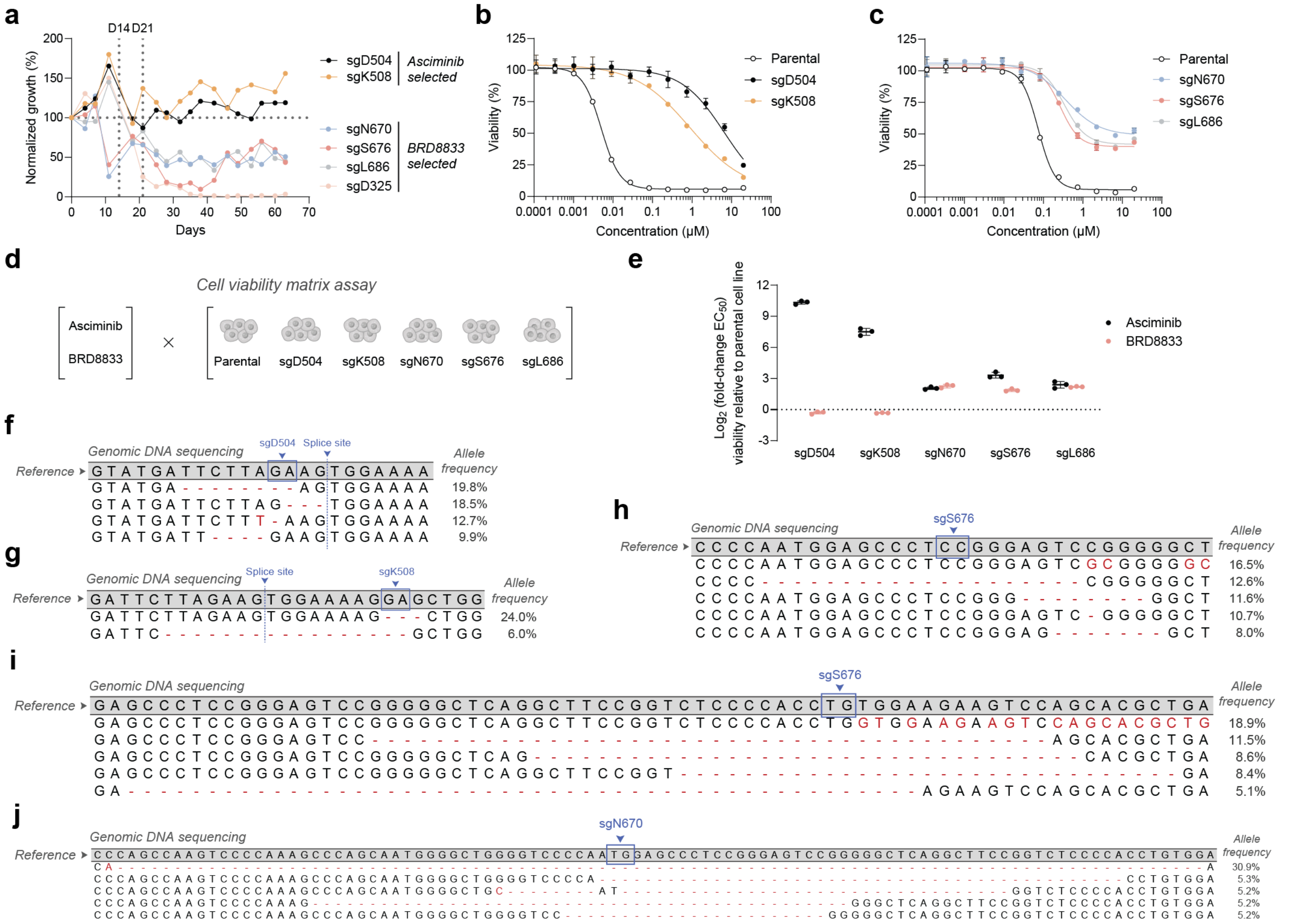


**Figure S10. CRISPR-suppressor scanning reveals different resistance mechanisms for asciminib and BRD8833. a** Growth rates of cells transduced with individual sgRNAs. Dose was increased after D14 and D21 of drug-selection as described in Table S2. **b–c** Cytotoxicity in K562 cells transduced with individual sgRNAs after 63 days of selection with asciminib (**b**) or BRD8833 (**c**). Cells were treated with their respective selection drug. Mean and SD of three independent replicates are shown. **d–e** Matrix cell viability assay to measure the drug response of K562 cells individually transduced with sgRNAs after 63 days of drug selection. Resistance is expressed as $\log_2$-fold change of an $EC_{50}$ versus the parental K562 cell line (**e**). Mean and SD of three independent replicates are shown. **f–j** Sequencing of genomic DNA. Cells were collected after 63 days of drug selection and sequenced near the sgRNA-targeted genomic loci.

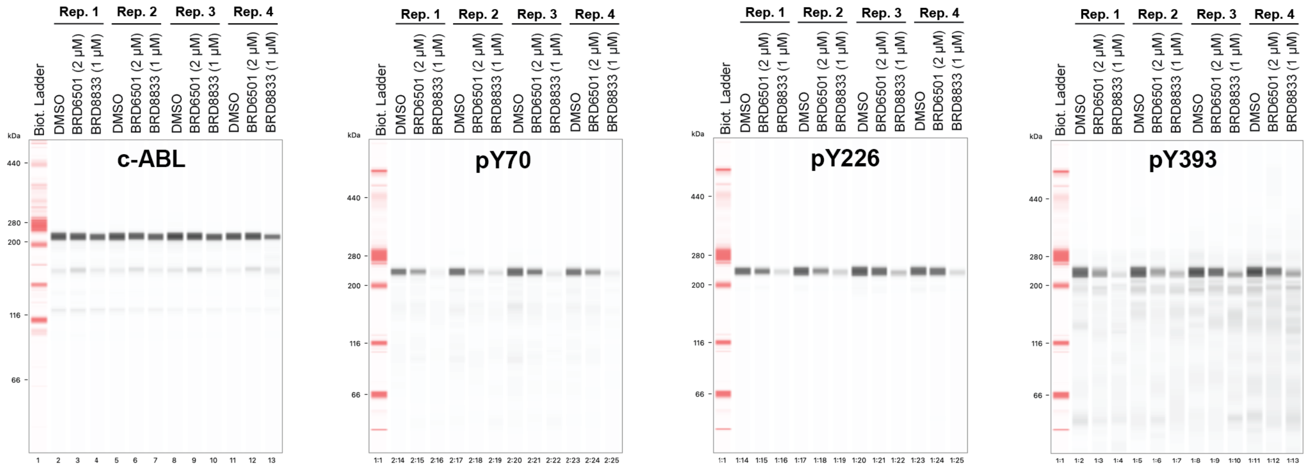


**Figure S11. Uncropped western blots used in Fig. 2f.** Full scans for Simple Western (JESS) shown in Fig. 2f. Each immunoblotting experiment was performed in four independent replicates. BCR-ABL phosphosites were mapped to the human ABL1 sequence (UniProt accession: P00519).

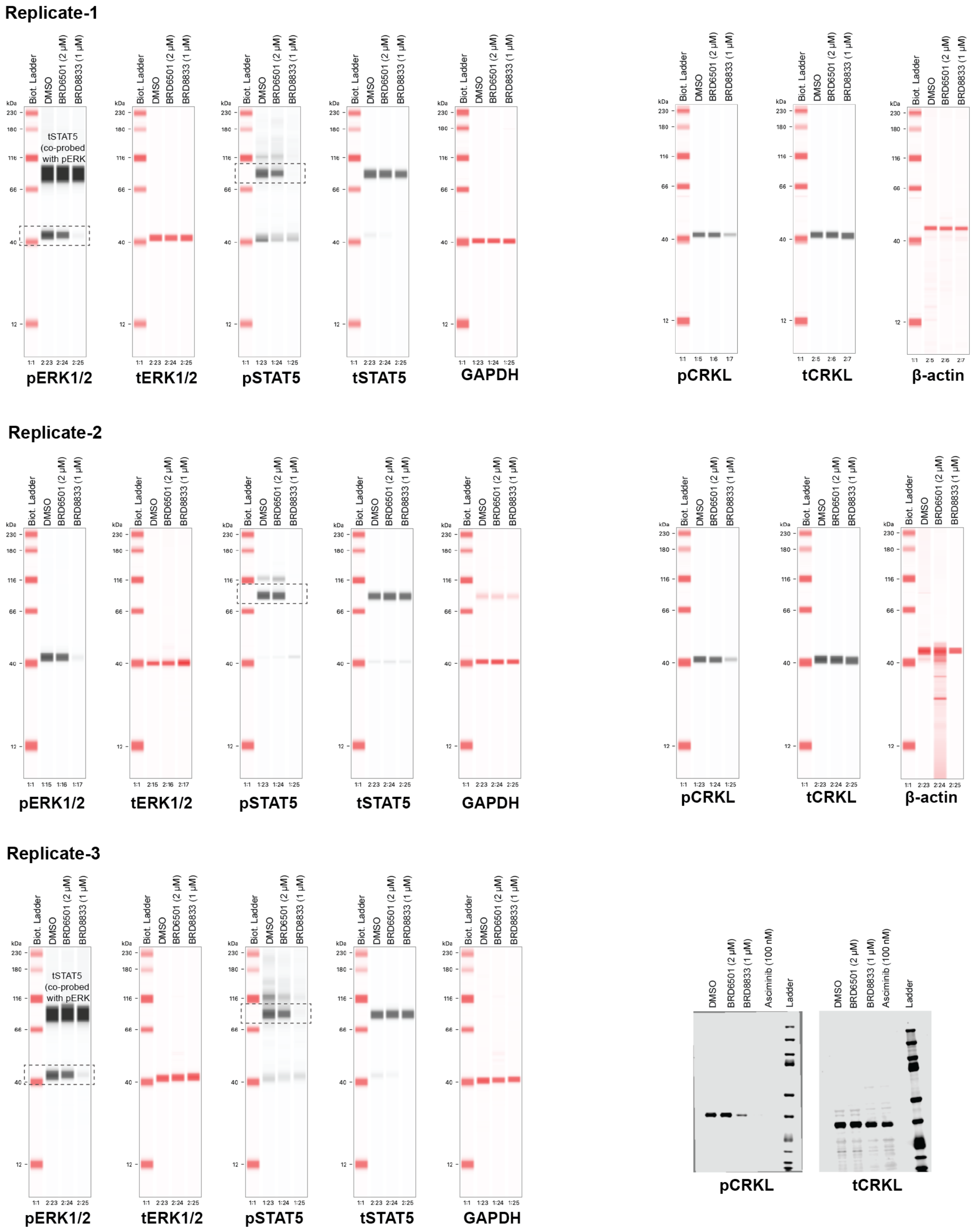


**Figure S12. Uncropped western blots used in Fig. 4a.** Full scans for Western blots and Simple Western (JESS) shown in Fig. 4a. Each immunoblotting experiment was performed in four independent replicates. Asciminib was included as a reference. BioRad Ref. #1610377 was used as a ladder for Western Blots.

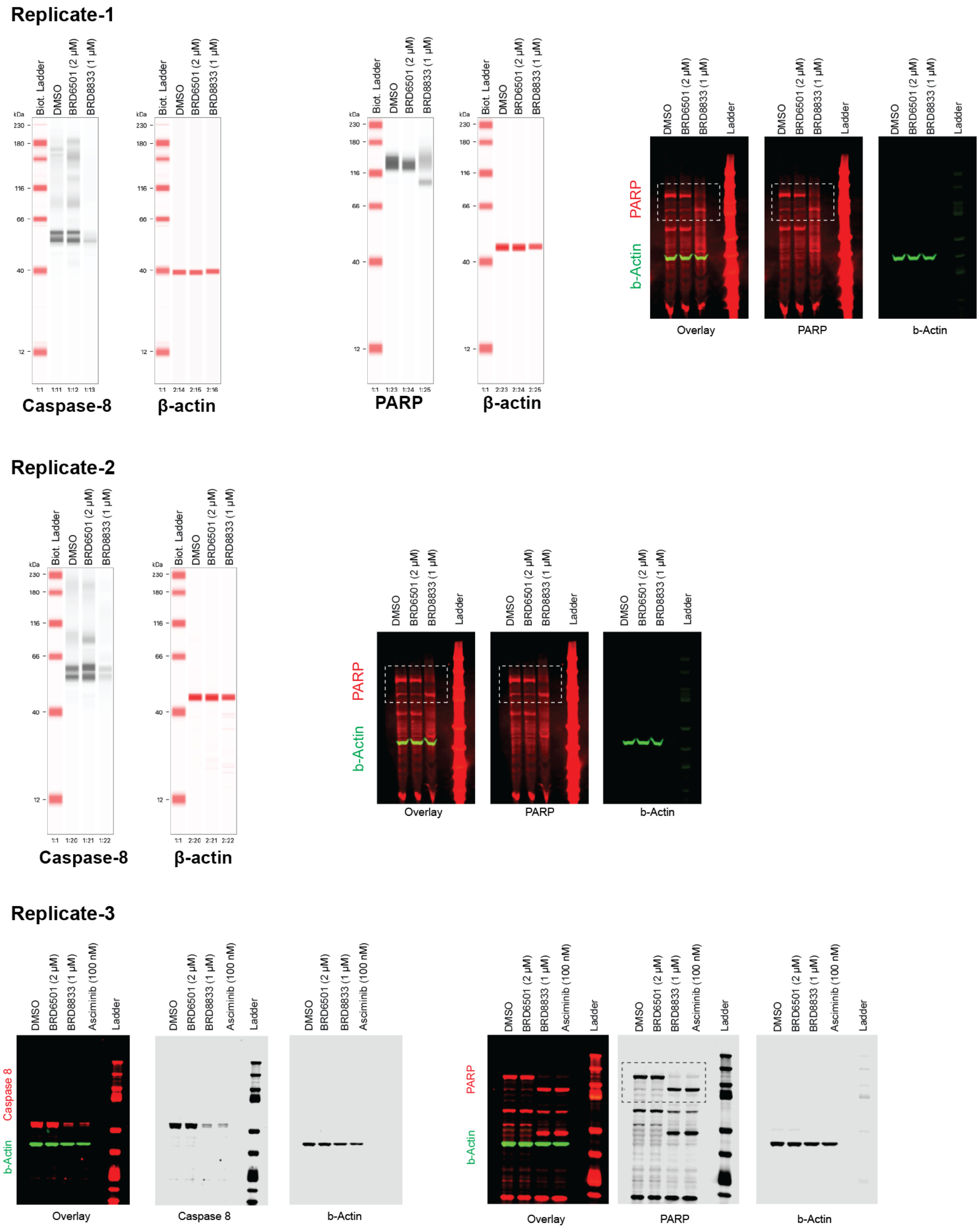


**Figure S13. Uncropped western blots used in Fig. 4b.** Full scans for Western blots and Simple Western (JESS) shown in Fig. 4b. Each immunoblotting experiment was performed in three independent replicates. Asciminib was included as a reference. BioRad Ref. #1610377 was used as a ladder for Western Blots.

**Table S1. LgBt and SmBt constructs to measure ternary complex formation.** *BCR-ABL1* genes fused N-terminally to either LgBt or SmBt were ordered from Genscript and cloned into pCDNA3.1 plasmid vector. Blue sequences correspond to *BCR-ABL1*, green sequence to *LgBt,* orange sequence to *SmBt*, and black sequences to spacers.

*BCR-ABL1 LgBt sequence*

ATGGAAGTCTTCACACTCGAAGATTTCGTTGGGGACTGGGAACAGACAGCCGCCTACAACCTGGACCAAGTCCTTGAACAGGGAGGTGTGTCCAGTTTGCTGCAGAAT
CTCGCCGTGTCCGTAACTCCGATCCAAAGGATTGTCCGGAGCGGTGAAAATGCCCTGAAGATCGACATCCATGTCATCATCCCGTATGAAGGTCTGAGCGCCGACCAA
ATGGCCCAGATCGAAGAGGTGTTTAAGGTGGTGTACCCTGTGGATGATCATCACTTTAAGGTGATCCTGCCCTATGGCACACTGGTAATCGACGGGGTTACGCCGAAC
ATGCTGAACTATTTCGGACGGCCGTATGAAGGCATCGCCGTGTTCGACGGCAAAAAGATCACTGTAACAGGGACCCTGTGGAACGGCAACAAAATTATCGACGAGCG
CCTGATCACCCCCGACGGCTCCATGCTGTTCCGAGTAACCATCAACAGTGGGAGTTCCGGTGGTGGCGGGAGCGGAGGTGGAGGCTCGAGCGGTGGAGCTCAGGG
GAATGTGGACCCGGTGGGCTTCGCGGAGGCGTGGAAGGCGCAGTTCCCGGACTCAGAGCCCCCGCGCATGGAGCTGCGCTCAGTGGGCGACATCGAGCAGGAGCT
GGAGCGCTGCAAGGCCTCCATTCGGCGCCTGGAGCAGGAGGTGAACCAGGAGCGCTTCCGCATGATCTACCTGCAGACGTTGCTGGCCAAGGAAAAGAAGAGCTAT
GACCGGCAGCGATGGGGCTTCCGGCGCGCGGCGCAGGCCCCCGACGGCGCCTCCGAGCCCCGAGCGTCCGCGTCGCGCCCGCAGCCAGCGCCCGCCGACGGAG
CCGACCCGCCGCCCGCCGAGGAGCCCGAGGCCCGGCCCGACGGCGAGGGTTCTCCGGGTAAGGCCAGGCCCGGGACCGCCCGCAGGCCCGGGGCAGCCGCGTC
GGGGGAACGGGACGACCGGGGACCCCCCGCCAGCGTGGCGGCGCTCAGGTCCAACTTCGAGCGGATCCGCAAGGGCCATGGCCAGCCCGGGGCGGACGCCGAG
AAGCCCTTCTACGTGAACGTCGAGTTTCACCACGAGCGCGGCCTGGTGAAGGTCAACGACAAAGAGGTGTCGGACCGCATCAGCTCCCTGGGCAGCCAGGCCATGC
AGATGGAGCGCAAAAAGTCCCAGCACGGCGCGGGCTCGAGCGTGGGGGATGCATCCAGGCCCCCTTACCGGGGACGCTCCTCGGAGAGCAGCTGCGGCGTCGAC
GGCGACTACGAGGACGCCGAGTTGAACCCCCGCTTCCTGAAGGACAACCTGATCGACGCCAATGGCGGTAGCAGGCCCCCTTGGCCGCCCCTGGAGTACCAGCCCT
ACCAGAGCATCTACGTCGGGGGCATGATGGAAGGGGAGGGCAAGGGCCCGCTCCTGCGCAGCCAGAGCACCTCTGAGCAGGAGAAGCGCCTTACCTGGCCCCGCA
GGTCCTACTCCCCCCGGAGTTTTGAGGATTGCGGAGGCGGCTATACCCCGGACTGCAGCTCCAATGAGAACCTCACCTCCAGCGAGGAGGACTTCTCCTCTGGCCAG
TCCAGCCGCGTGTCCCCAAGCCCCACCACCTACCGCATGTTCCGGGACAAAAGCCGCTCTCCCTCGCAGAACTCGCAACAGTCCTTCGACAGCAGCAGTCCCCCCAC
GCCGCAGTGCCATAAGCGGCACCGGCACTGCCCGGTTGTCGTGTCCGAGGCCACCATCGTGGGCGTCCGCAAGACCGGGCAGATCTGGCCCAACGATGGCGAGGG
CGCCTTCCATGGAGACGCAGATGGCTCGTTCGGAACACCACCTGGATACGGCTGCGCTGCAGACCGGGCAGAGGAGCAGCGCCGGCACCAAGATGGGCTGCCCTA
CATTGATGACTCGCCCTCCTCATCGCCCCACCTCAGCAGCAAGGGCAGGGGCAGCCGGGATGCGCTGGTCTCGGGAGCCCTGGAGTCCACTAAAGCGAGTGAGCTG
GACTTGGAAAAGGGCTTGGAGATGAGAAAATGGGTCCTGTCGGGAATCCTGGCTAGCGAGGAGACTTACCTGAGCCACCTGGAGGCACTGCTGCTGCCCATGAAGC
CTTTGAAAGCCGCTGCCACCACCTCTCAGCCGGTGCTGACGAGTCAGCAGATCGAGACCATCTTCTTCAAAGTGCCTGAGCTCTACGAGATCCACAAGGAGTTCTATG
ATGGGCTCTTCCCCCGCGTGCAGCAGTGGAGCCACCAGCAGCGGGTGGGCGACCTCTTCCAGAAGCTGGCCAGCCAGCTGGGTGTGTACCGGGCCTTCGTGGACA
ACTACGGAGTTGCCATGGAAATGGCTGAGAAGTGCTGTCAGGCCAATGCTCAGTTTGCAGAAATCTCCGAGAACCTGAGAGCCAGAAGCAACAAAGATGCCAAGGAT
CCAACGACCAAGAACTCTCTGGAAACTCTGCTCTACAAGCCTGTGGACCGTGTGACGAGGAGCACGCTGGTCCTCCATGACTTGCTGAAGCACACTCCTGCCAGCCA
CCCTGACCACCCCTTGCTGCAGGACGCCCTCCGCATCTCACAGAACTTCCTGTCCAGCATCAATGAGGAGATCACACCCCGACGGCAGTCCATGACGGTGAAGAAGG
GAGAGCACCGGCAGCTGCTGAAGGACAGCTTCATGGTGGAGCTGGTGGAGGGGGCCCGCAAGCTGCGCCACGTCTTCCTGTTCACCGACCTGCTTCTCTGCACCAA
GCTCAAGAAGCAGAGCGGAGGCAAAACGCAGCAGTATGACTGCAAATGGTACATTCCGCTCACGGATCTCAGCTTCCAGATGGTGGATGAACTGGAGGCAGTGCCCA
ACATCCCCCTGGTGCCCGATGAGGAGCTGGACGCTTTGAAGATCAAGATCTCCCAGATCAAGAGTGACATCCAGAGAGAGAAGAGGGCGAACAAGGGCAGCAAGGC
TACGGAGAGGCTGAAGAAGAAGCTGTCGGAGCAGGAGTCACTGCTGCTGCTTATGTCTCCCAGCATGGCCTTCAGGGTGCACAGCCGCAACGGCAAGAGTTACACGT
TCCTGATCTCCTCTGACTATGAGCGTGCAGAGTGGAGGGAGAACATCCGGGAGCAGCAGAAGAAGTGTTTCAGAAGCTTCTCCCTGACATCCGTGGAGCTGCAGATG
CTGACCAACTCGTGTGTGAAACTCCAGACTGTCCACAGCATTCCGCTGACCATCAATAAGGAAGATGATGAGTCTCCGGGGCTCTATGGGTTTCTGAATGTCATCGTC
CACTCAGCCACTGGATTTAAGCAGAGTTCAAAAGCCCTTCAGCGGCCAGTAGCATCTGACTTTGAGCCTCAGGGTCTGAGTGAAGCCGCTCGTTGGAACTCCAAGGAA
AACCTTCTCGCTGGACCCAGTGAAAATGACCCCAACCTTTTCGTTGCACTGTATGATTTTGTGGCCAGTGGAGATAACACTCTAAGCATAACTAAAGGTGAAAAGCTCC
GGGTCTTAGGCTATAATCACAATGGGGAATGGTGTGAAGCCCAAACCAAAAATGGCCAAGGCTGGGTCCCAAGCAACTACATCACGCCAGTCAACAGTCTGGAGAAAC
ACTCCTGGTACCATGGGCCTGTGTCCCGCAATGCCGCTGAGTATCTGCTGAGCAGCGGGATCAATGGCAGCTTCTTGGTGCGTGAGAGTGAGAGCAGTCCTGGCCAG
AGGTCCATCTCGCTGAGATACGAAGGGAGGGTGTACCATTACAGGATCAACACTGCTTCTGATGGCAAGCTCTACGTCTCCTCCGAGAGCCGCTTCAACACCCTGGCC
GAGTTGGTTCATCATCATTCAACGGTGGCCGACGGGCTCATCACCACGCTCCATTATCCAGCCCCAAAGCGCAACAAGCCCACTGTCTATGGTGTGTCCCCCAACTAC
GACAAGTGGGAGATGGAACGCACGGACATCACCATGAAGCACAAGCTGGGCGGGGGCCAGTACGGGGAGGTGTACGAGGGCGTGTGGAAGAAATACAGCCTGACG
GTGGCCGTGAAGACCTTGAAGGAGGACACCATGGAGGTGGAAGAGTTCTTGAAAGAAGCTGCAGTCATGAAAGAGATCAAACACCCTAACCTGGTGCAGCTCCTTGG
GGTCTGCACCCGGGAGCCCCCGTTCTATATCATCACTGAGTTCATGACCTACGGGAACCTCCTGGACTACCTGAGGGAGTGCAACCGGCAGGAGGTGAACGCCGTG
GTGCTGCTGTACATGGCCACTCAGATCTCGTCAGCCATGGAGTACCTGGAGAAGAAAAACTTCATCCACAGAGATCTTGCTGCCCGAAACTGCCTGGTAGGGGAGAAC
CACTTGGTGAAGGTAGCTGATTTTGGCCTGAGCAGGTTGATGACAGGGGACACCTACACAGCCCATGCTGGAGCCAAGTTCCCCATCAAATGGACTGCACCCGAGAG
CCTGGCCTACAACAAGTTCTCCATCAAGTCCGACGTCTGGGCATTTGGAGTATTGCTTTGGGAAATTGCTACCTATGGCATGTCCCCTTACCCGGGAATTGACCTGTCC
CAGGTGTATGAGCTGCTAGAGAAGGACTACCGCATGGAGCGCCCAGAAGGCTGCCCAGAGAAGGTCTATGAACTCATGCGAGCATGTTGGCAGTGGAATCCCTCTGA
CCGGCCCTCCTTTGCTGAAATCCACCAAGCCTTTGAAACAATGTTCCAGGAATCCAGTATCTCAGACGAAGTGGAAAAGGAGCTGGGGAAACAAGGCGTCCGTGGGG
CTGTGAGTACCTTGCTGCAGGCCCCAGAGCTGCCCACCAAGACGAGGACCTCCAGGAGAGCTGCAGAGCACAGAGACACCACTGACGTGCCTGAGATGCCTCACTC
CAAGGGCCAGGGAGAGAGCGATCCTCTGGACCATGAGCCTGCCGTGTCTCCATTGCTCCCTCGAAAAGAGCGAGGTCCCCCGGAGGGCGGCCTGAATGAAGATGAG
CGCCTTCTCCCCAAAGACAAAAAGACCAACTTGTTCAGCGCCTTGATCAAGAAGAAGAAGAAGACAGCCCCAACCCCTCCCAAACGCAGCAGCTCCTTCCGGGAGATG
GACGGCCAGCCGGAGCGCAGAGGGGCCGGCGAGGAAGAGGGCCGAGACATCAGCAACGGGGCACTGGCTTTCACCCCCTTGGACACAGCTGACCCAGCCAAGTCC
CCAAAGCCCAGCAATGGGGCTGGGGTCCCCAATGGAGCCCTCCGGGAGTCCGGGGGCTCAGGCTTCCGGTCTCCCCACCTGTGGAAGAAGTCCAGCACGCTGACC
AGCAGCCGCCTAGCCACCGGCGAGGAGGAGGGCGGTGGCAGCTCCAGCAAGCGCTTCCTGCGCTCTTGCTCCGCCTCCTGCGTTCCCCATGGGGCCAAGGACACG
GAGTGGAGGTCAGTCACGCTGCCTCGGGACTTGCAGTCCACGGGAAGACAGTTTGACTCGTCCACATTTGGAGGGCACAAAAGTGAGAAGCCGGCTCTGCCTCGGA
AGAGGGCAGGGGAGAACAGGTCTGACCAGGTGACCCGAGGCACAGTAACGCCTCCCCCCAGGCTGGTGAAAAAGAATGAGGAAGCTGCTGATGAGGTCTTCAAAGA
CATCATGGAGTCCAGCCCGGGCTCCAGCCCGCCCAACCTGACTCCAAAACCCCTCCGGCGGCAGGTCACCGTGGCCCCTGCCTCGGGCCTCCCCCACAAGGAAGAA
GCTGGAAAGGGCAGTGCCTTAGGGACCCCTGCTGCAGCTGAGCCAGTGACCCCCACCAGCAAAGCAGGCTCAGGTGCACCAGGGGGCACCAGCAAGGGCCCCGCC
GAGGAGTCCAGAGTGAGGAGGCACAAGCACTCCTCTGAGTCGCCAGGGAGGGACAAGGGGAAATTGTCCAGGCTCAAACCTGCCCCGCCGCCCCCACCAGCAGCC
TCTGCAGGGAAGGCTGGAGGAAAGCCCTCGCAGAGCCCGAGCCAGGAGGCGGCCGGGGAGGCAGTCCTGGGCGCAAAGACAAAAGCCACGAGTCTGGTTGATGCT
GTGAACAGTGACGCTGCCAAGCCCAGCCAGCCGGGAGAGGGCCTCAAAAAGCCCGTGCTCCCGGCCACTCCAAAGCCACAGTCCGCCAAGCCGTCGGGGACCCCC
ATCAGCCCAGCCCCCGTTCCCTCCACGTTGCCATCAGCATCCTCGGCCCTGGCAGGGGACCAGCCGTCTTCCACCGCCTTCATCCCTCTCATATCAACCCGAGTGTC
TCTTCGGAAAACCCGCCAGCCTCCAGAGCGGATCGCCAGCGGCGCCATCACCAAGGGCGTGGTCCTGGACAGCACCGAGGCGCTGTGCCTCGCCATCTCTAGGAAC
TCCGAGCAGATGGCCAGCCACAGCGCAGTGCTGGAGGCCGGCAAAAACCTCTACTCGTTCTGCGTGAGCTATGTGGATTCCATCCAGCAAATGAGGAACAAGTTTGC
CTTCCGAGAGGCCATCAACAAACTGGAGAATAATCTCCGGGAGCTTCAGATCTGCCCGGCGACAGCAGGCAGTGGTCCAGCGGCCACTCAGGACTTCAGCAAGCTCC
TCAGTTCGGTGAAGGAAATCAGTGACATAGTGCAGAGG

#### *BCR-ABL1 SmBt sequence*

ATGGAAGTGACCGGCTACCGGCTGTTCGAGGAGATTCTGGGGAGTTCCGGTGGTGGCGGGAGCGGAGGTGGAGGCTCGAGCGGTGGAGCTCAGGGGAATGTGGAC
CCGGTGGGCTTCGCGGAGGCGTGGAAGGCGCAGTTCCCGGACTCAGAGCCCCCGCGCATGGAGCTGCGCTCAGTGGGCGACATCGAGCAGGAGCTGGAGCGCTG
CAAGGCCTCCATTCGGCGCCTGGAGCAGGAGGTGAACCAGGAGCGCTTCCGCATGATCTACCTGCAGACGTTGCTGGCCAAGGAAAAGAAGAGCTATGACCGGCAG
CGATGGGGCTTCCGGCGCGCGGCGCAGGCCCCCGACGGCGCCTCCGAGCCCCGAGCGTCCGCGTCGCGCCCGCAGCCAGCGCCCGCCGACGGAGCCGACCCGC
CGCCCGCCGAGGAGCCCGAGGCCCGGCCCGACGGCGAGGGTTCTCCGGGTAAGGCCAGGCCCGGGACCGCCCGCAGGCCCGGGGCAGCCGCGTCGGGGGAAC
GGGACGACCGGGGACCCCCCGCCAGCGTGGCGGCGCTCAGGTCCAACTTCGAGCGGATCCGCAAGGGCCATGGCCAGCCCGGGGCGGACGCCGAGAAGCCCTTC
TACGTGAACGTCGAGTTTCACCACGAGCGCGGCCTGGTGAAGGTCAACGACAAAGAGGTGTCGGACCGCATCAGCTCCCTGGGCAGCCAGGCCATGCAGATGGAGC
GCAAAAAGTCCCAGCACGGCGCGGGCTCGAGCGTGGGGGATGCATCCAGGCCCCCTTACCGGGGACGCTCCTCGGAGAGCAGCTGCGGCGTCGACGGCGACTAC
GAGGACGCCGAGTTGAACCCCCGCTTCCTGAAGGACAACCTGATCGACGCCAATGGCGGTAGCAGGCCCCCTTGGCCGCCCCTGGAGTACCAGCCCTACCAGAGCA
TCTACGTCGGGGGCATGATGGAAGGGGAGGGCAAGGGCCCGCTCCTGCGCAGCCAGAGCACCTCTGAGCAGGAGAAGCGCCTTACCTGGCCCCGCAGGTCCTACT
CCCCCCGGAGTTTTGAGGATTGCGGAGGCGGCTATACCCCGGACTGCAGCTCCAATGAGAACCTCACCTCCAGCGAGGAGGACTTCTCCTCTGGCCAGTCCAGCCG
CGTGTCCCCAAGCCCCACCACCTACCGCATGTTCCGGGACAAAAGCCGCTCTCCCTCGCAGAACTCGCAACAGTCCTTCGACAGCAGCAGTCCCCCCACGCCGCAGT
GCCATAAGCGGCACCGGCACTGCCCGGTTGTCGTGTCCGAGGCCACCATCGTGGGCGTCCGCAAGACCGGGCAGATCTGGCCCAACGATGGCGAGGGCGCCTTCC
ATGGAGACGCAGATGGCTCGTTCGGAACACCACCTGGATACGGCTGCGCTGCAGACCGGGCAGAGGAGCAGCGCCGGCACCAAGATGGGCTGCCCTACATTGATGA
CTCGCCCTCCTCATCGCCCCACCTCAGCAGCAAGGGCAGGGGCAGCCGGGATGCGCTGGTCTCGGGAGCCCTGGAGTCCACTAAAGCGAGTGAGCTGGACTTGGA
AAAGGGCTTGGAGATGAGAAAATGGGTCCTGTCGGGAATCCTGGCTAGCGAGGAGACTTACCTGAGCCACCTGGAGGCACTGCTGCTGCCCATGAAGCCTTTGAAAG
CCGCTGCCACCACCTCTCAGCCGGTGCTGACGAGTCAGCAGATCGAGACCATCTTCTTCAAAGTGCCTGAGCTCTACGAGATCCACAAGGAGTTCTATGATGGGCTCT
TCCCCCGCGTGCAGCAGTGGAGCCACCAGCAGCGGGTGGGCGACCTCTTCCAGAAGCTGGCCAGCCAGCTGGGTGTGTACCGGGCCTTCGTGGACAACTACGGAG
TTGCCATGGAAATGGCTGAGAAGTGCTGTCAGGCCAATGCTCAGTTTGCAGAAATCTCCGAGAACCTGAGAGCCAGAAGCAACAAAGATGCCAAGGATCCAACGACCA
AGAACTCTCTGGAAACTCTGCTCTACAAGCCTGTGGACCGTGTGACGAGGAGCACGCTGGTCCTCCATGACTTGCTGAAGCACACTCCTGCCAGCCACCCTGACCAC
CCCTTGCTGCAGGACGCCCTCCGCATCTCACAGAACTTCCTGTCCAGCATCAATGAGGAGATCACACCCCGACGGCAGTCCATGACGGTGAAGAAGGGAGAGCACCG
GCAGCTGCTGAAGGACAGCTTCATGGTGGAGCTGGTGGAGGGGGCCCGCAAGCTGCGCCACGTCTTCCTGTTCACCGACCTGCTTCTCTGCACCAAGCTCAAGAAG
CAGAGCGGAGGCAAAACGCAGCAGTATGACTGCAAATGGTACATTCCGCTCACGGATCTCAGCTTCCAGATGGTGGATGAACTGGAGGCAGTGCCCAACATCCCCCT
GGTGCCCGATGAGGAGCTGGACGCTTTGAAGATCAAGATCTCCCAGATCAAGAGTGACATCCAGAGAGAGAAGAGGGCGAACAAGGGCAGCAAGGCTACGGAGAGG
CTGAAGAAGAAGCTGTCGGAGCAGGAGTCACTGCTGCTGCTTATGTCTCCCAGCATGGCCTTCAGGGTGCACAGCCGCAACGGCAAGAGTTACACGTTCCTGATCTC
CTCTGACTATGAGCGTGCAGAGTGGAGGGAGAACATCCGGGAGCAGCAGAAGAAGTGTTTCAGAAGCTTCTCCCTGACATCCGTGGAGCTGCAGATGCTGACCAACT
CGTGTGTGAAACTCCAGACTGTCCACAGCATTCCGCTGACCATCAATAAGGAAGATGATGAGTCTCCGGGGCTCTATGGGTTTCTGAATGTCATCGTCCACTCAGCCA
CTGGATTTAAGCAGAGTTCAAAAGCCCTTCAGCGGCCAGTAGCATCTGACTTTGAGCCTCAGGGTCTGAGTGAAGCCGCTCGTTGGAACTCCAAGGAAAACCTTCTCG
CTGGACCCAGTGAAAATGACCCCAACCTTTTCGTTGCACTGTATGATTTTGTGGCCAGTGGAGATAACACTCTAAGCATAACTAAAGGTGAAAAGCTCCGGGTCTTAGG
CTATAATCACAATGGGGAATGGTGTGAAGCCCAAACCAAAAATGGCCAAGGCTGGGTCCCAAGCAACTACATCACGCCAGTCAACAGTCTGGAGAAACACTCCTGGTA
CCATGGGCCTGTGTCCCGCAATGCCGCTGAGTATCTGCTGAGCAGCGGGATCAATGGCAGCTTCTTGGTGCGTGAGAGTGAGAGCAGTCCTGGCCAGAGGTCCATCT
CGCTGAGATACGAAGGGAGGGTGTACCATTACAGGATCAACACTGCTTCTGATGGCAAGCTCTACGTCTCCTCCGAGAGCCGCTTCAACACCCTGGCCGAGTTGGTT
CATCATCATTCAACGGTGGCCGACGGGCTCATCACCACGCTCCATTATCCAGCCCCAAAGCGCAACAAGCCCACTGTCTATGGTGTGTCCCCCAACTACGACAAGTGG
GAGATGGAACGCACGGACATCACCATGAAGCACAAGCTGGGCGGGGGCCAGTACGGGGAGGTGTACGAGGGCGTGTGGAAGAAATACAGCCTGACGGTGGCCGTG
AAGACCTTGAAGGAGGACACCATGGAGGTGGAAGAGTTCTTGAAAGAAGCTGCAGTCATGAAAGAGATCAAACACCCTAACCTGGTGCAGCTCCTTGGGGTCTGCAC
CCGGGAGCCCCCGTTCTATATCATCACTGAGTTCATGACCTACGGGAACCTCCTGGACTACCTGAGGGAGTGCAACCGGCAGGAGGTGAACGCCGTGGTGCTGCTGT
ACATGGCCACTCAGATCTCGTCAGCCATGGAGTACCTGGAGAAGAAAAACTTCATCCACAGAGATCTTGCTGCCCGAAACTGCCTGGTAGGGGAGAACCACTTGGTGA
AGGTAGCTGATTTTGGCCTGAGCAGGTTGATGACAGGGGACACCTACACAGCCCATGCTGGAGCCAAGTTCCCCATCAAATGGACTGCACCCGAGAGCCTGGCCTAC
AACAAGTTCTCCATCAAGTCCGACGTCTGGGCATTTGGAGTATTGCTTTGGGAAATTGCTACCTATGGCATGTCCCCTTACCCGGGAATTGACCTGTCCCAGGTGTATG
AGCTGCTAGAGAAGGACTACCGCATGGAGCGCCCAGAAGGCTGCCCAGAGAAGGTCTATGAACTCATGCGAGCATGTTGGCAGTGGAATCCCTCTGACCGGCCCTC
CTTTGCTGAAATCCACCAAGCCTTTGAAACAATGTTCCAGGAATCCAGTATCTCAGACGAAGTGGAAAAGGAGCTGGGGAAACAAGGCGTCCGTGGGGCTGTGAGTAC
CTTGCTGCAGGCCCCAGAGCTGCCCACCAAGACGAGGACCTCCAGGAGAGCTGCAGAGCACAGAGACACCACTGACGTGCCTGAGATGCCTCACTCCAAGGGCCAG
GGAGAGAGCGATCCTCTGGACCATGAGCCTGCCGTGTCTCCATTGCTCCCTCGAAAAGAGCGAGGTCCCCCGGAGGGCGGCCTGAATGAAGATGAGCGCCTTCTCC
CCAAAGACAAAAAGACCAACTTGTTCAGCGCCTTGATCAAGAAGAAGAAGAAGACAGCCCCAACCCCTCCCAAACGCAGCAGCTCCTTCCGGGAGATGGACGGCCAG
CCGGAGCGCAGAGGGGCCGGCGAGGAAGAGGGCCGAGACATCAGCAACGGGGCACTGGCTTTCACCCCCTTGGACACAGCTGACCCAGCCAAGTCCCCAAAGCCC
AGCAATGGGGCTGGGGTCCCCAATGGAGCCCTCCGGGAGTCCGGGGGCTCAGGCTTCCGGTCTCCCCACCTGTGGAAGAAGTCCAGCACGCTGACCAGCAGCCGC
CTAGCCACCGGCGAGGAGGAGGGCGGTGGCAGCTCCAGCAAGCGCTTCCTGCGCTCTTGCTCCGCCTCCTGCGTTCCCCATGGGGCCAAGGACACGGAGTGGAGG
TCAGTCACGCTGCCTCGGGACTTGCAGTCCACGGGAAGACAGTTTGACTCGTCCACATTTGGAGGGCACAAAAGTGAGAAGCCGGCTCTGCCTCGGAAGAGGGCAG
GGGAGAACAGGTCTGACCAGGTGACCCGAGGCACAGTAACGCCTCCCCCCAGGCTGGTGAAAAAGAATGAGGAAGCTGCTGATGAGGTCTTCAAAGACATCATGGA
GTCCAGCCCGGGCTCCAGCCCGCCCAACCTGACTCCAAAACCCCTCCGGCGGCAGGTCACCGTGGCCCCTGCCTCGGGCCTCCCCCACAAGGAAGAAGCTGGAAA
GGGCAGTGCCTTAGGGACCCCTGCTGCAGCTGAGCCAGTGACCCCCACCAGCAAAGCAGGCTCAGGTGCACCAGGGGGCACCAGCAAGGGCCCCGCCGAGGAGTC
CAGAGTGAGGAGGCACAAGCACTCCTCTGAGTCGCCAGGGAGGGACAAGGGGAAATTGTCCAGGCTCAAACCTGCCCCGCCGCCCCCACCAGCAGCCTCTGCAGG
GAAGGCTGGAGGAAAGCCCTCGCAGAGCCCGAGCCAGGAGGCGGCCGGGGAGGCAGTCCTGGGCGCAAAGACAAAAGCCACGAGTCTGGTTGATGCTGTGAACAG
TGACGCTGCCAAGCCCAGCCAGCCGGGAGAGGGCCTCAAAAAGCCCGTGCTCCCGGCCACTCCAAAGCCACAGTCCGCCAAGCCGTCGGGGACCCCCATCAGCCC
AGCCCCCGTTCCCTCCACGTTGCCATCAGCATCCTCGGCCCTGGCAGGGGACCAGCCGTCTTCCACCGCCTTCATCCCTCTCATATCAACCCGAGTGTCTCTTCGGAA
AACCCGCCAGCCTCCAGAGCGGATCGCCAGCGGCGCCATCACCAAGGGCGTGGTCCTGGACAGCACCGAGGCGCTGTGCCTCGCCATCTCTAGGAACTCCGAGCA
GATGGCCAGCCACAGCGCAGTGCTGGAGGCCGGCAAAAACCTCTACTCGTTCTGCGTGAGCTATGTGGATTCCATCCAGCAAATGAGGAACAAGTTTGCCTTCCGAG
AGGCCATCAACAAACTGGAGAATAATCTCCGGGAGCTTCAGATCTGCCCGGCGACAGCAGGCAGTGGTCCAGCGGCCACTCAGGACTTCAGCAAGCTCCTCAGTTCG
GTGAAGGAAATCAGTGACATAGTGCAGAGG

**Table S2. Drug dosing regimen for CRISPR-suppressor scanning.** The CRISPR-suppressor scanning dose regimen was established based on the drug sensitivity of the K562 parental cell line. $EC_{50}$ values of 5 nM for asciminib and 69 nM for BRD8833 were used as reference points. The drug dose was gradually escalated, selecting first at the lowest dose for two weeks (days 0–14), then an intermediate dose for one week (days 14–21), and then the highest dose for six weeks (days 21–63). The selection at the highest dose was extended to allow BRD8833 treated cells to recover growth. The same treatment was used for selecting drug-resistant cells for sgRNA validation.

| Drug | Dose during the indicated treatment period (nM) | | |
|---|---|---|---|
| | Day 0 to 14 | Day 14 to 21 | Day 21 to 63 |
| Asciminib | 2 | 4 | 100 |
| BRD8833 | 40 | 80 | 500 |

**Table S3. Oligonucleotides used for sgRNA validation.** Single stranded oligonucleotides containing protospacer sequences with overhangs (ordered from Eton Bioscience) for cloning into Esp3I-digested lentiCRISPRv2 vector (Addgene #52961) are provided.

| sgRNA target residue | Target strand | Protospacer | Oligonucleotide Sequence (5' → 3') | |
|---|---|---|---|---|
| | | | Forward | Reverse |
| D325 | - | TGCACTCCCTCAGGTAGTCC | CACCGTGCACTCCCTCAGGTAGTCC | CGGACTACCTGAGGGAGTGCACAAA |
| D504 | + | CCCTGTATGATTCTTAGAAG | CACCGCCCTGTATGATTCTTAGAAG | CCTTCTAAGAATCATACAGGGCAAA |
| K508 | + | CTTAGAAGTGGAAAAGGAGC | CACCGCTTAGAAGTGGAAAAGGAGC | CGCTCCTTTTCCACTTCTAAGCAAA |
| N670 | - | GACTCCCGGAGGGCTCCATT | CACCGGACTCCCGGAGGGCTCCATT | CAATGGAGCCCTCCGGGAGTCCAAA |
| S676 | + | GGAGCCCTCCGGGAGTCCGG | CACCGGGAGCCCTCCGGGAGTCCGG | CCCGGACTCCCGGAGGGCTCCCAAA |
| L686 | - | TGCTGGACTTCTTCCACAGG | CACCGTGCTGGACTTCTTCCACAGG | CCCTGTGGAAGAAGTCCAGCACAAA |

**Table S4. Primers used for indel sequencing.** Two rounds of PCR were used for amplicon sequencing. For the first round, gene-specific primers (sequences in green) with common overhangs were used to synthesize amplicons from template gDNA or cDNA. The second PCR round was used to append Illumina adapter sequences with dual 8-nt indexes (sequences in blue) for NGS sample multiplexing.

| Amplicon description | PCR round | Template | Direction | Oligonucleotide Sequence (5' → 3') |
|---|---|---|---|---|
| sgRNA# 504,506 | 1 | gDNA | Forward | TCGTCGGCAGCGTCAGATGTGTATAAGAGACAGCGGTATCCACGTGCCTTTTC |
| | | | Reverse | GTCTCGTGGGCTCGGAGATGTGTATAAGAGACAGGACTTACCGCTCTCTCCCTG |
| sgRNA# 504,506 | 1 | cDNA | Forward | TCGTCGGCAGCGTCAGATGTGTATAAGAGACAGCTGCCCAGAGAAGGTCTATG |
| | | | Reverse | GTCTCGTGGGCTCGGAGATGTGTATAAGAGACAGGACTTACCGCTCTCTCCCTG |
| sgRNA# 670,676,686 | 1 | gDNA | Forward | TCGTCGGCAGCGTCAGATGTGTATAAGAGACAGCTGGCTTTCACCCCCTTGGA |
| | | | Reverse | GTCTCGTGGGCTCGGAGATGTGTATAAGAGACAGGTCAAACTGTCTTCCCGTGG |
| sgRNA# 670,676,686 | 1 | cDNA | Forward | TCGTCGGCAGCGTCAGATGTGTATAAGAGACAGGAAGAGGGCCGAGACATCAG |
| | | | Reverse | GTCTCGTGGGCTCGGAGATGTGTATAAGAGACAGGTCAAACTGTCTTCCCGTGG |
| NGS Indexing | 2 | gDNA, cDNA | Forward | AATGATACGGCGACCACCGAGATCTACACTAGATCGCTCGTCGGCAGCGTC |
| | | gDNA, cDNA | Reverse | CAAGCAGAAGACGGCATACGAGATTCGCCTTAGTCTCGTGGGCTCGG |
| | | gDNA, cDNA | Forward | AATGATACGGCGACCACCGAGATCTACACCTCTCTATTCGTCGGCAGCGTC |
| | | gDNA, cDNA | Reverse | CAAGCAGAAGACGGCATACGAGATCTAGTACGGTCTCGTGGGCTCGG |
| | | gDNA, cDNA | Forward | AATGATACGGCGACCACCGAGATCTACACTATCCTCTTCGTCGGCAGCGTC |
| | | gDNA, cDNA | Reverse | CAAGCAGAAGACGGCATACGAGATTTCTGCCTGTCTCGTGGGCTCGG |
| | | gDNA, cDNA | Forward | AATGATACGGCGACCACCGAGATCTACACAGAGTAGATCGTCGGCAGCGTC |
| | | gDNA, cDNA | Reverse | CAAGCAGAAGACGGCATACGAGATGCTCAGGAGTCTCGTGGGCTCGG |
| | | gDNA, cDNA | Forward | AATGATACGGCGACCACCGAGATCTACACGTAAGGAGTCGTCGGCAGCGTC |
| | | gDNA, cDNA | Reverse | CAAGCAGAAGACGGCATACGAGATAGGAGTCCGTCTCGTGGGCTCGG |
| | | gDNA, cDNA | Forward | AATGATACGGCGACCACCGAGATCTACACACTGCATATCGTCGGCAGCGTC |
| | | gDNA, cDNA | Reverse | CAAGCAGAAGACGGCATACGAGATCATGCCTAGTCTCGTGGGCTCGG |
| | | gDNA, cDNA | Forward | AATGATACGGCGACCACCGAGATCTACACAAGGAGTATCGTCGGCAGCGTC |
| | | gDNA, cDNA | Reverse | CAAGCAGAAGACGGCATACGAGATGTAGAGAGGTCTCGTGGGCTCGG |
| | | gDNA, cDNA | Forward | AATGATACGGCGACCACCGAGATCTACACCTAAGCCTTCGTCGGCAGCGTC |
| | | gDNA, cDNA | Reverse | CAAGCAGAAGACGGCATACGAGATCCTCTCTGGTCTCGTGGGCTCGG |
| | | gDNA, cDNA | Forward | AATGATACGGCGACCACCGAGATCTACACGCGTAAGATCGTCGGCAGCGTC |
| | | gDNA, cDNA | Reverse | CAAGCAGAAGACGGCATACGAGATAGCGTAGCGTCTCGTGGGCTCGG |

**Data Notes S1. Global phosphoproteomic data.** Tyrosine phosphopetides identified through global phosphoproteomic analysis are shown. Columns list uniprot accessions, gene, peptide sequence, log2 fold change (Log2FC), and statistical significance (p-value) calculated using Welch's t-test. Phosphoproteome was compared between BRD8833 vs. DMSO, BRD8833 vs. Control(BRD6501), and Control(BRD6501) vs. DMSO.

**Data Notes S2. RNAseq data.** NGS reads were trimmed for adapters and polyA tails and aligned against the human transcriptome. Gene expression was calculated as $\log_2$ counts per million (CPM), filtering out low-expressing genes with CPM < 0.33 in at least 4 samples. Differentially expressed genes between treatment conditions were detected by moderated t-tests. The different columns list weighed p-values, FDR, log fold-change, and fold-change of each treatment compared to the vehicle (DMSO) condition. Gene expression was measured after 6 hours, 12 hours, and 24 hours of compound treatment. Cells were treated either with vehicle, asciminib, BRD8833, or control ABL binder BRD6501 in four independent replicates.

**Data Notes S3. PRISM screening data.** The different columns list the drug responses of all cell lines used in the PRISM screen and their relevant annotation according to DepMap. The AUC values correspond to drug sensitivities of different cell lines. Cells were treated with either BRD8833 (eight-point, three-fold serial dilution: 0.01 nM to 20 μM), control ABL activator BRD6501, or imatinib (eight-point, three-fold serial dilution: 0.005 nM to 10 μM) in three independent replicates.

**Data Notes S4. CRISPR-suppressor scanning.** sgRNA library and relative enrichment expressed as $\log_2$ counts per million (CPM) for each selection are listed. A total of 650 sgRNAs targeting the ABL1 gene were designed using the SpCas9 PAM sequence "NGG." Additionally, 81 control sgRNAs were included: 78 as negative controls targeting non-essential genes and three as positive controls targeting essential genes. The column "cut_site_AA" indicates the predicted amino acid residue where the sgRNA induces a cut. Protospacer counts were expressed as CPM and $\log_2$-transformed after adding a pseudo-count of one to ensure a minimum value of 1, even for sgRNAs with no detected reads. Data are presented for drug selection at 14, 21, and 63 days. A resistance score was calculated by normalizing the $\log_2$ (CPM) of drug-treated samples to the corresponding vehicle-treated controls. sgRNAs selected for validation with asciminib or BRD8833 are shown in green and orange, respectively.

**Data Notes S5. Deep sequencing of genomic DNA.** After completing 63 days of drug selection, the genomic DNA was extracted and the region around the selected sgRNA was PCR amplified to append NGS adapters and indexes analyzed by deep sequencing. Allele frequencies were determined using CRISPRessoV2. Alleles with more than 5% frequencies are shown.

**Data Notes S6. Deep sequencing of cDNA.** Since several sgRNAs target cut sites near a splice site (i.e., D504, K508), the cDNA was analyzed to determine the outcome of CRISPR-suppressor scanning on BCR-ABL. mRNA was extracted and the corresponding cDNA around the gRNA target site amplified to generate amplicons with NGS adapters and indexes. Allele frequencies were analyzed following the same procedures described for the genomic DNA. Alleles with more than 5% frequencies are shown. Most sgRNAs conferring resistance to BRD8833 result in a frameshift and premature stop codon, indicated by the asterisk in the translated product.
